# Loss of Polycomb Protein EZH2 causes major depletion of H3K27 and H3K9 tri-methylation and developmental defects in the fungus *Podospora anserina*

**DOI:** 10.1101/2020.08.21.261065

**Authors:** F Carlier, R Debuchy, L Maroc, C Souaid, D Noordermeer, P Grognet, F Malagnac

## Abstract

Selective gene silencing is key to development. The H3K27me3 enriched heterochromatin maintains transcription repression established during early development and regulates cell fate. Conversely, H3K9me3 enriched heterochromatin prevents differentiation but constitutes a permanent protection against transposable element. We exploited the fungus *Podospora anserina*, a valuable alternative to higher eukaryote models to question the biological relevance and interplay of these two distinct heterochromatin conformations. We found that H3K27me3 and H3K9me3 modifications are mutually exclusive within gene-rich regions but not within repeats. Lack of PaKmt6 EZH2-like enzyme resulted in loss of H3K27me3 and in significant H3K9me3 reduction, whereas lack of PaKmt1 SU(VAR)3-9-like enzyme caused loss of H3K9me3 only. We established that *P. anserina* developmental programs require H3K27me3 mediated silencing unlike most fungi studied to date. Our findings provide new insight into roles of these histone marks and into the relationship between chromatin modifications and development.

## Introduction

Histones are subjected to a variety of post-translational covalent modifications (Strahl and Allis 2000) that impact the overall degree of packing of the genome. In most organisms, the opened euchromatin is enriched in methylation of H3K4 (Lysine 4 of histone 3) nearby the transcriptional start sites (TSS) of the active genes and is associated with methylated H3K36 located in the body of the transcribed genes (Kouzarides 2007). By contrast, histones present in the compacted heterochromatin are enriched in either tri-methylated H3K27 (H3K27me3) or tri-methylated H3K9 (H3K9me3) (Bhaumik, Smith, and Shilatifard 2007). The compact architecture of heterochromatin limits the accessibility of the transcription machinery to the embedded DNA, thereby silencing gene expression. Repeat-rich genomic regions enriched in H3K9me3 are referred to as ‘constitutive’ heterochromatin because the subsequent silencing is constant across development (Saksouk, Simboeck, and Déjardin 2015). In contrast, the ‘facultative’ heterochromatin corresponds to the deposition of H3K27me3 on gene-rich regions, whose silencing is transient and dynamic across developmental processes, allowing cell type-specific differentiation and rapid adaptation of gene expression (Trojer and Reinberg 2007)

H3K9me3 is catalyzed by the SET-domain SU(VAR)3–9 enzymes (Martens et al. 2005; Nakayama et al. 2001; Rea et al. 2000; Rice et al. 2003) and bound by the chromodomain of the Heterochromatin Protein 1 (HP1, (James and Elgin 1986; Eissenberg et al. 1990; Singh et al. 1991). Subsequent HP1 oligomerization, which results in nucleosome binding (Canzio et al. 2011) and phase separation (Larson et al. 2017; Sanulli et al. 2019; Strom et al. 2017) further enhances chromatin compaction and spreads this structure over large genomic compartments. When present, high level of DNA cytosine methylation is found in H3K9me3 enriched regions. This modification is essential to genome stability by preventing either expression of transposable elements or illicit mitotic recombination. As demonstrated in mouse and drosophila models, when H3K9me3 enzymatic activity is absent or reduced, the embryos died because of various developmental defects (Liu et al. 2014) Moreover, since H3K9me3-dependant heterochromatin prevents the DNA binding of a large palette of transcription factors, it has long been considered as a barrier to cell fate changes (Becker, Nicetto, and Zaret 2016).

First identified in Drosophila, the conserved Polycomb group (PcG) protein complexes were shown to be both writers (Polycomb Repressive Complex 2, PRC2) and readers (Polycomb Repressive Complex 1, PRC1) of H3K27me3 (Lewis 1978; Ringrose and Paro 2004; Schuettengruber and Cavalli 2009). The catalytic subunit of the PRC2 is made of the SET domain protein enhancer of zest homolog 2 (EZH2), which tri-methylates H3K27. The H3K27me3-polycomb modification is instrumental for maintaining transcription repression established during early development and therefore has been linked to biological function related to cell fate, as different as X-inactivation (Wang et al. 2001) and hematopoiesis (Majewski et al. 2008). Conversely, when PCR2 functions are impaired, development of plants (Goodrich et al. 1997), insects (Birve et al. 2001) and mammals (O’Carroll et al. 2001a; Pasini et al. 2004) is disrupted, presumably because lineage specific programs are no longer accurate. Moreover, mutations affecting EZH2 leads to cell proliferation in cancer <u>(Canzio et al., 2011)</u>, while others are associated to congenital disorders such as the Weaver syndrome (Gibson et al. 2012) or the Ataxia-telangiectasia disease (Li et al. 2013).

In animals and flowering plants, H3K9me3 and H3K27me3 marks are quite mutually exclusive (Wiles and Selker 2017) and although these two histone modifications both result in gene expression silencing it is likely through different routes. Given the complexity and the essential nature of either of these pathways in metazoans, their respective mechanisms remain poorly understood. Even yeasts, although being otherwise powerful model systems, have some limitation in terms of chromatin remodeling studies. Indeed, if *Schizosaccharomyces pombe* possesses the Clr4 enzyme that catalyzes H3K9me3 (Rea et al. 2000), no such modification exists in *Saccharomyces cerevisiae*. As for the PRC2 complex, it is absent in all ascomycete yeasts. By contrast, filamentous fungi of the Pezizomycotina clade present canonic features for both H3K9me3- and H3K27me3-dependent heterochromatin (Shaver et al., 2010). Unlike yeasts, filamentous fungi differentiate complex multicellular structures that require mobilization of specific lineage programs. In addition, the possibility to set up very efficient and straightforward genetic approaches (i.e. construction of mutants bearing multiple deletions) was already proven to be instrumental to address complex questions such as functional redundancy and pathway interplay. These characteristics make filamentous fungi a valuable experimental alternative to higher eukaryote models.

Genetic screens performed in *Neurospora crassa*, established that H3K9me3 methyltransferase DIM-5, and HP1-homolog are both essential for developmental programs that ensure normal growth, full fertility and DNA methylation (Tamaru and Selker, 2001, Freitag et al., 2004). By contrast, in *Aspergillus nidulans*, deletions of genes encoding either the DIM-5-homolog ClrD and the HP1-homolog HepA result in enhanced expression of a subset of secondary metabolite (SM) clusters but not in developmental defect (Reyes-Dominguez et al. 2010). However in the closely related maize pest *Fusarium verticillioides, FvDIM5* deletion mutants display defects in both developmental programs (i.e. growth, conidiation, perithecium production) and overproduction of some SM clusters (Gu et al. 2017). The role of H3K27me3-dependant heterochromatin is also contrasted. In *N. crassa*, disruption of H3K27me3 methyltransferase activity did not lead to developmental defects (Jamieson et al. 2013) but to abnormal chromosome conformation (Klocko et al. 2016). By contrast, in the pathogenic fungus *Zymoseptoria tritici* absence of H3K27me3 reduces the loss of its accessory chromosomes, whereas loss of H3K9me3 induced many genomic rearrangements (Möller et al., 2019). Moreover, in *N. crassa* and *Z. tritici*, depletion of H3K9me3 resulted in massive redistribution of H3K27me3 in genomic compartments that are embedded in constitutive H3K9me3 heterochromatin in their respective wild-type backgrounds (Jamieson et al., 2016, Möller et al., 2019). In contrast, the absence of H3K27me3 machinery did not impair the H3K9me3 genomic localization in either of them. H3K27me3 loss in the plant pathogens *Fusarium graminearum* and *Fusarium fujikoroi* resulted in growth defects, sterility and in constitutive expression of genes encoding mycotoxins, pigments-producing enzymes and other SM (Connolly et al., 2013, Studt et al., 2016b). Hence, although described as being distributed in distinct genomic compartments in fungal wild-type backgrounds, both H3K9me3 and H3K27me3 can cooperate to regulate complex traits such as symbiosis (Chujo and Scott 2014a).

To further explore the functions and interplay of the constitutive versus facultative heterochromatin, we used the model system *P. anserina* to establish the genome-wide distribution of H3K4me3, H3K9me3 and H3K27me3. Surprisingly, most of the H3K9me3 are found in regions also enriched in H3K27me3, while both the H3K9me3 and H3K27me3 are co-exclusive with the H3K4me3. We demonstrated that absence of PaKmt1, the H3K9me3 methyltransferase, resulted in loss of H3K9me3, which had a limited impact on *P. anserina* growth, differentiation and sexual reproduction. Surprisingly we observed that absence of PaKmt6, the H3K27me3 methyltransferase, resulted in both H3K27me3 and H3K9me3 drastic depletion and caused severe defects in most of the aspects of its life cycle including growth, differentiation processes and sexual reproduction.

## Results

### Heterochromatin remodelers are present in *P. anserina* genome and expressed during its life cycle

*P. anserina* genome contains a single gene (*Pa_6_990*) encoding a putative H3K9me3 methyltransferase, named PaKmt1 (342 aa) and a single gene (*Pa_1_6940*) encoding a putative H3K27me3 methyltransferase, named PaKmt6 (1090 aa). Both of these highly conserved proteins show a canonical SET domain typical of the *bona fide* histone methyltransferases (Fig. 1A, Fig S1A, B and C). PaKmt1 displays a pre and post-SET domains (PS50867 and PS50868). PaKmt6 harbors a CXC domain (PS51633), which consists of two copies of a cysteine-rich motif frequently found in polycomb group (Pc-G) proteins.

**Figure 1.**
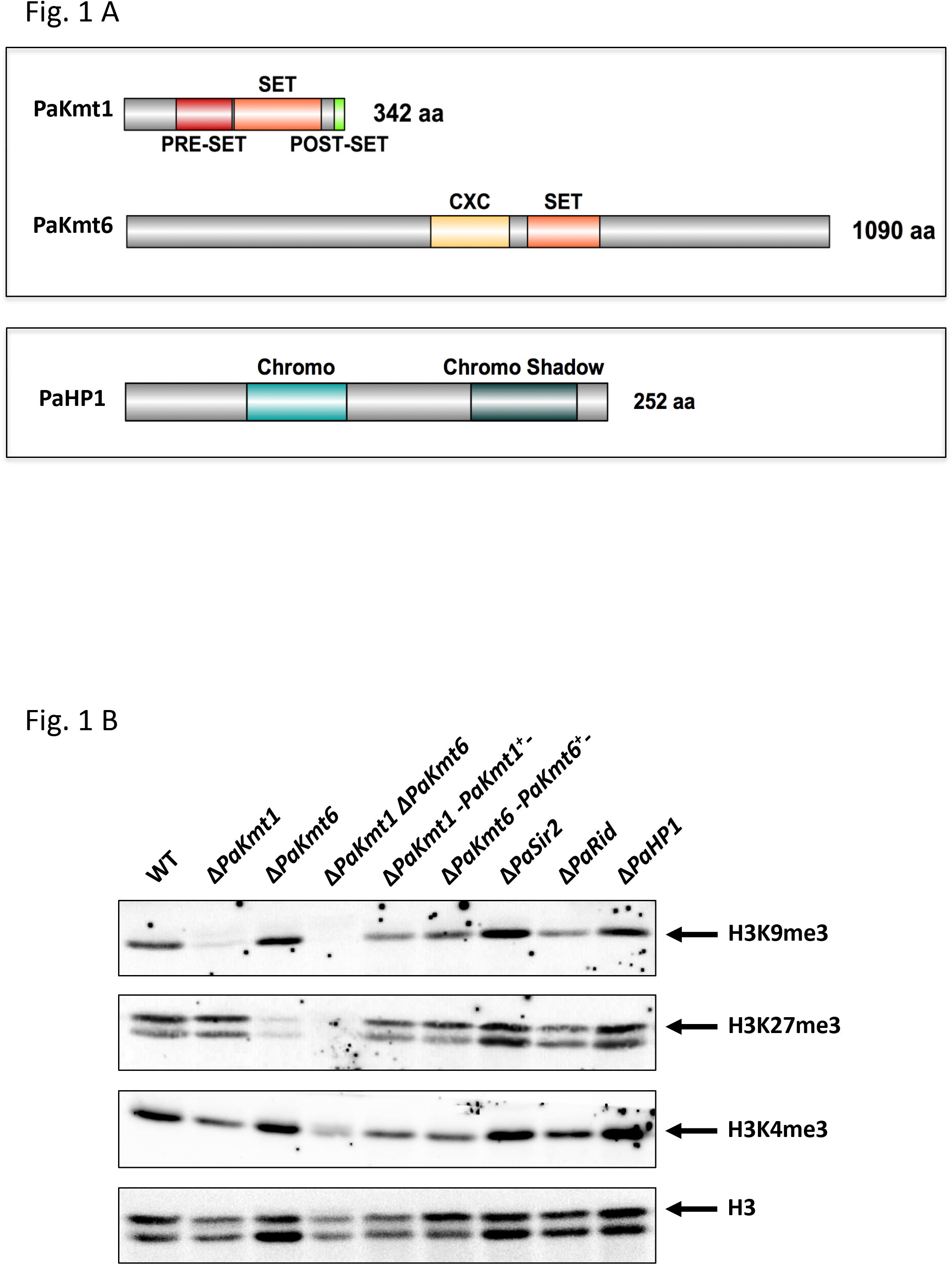
Structure and functions of the histone methyltransferases PaKmt1 and PaKmt6. **A. Domain structure of histone methyltransferases PaKmt1 and PaKmt6.** Sizes in amino acid (aa) are given (right). Pre-SET (red, IPR007728), SET (orange, IPR001214) and Post-SET (green, IPR003616) conserved domains are required for H3K9 methyltransferase activity of KMT1 homologue proteins. CXC (yellow, IPR026489) is a cysteine rich conserved domain located in the H3K27 methyltransferase catalytic domain of KMT6 homologues. The H3K9me3 reader PaHP1 is also present in *P. anserina*’s genome. It displays canonic chromo domain (PS00598) and chromo shadow domain (PS50013). **B. Detection of histone post-translational modifications** Western-immunoblot analysis of H3K9me3, H3K27me3 and H3K4me3 modifications in wild-type (WT); single mutants Δ*PaKmt1*, Δ*PaKmt6*, Δ*PaSir2* (Boivin, Gaumer, and Sainsard-Chanet 2008), Δ*PaHP1*, Δ*PaRid* (Grognet et al. 2019); double mutant Δ*PaKmt1* Δ*PaKmt6*; complemented strains Δ*PaKmt1-PaKmt1*^+^ (*PaKmt1,PaKmt1-GFP-HA*), Δ*PaKmt6-PaKmt6*^+^ (*PaKmt6,PaKmt6-HA*). Nuclear protein samples were extracted from protoplasts. A total of 35 μg of nuclear protein was loaded in each lane. Antibodies were raised against native H3 histone (H3), H3K9me3, H3K27me3 and H3K4me3 modifications. Expected size of histone 3 is 17 kDa (arrows). Probing with anti-H3K27me3 and anti-H3 antibodies reveals an additional signal, lower than 15 kDa. Deletion of either *PaSir2* or *PaRid*, encoding respectively a NAD^+^-dependent histone deacetylase and a putative DNA-methyltransferase, has no impact on methylation of H3K9me3, H3K27me3 or H3K4me3.

RNA-seq experiments detected *PaKmt6* transcripts of predicted structure in both vegetative mycelium and fruiting bodies (P. Silar et al. 2019). However, while annotation of the *PaKmt1* gene predicted that its ORF was made of two exons, RNA-seq experiments showed ambiguous profiles for *PaKmt1* transcripts (P. Silar et al. 2019). Dedicated reverse-transcription polymerase chain reaction (RT-PCR) experiments revealed two distinct transcripts (Fig. S2). From one-day-old mycelium RNA extract, we amplified a single 1108-bp fragment corresponding to the *PaKmt1* spliced transcript. However, from either four-day-old mycelium RNA extract or fruiting bodies RNA extracts (two days and four days post-fertilization fruiting bodies), we amplified an additional 1170-bp fragment, which corresponds to a *PaKmt1* unspliced transcript. If translated, the 1170-bp fragment would result in a truncated protein of 76 amino acids lacking the conserved functional motifs (pre-SET and SET domains). This observation suggests that, in addition to the functional full length PaKmt1 H3K9 methyltransferase, *P. anserina* may produce a truncated version of this protein at specific developmental stages of its life cycle.

### Genome-wide distribution of H3K4me3, H3K9me3 and H3K27me3 histone modifications in *P. anserina* genome

By using western-blotting, we first showed that H3K4me3, H3K9me3 and H3K27me3 marks were readily present in *P. anserina* genome (Fig. 1B, see WT line). Then, to further characterize the genomic patterns of these three histone modifications, we performed ChIP-seq experiments on chromatin samples collected from vegetative mycelia grown 60h at 27°C (Fig. 2A & 2B, Fig. S3, Table S1 & S2).

**Figure 2.**
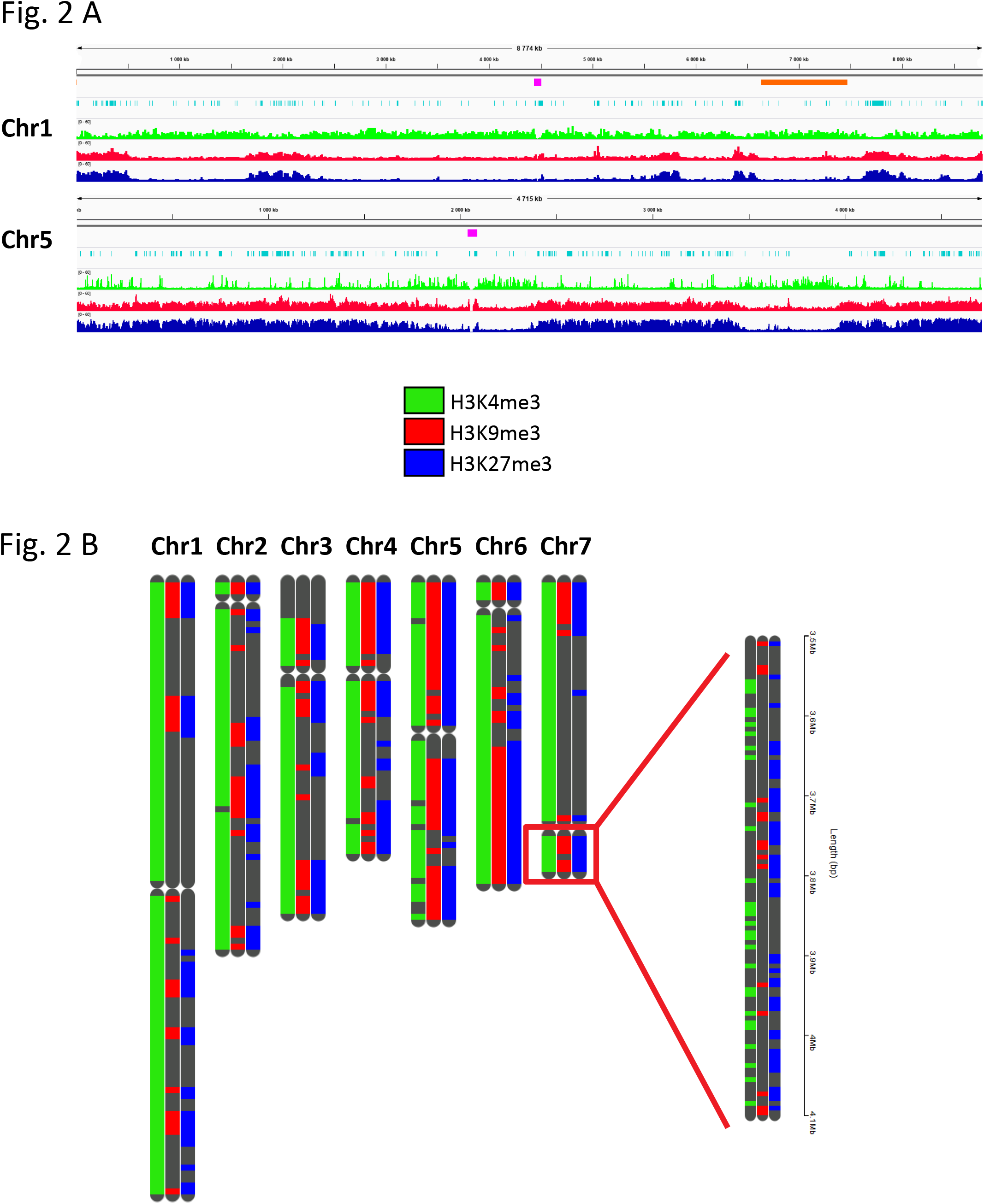
Distribution of histone marks in the wild-type strain. **A. Wild-type distribution of H3K4me3, H3K9me3 and H3K27me3 on chromosomes one and five.** Patterns are represented with the same scale on chromosomes one (top) and five (bottom) of *P. anserina*. H3K4me3 (green), H3K9me3 (red), H3K27me3 (blue). Both centromeres are depicted (pink). The “Mat region” (orange) correspond to the 800 kpb region devoid of recombination and containing the mating-type locus (Grognet, Bidard, et al. 2014). Repeated sequences are depicted in light blue. **B. Visualization of histone marks localization across the genome.** Each of the seven *P. anserina*’s chromosomes is depicted three times so each of the three marks can be mapped even if overlapping. Presence of H3K4me3, H3K9me3 and H3K27me3 is shown in green, red and blue, respectively. Centromeres are depicted as a neck on the chromosomes. The grey telomeric area of chromosome 3 represents the rDNA array. A close-up of the small arm of chromosome 7 is displayed to show the precise localization of the marks that cannot be seen on the somewhat rougher whole genome visualization.

The H3K4me3 modification covered nearly 11% of the *P. anserina* genome (Fig. S4). With 3050 detected peaks (Table S1), this mark appeared to be present in blocks of approximately 1.3 kb in gene-rich regions (Fig. 2A, Fig. 3A), mostly located in 5’ of CDS (Fig. 3B). As in other eukaryotes, H3K4me3 showed a positive correlation with active transcriptional activity (Fig. 3C). On rare instances, when H3K4me3 peaks appeared to be localized in repeats, it mainly corresponded to transposable elements (TE) relics or transposase genes inserted into CDSs (49 out of the 63 H3K4me3 peaks found in repeats).

**Figure 3.**
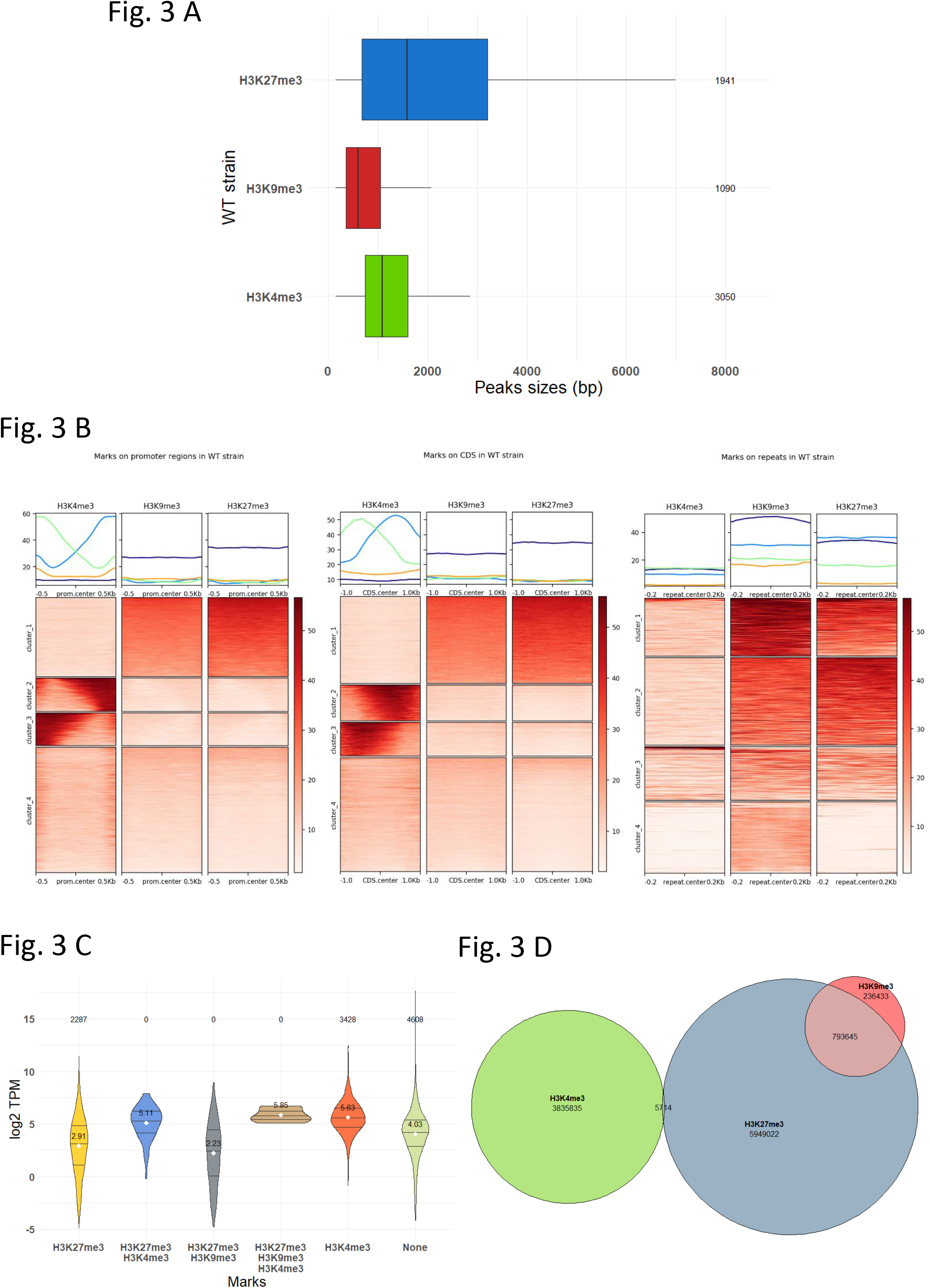
Relations of histone marks and genes in the wild-type strain. **A. Peak sizes distribution in the wild-type strain.** Box plots showing sizes (in base pair) of the MACS2 predicted peak of H3K4me3, H3K9me3 and H3K27me3 are shown in green, red and blue, respectively. The box is delimited, bottom to top, by the first quartile, the median and the third quartile. Numbers on the right of each box represents the number of peaks detected. Outliers are not shown. **B. Heatmaps of histone marks on different genomic regions in the wild-type strain.** Heatmaps and plots of normalized H3K4me3, H3K9me3 and H3K27me3 ChIP signal in wild type strain over promoters (left panel), CDS (middle) and repeats (right panel) regions. H3K4me3 clusters 2 and 3 on promoter regions are actually identical. The difference is due to the fact that genomic regions coordinates are provided regardless of their orientation. Hence, the two clusters are composed of active promoter regions in both orientation and shows that H3K4me3 is enriched close to the start codon. The same thing can be seen in a lesser extent on the middle panel. **C. Violin plots of gene expression according to the histone mark.** Gene expression was inferred from the TPM (Transcripts Per Kilobase Million) values calculated in (P. Silar et al. 2019). Log2 of the TPM values were plotted according to the marks detected of the CDS. The three lines of the violin plot represent the first quartile, the median and the third quartile. The white dot and the number above represent the mean. The numbers on top represent the number of gene in each category. **D. Euler diagram showing overlap between the different histone marks.** H3K4me3, H3K9me3 and H3K27me3 are represented by the green, red and blue areas, respectively. The numbers indicate the number of base pair covered by each mark. Overlap between areas shows the number of base pair covered by two marks.

Conversely, H3K9me3 (Fig. 2A & 2B, Fig. S3) was the less abundant mark found in *P. anserina* genome (1090 detected peaks, 2.94% of the genome (Table S1 and Fig. S4)) where it formed approximately 0.9 kb-long blocks (Fig. 3A), mainly found at repeat-rich regions, i.e. TE (728 peaks), centromeric regions and telomeric regions (Fig. 2A, Fig. 3B). Although low amount of H3K9me3 could be observed in some CDS (Fig. 3B), MACS2 peak calling analysis did not detect any significant enrichment (Table S1). In accordance with its genomic distribution, the H3K9me3 modifications showed a negative correlation with active transcriptional activity (Fig. 3C).

Being the second most common mark found in the *P. anserina* genome in terms of peak numbers (1941 detected peaks, Table S1), H3K27me3 (Fig. 2A & 2B, Fig. S3) covered none of the less the most sizeable part of it (18%, Fig. S4), showing large blocks of approximately 3.5 kb-long (Fig. 3A). This mark was found enriched at sub-telomeric regions and on a fraction of TE clusters (Fig. 2B & 3B), but also on about one third of the annotated CDSs (2992 detected peaks, Table S2). As such, most of these H3K27me3-marked CDS were likely not expressed during the vegetative growth phase (Fig. 3C).

On a broad scale, we noticed that H3K4me3, H3K9me3 and H3K27me3 modifications were not equally distributed on *P. anserina* chromosomes as shown in Fig. S4. Intriguingly, H3K27 methylation appeared enriched on chromosomes four and six and even more so on chromosome five. However, by contrast to *Zymoseptoria tritici* (Schotanus et al. 2015), the *P. anserina’s* chromosomes enriched in H3K27me3 are not dispensable. The same trend was observed for H3K9me3 on chromosome five, which is indeed enriched in TEs and repeats. Conversely, H3K4me3 appeared less abundant on chromosome five.

Zooming in *P. anserina’s* genomic patterns, it appeared that distribution of H3K4me3 was found mutually exclusive with both H3K9me3 and H3K27me3 (Fig. 2B see close up, Fig. 3B & 3D and Fig. S3). However, we did find genes that harbored both H3K4me3 and H3H27me3 marks in these experimental conditions (Table S2). Remarkably, this set was significantly enriched (p-value < 0.001) in genes encoding proteins involved in non-allelic heterokaryon incompatibility (HI), transcription and chromatin remodeling (Table S2). Notably, patterns of H3K27me3 and H3K9me3 modifications could be found overlapping in gene-poor compartments scattered across the genome (Fig. 2A and Fig. S3), especially on repeats, which included transposase promoters and CDS (Fig. 3B: cluster_1, and Table S2). Indeed, we identified 954 H3K27me3 and H3K9me3 common peaks, which represented 90% of the H3K9me3 peaks (Table S1).

### PaKmt1 and PaKmt6 are histone methyltransferases and their deletion impact both H3K9me3 and H3K27me3 distribution

To determine the impact of the lack of heterochromatin marks, we constructed the Δ*PaKmt1* and the Δ*PaKmt6* deletion mutants (Fig. S5). Double Δ*PaKmt1ΔPaKmt6* mutant strains were obtained by crossing the two corresponding single mutants. All three mutants were alive, showing that neither *PaKmt1* nor *PaKmt6* are essential genes. Western-blotting showed that H3K9me3 was almost absent in proteins extracted from the single Δ*PaKmt1* mutant (Fig. 1B). Likewise, the H3K27me3 was almost absent in extracts from the single Δ*PaKmt6* mutant. Neither mark was present in the double Δ*PaKmt1ΔPaKmt6* mutants. As expected, H3K4me3 was detected in all samples. These observations clearly show that PaKmt1 displays a H3K9me3 methyltransferase enzymatic activity, while PaKmt6 is displays a H3K27me3 methyltransferase enzymatic activity.

ChIP-seq experiments performed in the Δ*PaKmt1* background showed low amount of H3K9me3 (Fig. 4A and 4C, Fig. S3 and Fig S4), however MACS2 peak calling analysis did not detected any significant peak enrichment (Fig 4A, Table S1 and Fig. S4), by contrast to the same analysis performed in the wild-type background. This would mean that PaKmt1 is responsible for the wild-type distribution of H3K9me3 modifications at the specific locations described above. Together with the western-blot analysis, these results would further suggest that PaKmt1 is the main *P. anserina’s* H3K9me3 methyltransferase. Unexpectedly, our data revealed variations of the H3K27me3 patterns in the Δ*PaKmt1* mutant background (Fig. 4A, 4B and Fig. S4). Firstly, the average length of the H3K27me3 blocks was reduced by two third (Fig. 4C). Secondly, about 30% of the H3K27me3 peaks were missing, especially those located in the gene-rich blocks (Fig. 4A, 4B, Table S1 and Fig. S3), although chromosomes four, five and six still were enriched in H3K27me3 compared to the others (Fig. S4). Thirdly, the remaining H3K27me3 peaks did not show any extensive re-localization when compared with the H3K27me3 wild-type genome-wide distribution (Fig 4A, Fig. 4B CDS and Fig. S3). However, some H3K27me3 enrichment was observed in one cluster of repeated elements (Fig. 4B cluster_3). This observation is in stern contrast to the extensive re-localization of the H3K27me3 marks that was reported as a consequence of H3K9me3 loss in both *N. crassa* and *Z. tritici* (Jamieson et al. 2013; 2016; Möller et al. 2019).

**Figure 4.**
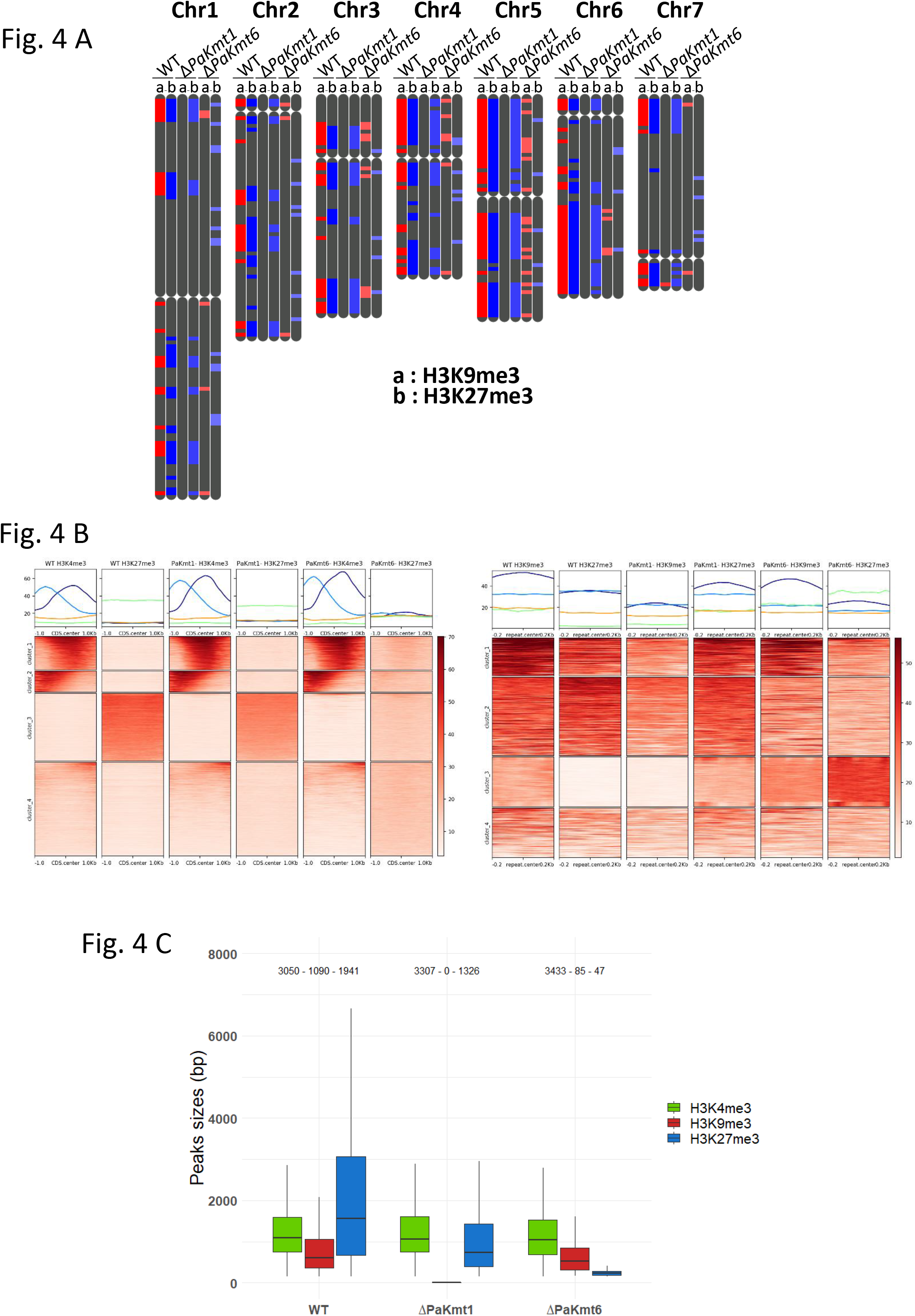
Distribution of histone marks in Δ*PaKmt6* and Δ*PaKmt1* strains. **A. Visualization of H3K9me3 and H3K27me3 localization across the genome in wild-type and mutant backgrounds.** Each of the seven *P. anserina’s* chromosomes is depicted three times corresponding to (a) the wild type strain, (b) the Δ*PaKmt1* strain and (c) the Δ*PaKmt6* strain. Centromeres are depicted as a neck on the chromosomes. The grey telomeric area of chromosome 3 represents the rDNA. A close-up of the small arm of chromosome 7 is displayed to show the precise localization of the marks that cannot be seen on the somewhat rougher whole genome visualization. **B. Heatmaps of H3K4me3 and H3K27me3 on CDS and repeats in the Δ*PaKmt1* and Δ*PaKmt6* mutant strains.** Heatmaps and plots of normalized H3K4me3 and H3K27me3 ChIP signal in mutant and wild type strains over CDS regions. **C. Peak sizes distribution in the Δ*PaKmt1* and Δ*PaKmt6* mutant strains.** Box plots showing sizes in base pair of the MACS2 predicted peak of H3K4me3, H3K9me3 and H3K27me3 are shown in green, red and blue, respectively. The box is delimited, bottom to top, by the first quartile, the median and the third quartile. Numbers on top of each box represents the number of peaks detected. Outliers are not shown.

ChIP-seq experiments performed in the Δ*PaKmt6* mutant strains revealed that this deletion drastically reduced the overall H3K27me3 (Fig. 4A, 4B & 4C and Fig. S3) content (significant genome coverage dropped from 18.95% to 0.04%, Table S1 and Fig. S4). The presence of these residual H3K27me3 marks was reminiscent of what we observed by western-blot analysis, where a faint band was still visible in the Δ*PaKmt6* mutants but disappeared in the Δ*PaKmt1* Δ*PaKmt6* double mutants (Fig. 1B). We performed a dot-blot analysis that ruled out a potential cross-hybridization, because the H3K27me3 antibody used in this study cannot detect the H3K9me3 epitopes (Fig. S6). Altogether, these findings suggest that PaKmt1 may have a residual H3K27me3 methyltransferase activity. But intriguingly, most of the remaining H3K27me3 marks were no longer localized into the H3K27me3 blocks detected in the wild-type genetic background (Fig. 4A and Fig. S3) and even found enriched in one cluster of repeated elements (Fig. 4B cluster 3). To our surprise, we also noticed that the loss of PaKmt6 severely impacted the amount of H3K9me3 marks, since in the Δ*PaKmt6* background, its genome coverage dropped from 2.94% to 0.18% (Fig. 4A, Table S1 and Fig. S4). This means that the diminution of the H3K9me3 peaks was as strong as that of the H3K27me3 in this mutant background. Nevertheless, the remaining H3K9me3 peaks in the Δ*PaKmt6* background were not displaced, nor the missing H3K9me3 blocks replaced by H3K27me3 ones (Fig. 4A).

Finally, H3K4me3 patterns stayed unchanged in all the mutant backgrounds, suggesting that loss of either H3K27me3 or H3K9me3 has no impact on the H3K4 methyltransferase activity (Fig. 4A & 4C, Table S1 and Fig. S4) and global chromatin relaxation did not interfere with genome-wide H3K4me3 patterns.

### Loss of H3K27me3 in Δ*PaKmt6* background modifies gene expression

To assay the impact of H3K27me3 or/and H3K9me3 loss on gene expression, we performed qRT-PCR experiments on vegetative mycelia grown 60 hours at 27°C, of either wild-type genotype (5 independent samples) or mutant genotypes (Δ*PaKmt1* and Δ*PaKmt6*, 5 independent samples each). We first focused on six genes that harbor H3K27me3 marks in wild-type background (Fig. 5A, Fig S7 and Table S2). In the Δ*PaKmt6* background (Fig. 5A, Table S2), all of them lost their H3K27me3 marks and were over expressed (fold change ≥ 2). This result is consistent with H3K27me3 being involved in regulation of gene expression during development programs since *Pa_1_6263*, which showed the most dramatic up-regulation effect (fold change > 25, p_value = 0.001 Fig. 5A), is supposed to be expressed during the sexual phase only (P. Silar et al. 2019). Expression of genes encoding secondary metabolites (SM) is known to be sensitive to chromatin compaction (Gacek and Strauss 2012; Connolly, Smith, and Freitag 2013; Studt, Janevska, et al. 2016; Studt, Rösler, et al. 2016). Indeed, when we assayed the expression of two such genes (*Pa_6_7270* and *Pa_6_7370*) in the Δ*PaKmt6* background, we found them both up regulated (Fig. 5A, Table S2). Notably, loss of PaKmt1 had no significant impact (fold change ≤ 2) on the expression of this set of genes, even if some of the H3K27me3 marks were lost in this mutant background (Fig. S7 and Table S2).

**Figure 5.**
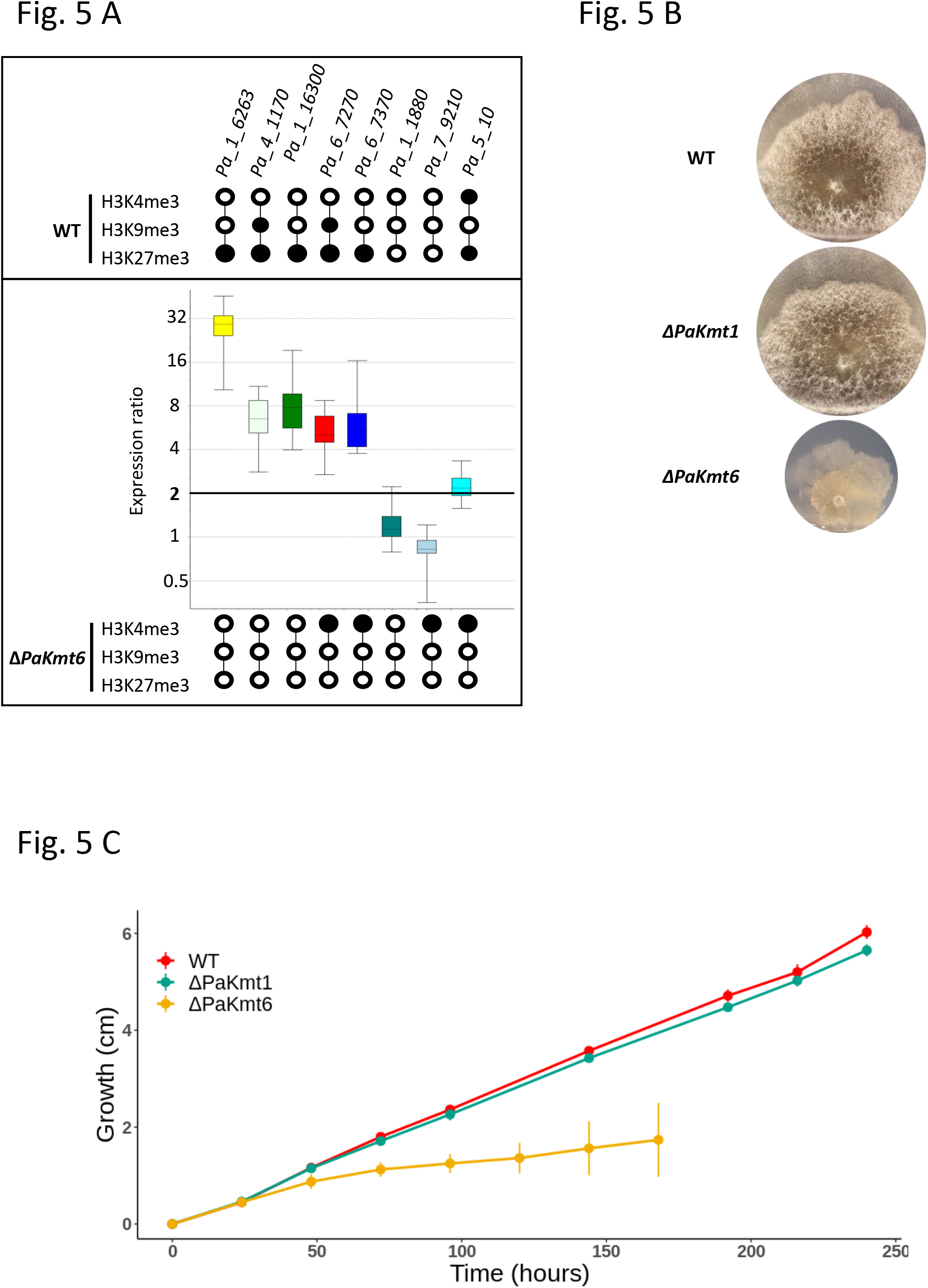
Gene expression in the Δ*PaKmt6* strain and growth features of the Δ*PaKmt6* and Δ*PaKmt1* strains. **A. Selected genes relative expression in Δ*PaKmt6* mutant strains.** Upper panel: histone marks in the wild-type strain. For each selected gene, a black dot shows the presence of the mark. Conversely a white dot shows its absence. *Pa_4_1170* and *Pa_1_16300* encode two putative Glycoside Hydrolases from family 7 and 61, respectively (Espagne et al. 2008). *Pa_6_7270* and *Pa_6_7370* belong to a secondary metabolite gene cluster located on one of chromosome 7 sub-telomeric regions. *Pa_1_1880* encodes the putative DNA repair RAD50 protein. *Pa_7_9210* encodes a putative exonuclease. They are both readily expressed at all stages of the life cycle and their expression remained unchanged in any of the two mutant backgrounds*. Pa_5_10* encodes the meiotic drive element *Spok2* located on Chromosome 5 (Grognet, Lalucque, et al. 2014b). Middle panel: normalized expression ratio of the selected genes relative to the wild-type strain. Overexpression (Fold change ≥ 2, dark black line): *Pa_1_6263:* expression ratio = 25,987,p-value = 0.001; *Pa_4_1170*: expression ratio = 6,375, p-value = 0.004; *Pa_1_16300*: expression ratio = 7,588, p-value = 0.003; *Pa_6_7270*: expression ratio = 5,280, p-value < 0001; *Pa_6_7370*: expression ratio = 6,016, p-value = 0.002; Pa_1_1880: expression ratio = 1.216, p-value = 0.192; Pa_7_9210: expression ratio = 0.756, p-value = 0.061; *Pa_5_10*: expression ratio = 2,212, p-value < 0.001. Three normalization genes (AS1, GPD and PDF2) were selected with geNorm (Vandesompele et al. 2002) among eight housekeeping genes. Average expression stability of these three normalization genes is M = 0.230 and their pairwise variation is V3/4 = 0.071. See Material and Methods for details and Table S3 for Cq and selection of normalization genes. Lower panel: histone marks in the Δ*PaKmt6* mutant strain. They are indicated as in the upper panel (see above). **B. Phenotypic characterization of Δ*PaKmt1* and Δ*PaKmt6* single mutant strains as well as Δ*PaKmt1 ΔPaKmt6* double mutant strains**. The *mat+* strains of the indicated genotypes were grown on M2 minimal medium for 5 days at 27°C (right) or M2 plus yeast extract for 3 days at 27°C (left). **C. Vegetative growth kinetics of Δ*PaKmt1* and Δ*PaKmt6* single mutant strains on M2 minimum medium.** See Material and Methods section for details.

### Loss of PaKmt6 has stronger impact on vegetative growth than loss of PaKmt1

We then assessed the impact of loss of either H3K27me3 or H3K9me3, or both, on *P. anserina’s* life cycle. Pigmentation, branching and the overall aspect of Δ*PaKmt1* mycelia were similar to those of the wild-type strains, except for reduced production of aerial hyphae (Fig. 5B). The Δ*PaKmt1* mutants also displayed a slight growth defect at optimal temperature (27°C) (6% decrease for Δ*PaKmt1*, Fig. 5C). This growth defect was stronger when the Δ*PaKmt1* mutants were grown at cold temperature (11°C, 28% decrease, Table 1). Upon cultivation on solid media, we noticed that the Δ*PaKmt1* mutants showed a delay in resuming vegetative development after a period of growth arrest (cells in stationary phase or storage at 4°C). To further explore this phenotype, we set up a stop and re-start growth assay described in (Fig. S8A). The wild-type strain was able to resume proper mycelium growth either from implants cut out of the apex of the colony (Plug#1, Fig. S8A) or out of two distinct locations of the inner stationary phase (Plug#2 & Plug#3, Fig. S8A). By contrast, Δ*PaKmt1* mycelia developing from implants cut out of the inner stationary phase of the colony (Plug#3, Fig. S8A) were growing slow and generated irregular colony margins and almost no aerial hyphae. This already defective and erratic growth was further degraded after five days of culture. However, mycelia of Δ*PaKmt1* mutants that developed from implants cut out of the apex of the colony (Plug#1, Fig. S8A) grew ordinarily. When plugs of any of these thalli where cut out of the growing edge, they all regenerated normally (Fig. S8A), confirming that this defect was not permanent but rather linked to the disability of the Δ*PaKmt1* strains to resume growth properly. Because these observations were reminiscent of those of Crippled Growth (CG), an epigenetically triggered cell degeneration (Nguyen et al. 2017), we made sure that the Δ*PaKmt1* mutation did not promote CG (Fig. S8B) but behaved as the wild-type strain in that respect. Therefore, this reduced capability to resume growth may be indicative of a lesser plasticity to perform adaptive developmental program under changing environmental conditions.

**Table 1:** Mycelium growth at different temperatures. Relative growth rates (WT %) at optimal temperature (27°C) and sub-optimal temperature (11°C and 35°C) on M2 minimal medium. Δ*PaKmt6* mutant strains display irregular and slow growth at 27°C and 35°C, which cannot be quantified. At 11°C, growth is too weak to be assayed (ND). At 27°C, double and triple mutants containing the Δ*PaKmt6* allele display aggravated Δ*PaKmt6*-like growth defects.

|  | 11°C | 27°C | 35°C |
| --- | --- | --- | --- |
| *WT* | 100 | 100 | 100 |
| *ΔPaKmt1* | 72.4 | 94.2 | 91.8 |
| *ΔPaKmt1 -PaKmt1<sup>+</sup>-* | 104 | 100 | 99.7 |
| *ΔPaKmt6* | ND | Irregular slow growth | Irregular slow growth |
| *ΔPaKmt6 -PaKmt6<sup>+</sup>-* | 98.3 | 99.0 | 99.3 |
| *ΔPaKmt1 ΔPaKmt6* | ND | *ΔPaKmt6*<br>aggravated | *ΔPaKmt6* |
| *ΔPaHP1* | 65.7 | 76.2 | 74.9 |
| *ΔPaHP1 -PaHP1<sup>+</sup>-* | 99.1 | 101 | 98.5 |
| *ΔPaKmt1 ΔPaPH1* | 60.7 | 89.9 | 80.7 |
| *ΔPaPH1 ΔPaKmt6* | ND | *ΔPaKmt6*<br>aggravated | *ΔPaKmt6* |
| *ΔPaKmt1 ΔPaPH1 ΔPaKmt6* | ND | *ΔPaKmt6*<br>aggravated | *ΔPaKmt6* |

Pinkish pigmentation (see the following paragraph), crooked branching pattern and the overall aspect of Δ*PaKmt6* mycelia were quite different to those of the wild-type strains (Fig. 5B). First of all, Δ*PaKmt6* mutant growth-rate was 43% lower than wild-type strains during the first 60 hours of growth on M2 at 27°C (Fig. 5C. Remarkably, after this time point, the mycelium development became so erratic and irregular that the standard assay consisting in thallus radius measures seemed no more accurate. Indeed, Δ*PaKmt6* mycelia showed “stop-restart” growth resulting in patchy mycelium sectors, which literally sprouted out of the colony margins like cauliflower inflorescences (Fig. 5B). Large error bars of the corresponding growth-curve represented these outgrowths (Fig. 5C). Surprisingly, we were able to grow the Δ*PaKmt6* mutants in race tubes to determine that this mutation had no effect on senescence (Fig. S9). Finally, when we constructed the Δ*PaKmt1ΔPaKmt6* double mutant, the phenotypes generated by the Δ*PaKmt6* null allele were epistatic upon those generated by the Δ*PaKmt1* allele.

### The Δ*PaKmt6* mutant produces more male gametes than the wild-type strain

In addition to the growth defect, the Δ*PaKmt6* mutant mycelium showed a powdery aspect that was different from that of the wild-type strain. When examined under the microscope, we observed a striking over-production of male gametes (spermatia) that accounted for the powdery phenotype of the Δ*PaKmt6* mutants. In this mutant context, male gametes production was about 130 times higher than in wild-type background (Fig. 6A). We then tested if these mutant male gametes were as fertile as the wild-type ones. To do so, we fertilized wild-type female gametes (ascogonia) with a defined quantity of either mutant or wild-type male gametes of compatible mating type. After seven days of growth at 27°C, fruiting bodies were numbered, given that each of them originated from a single fertilization event. Fertilization capabilities of the mutant male gametes were as efficient as the wild-type ones, since similar numbers of fruiting bodies were obtained in both conditions (4 independent experiments). This indicates that the Δ*PaKmt6* mutant male gamete over production had no impact on their biological properties.

**Figure 6.**
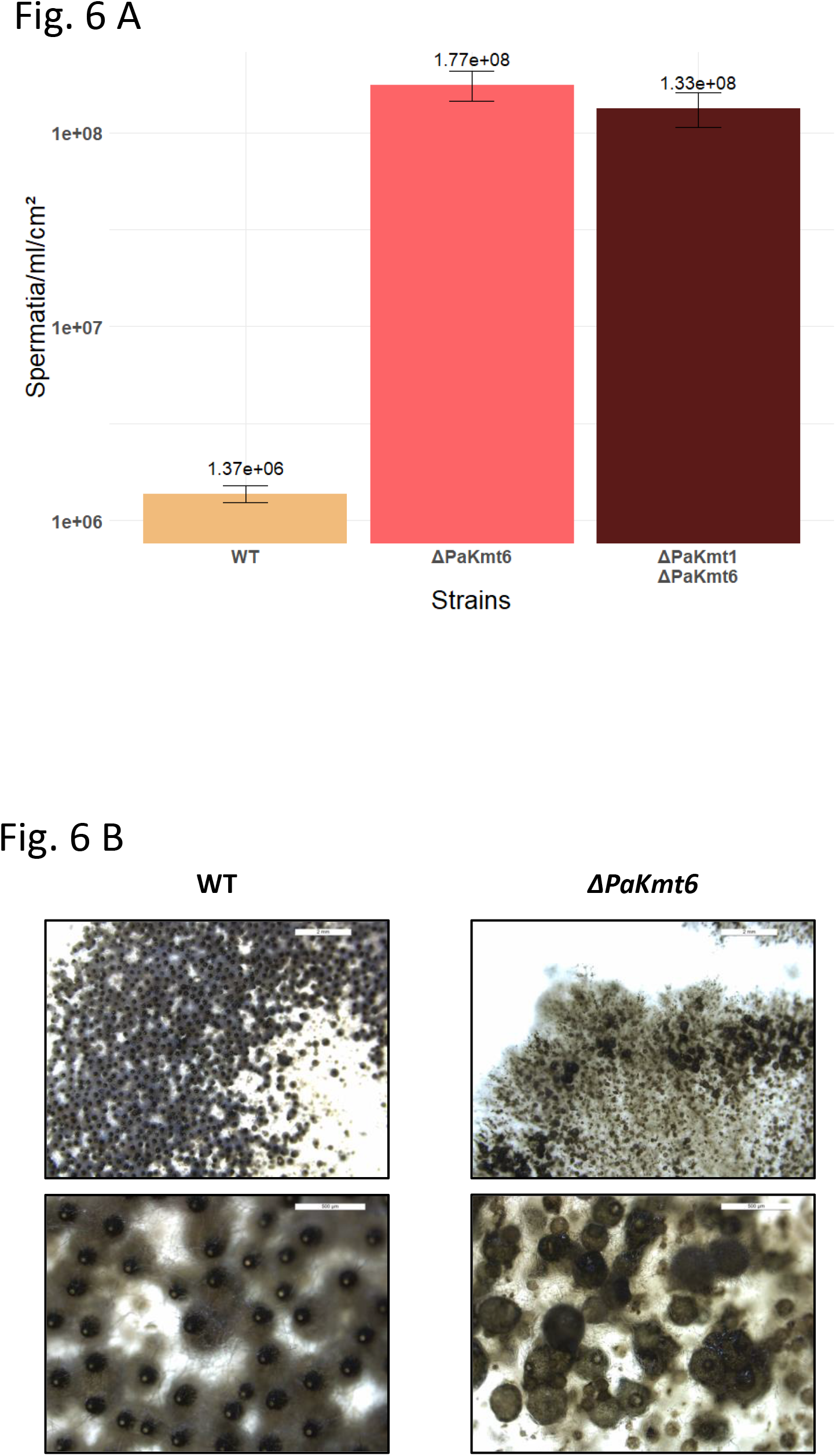
Sexual features of the Δ*PaKmt6* strain. **A. Spermatium production in wild-type and mutant backgrounds.** See Material and Methods section for details. **B. Δ*PaKmt6* homozygote crosses result in non-homogeneous distribution of late appearing fruiting bodies showing multiple morphological defects**. These crippled structures that could either be due to either lack of neck, lack of ostiols, lack setae, or non-canonical pear-shape envelopes, supernumerary necks, misdirected orientation, upside down growth in agar, fusions of fruiting bodies, etc.

### The Δ*PaKmt6* mutant formed crippled fruiting bodies yielding a reduced number of ascospores

We then investigated the ability of the histone methyltransferase mutant strains to perform sexual reproduction. Sexual development and ascospore production of crosses involving two Δ*PaKmt1* parental strains were indistinguishable from those of wild-type strains. By contrast, when we performed homozygous Δ*PaKmt6* crosses, fruiting bodies were formed, which means that fertilization occurred, but their developmental time frame was delayed. They also displayed a large palette of morphological defects *e.g*., lack of necks or extra necks, lack of ostioles, upward/downward mis-orientation, development into agar, fusion, etc. (Fig. 6B) and their ascospore production was reduced. Heterozygous orientated crosses showed that when the Δ*PaKmt6* null allele was present in the female gamete genome and the wild-type allele was present in the male gamete genome, the fruiting bodies were crippled as in the homozygous Δ*PaKmt6* crosses, which resulted in a reduced ascus production (Fig. S10A). In contrast, a heterozygous cross involving a female wild-type strain produced fully fertile fruiting bodies, indicating a Δ*PaKmt6* maternal effect.

Crosses involving the double Δ*PaKmt1* Δ*PaKmt6* mutant strains displayed a typical Δ*PaKmt6* phenotype with respect to the perithecium number, morphology and ascospore production, demonstrating that Δ*PaKmt6* null allele was epistatic. In addition, one third of the asci recovered from homozygous Δ*PaKmt1* Δ*PaKmt6* crosses showed less pigmented ascospores than those of wild-type crosses (with a color panel spanning from white to green). In general, deficit in pigmentation is indicative of incomplete maturation during the ascospore formation process.

### PaKmt6 is essential to the development of the envelope of fruiting bodies

To check whether the Δ*PaKmt6* mutant’s reduced fertility resulted from either a fruiting body envelope defect or a zygotic tissue defect, we set up tricaryon crosses involving the Δ*mat* strain (E. Coppin et al. 1993; Jamet-Vierny et al. 2007). Because the Δ*mat* strain lacks the genes required for fertilization, it do not participate either as male or female in the sexual reproduction. However, the Δ*mat* mycelium is able to provide maternal hyphae required to build fruiting bodies. Consequently, the Δ*mat* strain can complement mutants defective for fruiting body tissues formation but not for zygotic tissue defect. We observed that the Δ*mat*; Δ*PaKmt6 mat*+; Δ*PaKmt6 mat*− tricaryon yielded few but fully matured and well-formed fruiting bodies only that produced as many asci as the wild-type ones (Fig. S10B). This experiment demonstrates that Δ*PaKmt6* mutation compromised the fruiting body envelope formation but had no impact on zygotic development.

### The *PaKmt6* null allele reduces the ascospore germination efficiency

Haploid ascospores originating from homozygous Δ*PaKmt1* crosses germinated as those of wild-type crosses (> 96% germination rate, N=100, Table 2). However, about two third of the wild-type looking ascospores produced by the homozygous Δ*PaKmt6* crosses did not germinate (3 independent experiments, N=100 for each experiment, Table 2). The reduced germination efficiency was further enhanced in Δ*PaKmt1ΔPaKmt6* double mutant (Table 2). On germination medium, wild-type ascospores germination occurs at a germ pore located at the anterior side of the ascospore, by the extrusion of a spherical structure. This transient structure, called a germination peg, gives rise to a polarized hypha that develops into mycelium. In the Δ*PaKmt6* mutants, the ascospore germination process was stopped at an early stage, since germination pegs were not formed. However, the Δ*PaKmt6* defect was ascospore autonomous since ascospores from wild-type progeny obtained in WT x Δ*PaKmt6* crosses germinated normally, whereas those from Δ*PaKmt6* progeny did not. However, *PaKmt6*^+^/Δ*PaKmt6* dikaryotic ascospores had a wild-type germination rate showing that the Δ*PaKmt6* defect was recessive (2 independent experiments, N=80 dikaryotic ascospores). It has been previously shown that unpigmented ascospores carrying the *pks1-193* mutant allele germinate at high frequency (over 95%) on both inducing and non-inducing media (Malagnac et al. 2004). Indeed, the *P. anserina pks1-193* mutants carry a recessive null mutation in the gene encoding the polyketide synthase involved in the first step of DHN melanin biosynthesis (Evelyne Coppin and Silar 2007). Therefore, these mutants form unmelanized ascospores that germinate spontaneously. When crosses between *pks1-193* Δ*PaKmt6 mat*- and *pks1*^+^ Δ*PaKmt6 mat*+ strains were performed, they generated the expected asci made of two unpigmented and two pigmented ascospores. If the pigmented ascospores (*pks1*^+^ Δ*PaKmt6*) had to be transferred to inducing germination medium for some of them to germinate (app. 30%), the unpigmented ones (*pks1-193* Δ*PaKmt6*) germinated efficiently (over 90%) even on non-inducing medium. This result shows that the absence of melanin suppressed the Δ*PaKmt6* ascospore germination defect, as for those of the Δ*PaPls1* and the Δ*PaNox2* mutants (Lambou et al. 2008).

**Table 2:**
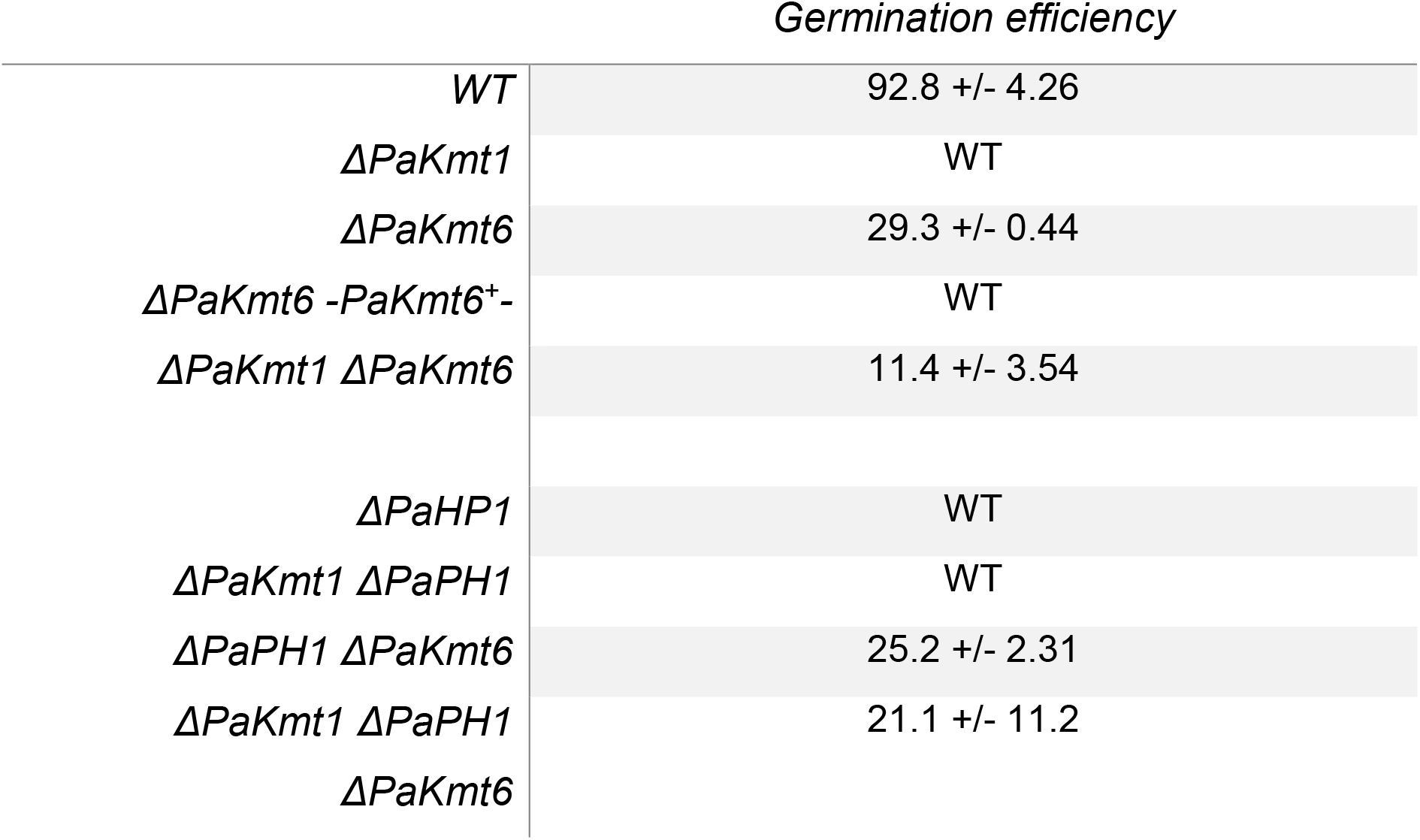
Ascospore germination efficiency. Number of germinated ascospores out of a 100 issued from homozygote Δ*PaKmt6* crosses and transferred on appropriate germination medium. WT means wild-type germination efficiency (>95 %).

### H3K9me3 reader HP1 is present in *P. anserina*

In fungi, the PRC1 complex is not present (Wiles and Selker 2017). However, we searched the *P. anserina* genome for a homolog of the heterochromatin protein 1 (HP1, (Honda and Selker 2008)) which is known to bind the H3K9me3 marks of the constitutive heterochromatin. We identified one gene (*Pa_4_7200*) encoding the protein PaHP1 (252 aa, 57% identity, e-value = 4e^-78^) (Fig. 1A). It displays conserved chromo domain (PS00598) and chromo shadow domain (PS50013) (Fig. S11A & S11B). By self-aggregation, chromo shadow domain containing proteins can bring together nucleosomes and thus condense the corresponding chromatin region (Canzio et al. 2011). As for *PaKmt6*, RNA-seq data showed that the *PaHP1* gene is expressed in both vegetative mycelium and fruiting bodies tissues (P. Silar et al. 2019).

### Loss of PaHP1 results in fewer but broader H3K9me3 domains

To explore potential role(s) of PaHP1, we deleted the corresponding *Pa_4_7200* gene (Fig. S12A & 12B). No difference in H3K4me3, H3K9me3 or H3K27me3 was detected by western-blotting performed on nuclear proteins extracted from either the wild-type strains or the single Δ*PaHP1* mutant strains (Fig. 1B). When we performed ChIP-seq experiment in the Δ*PaHP1* background, the H3K27me3 pattern was similar to that of Δ*PaKmt1* strains, while the H3K4me3 pattern was similar to that of the wild-type strains (Fig. 7A, 7B and Fig S3). However, about half of H3K9me3 genome coverage was lost (Fig. S3 and S4, Table S1), whereas some H3K9me3 enrichment was observed in one cluster of repeated elements (Fig. 7B cluster_4, Table S1). Yet, if the density of H3K9me3 was reduced, the average size of the peaks was quite larger (Fig. 7C). This suggests that, in the absence of the recruitment platform mediated by the binding of PaHP1 to the constitutive heterochromatin, the compact structuration may be partly relaxed and the histone modifications lost, but also that the boundaries between heterochromatin and euchromatin may be less clear-cut. However, in this mutant context the H3K9me3 reduction has no impact on gene expression (Fig. S13).

**Figure 7.**
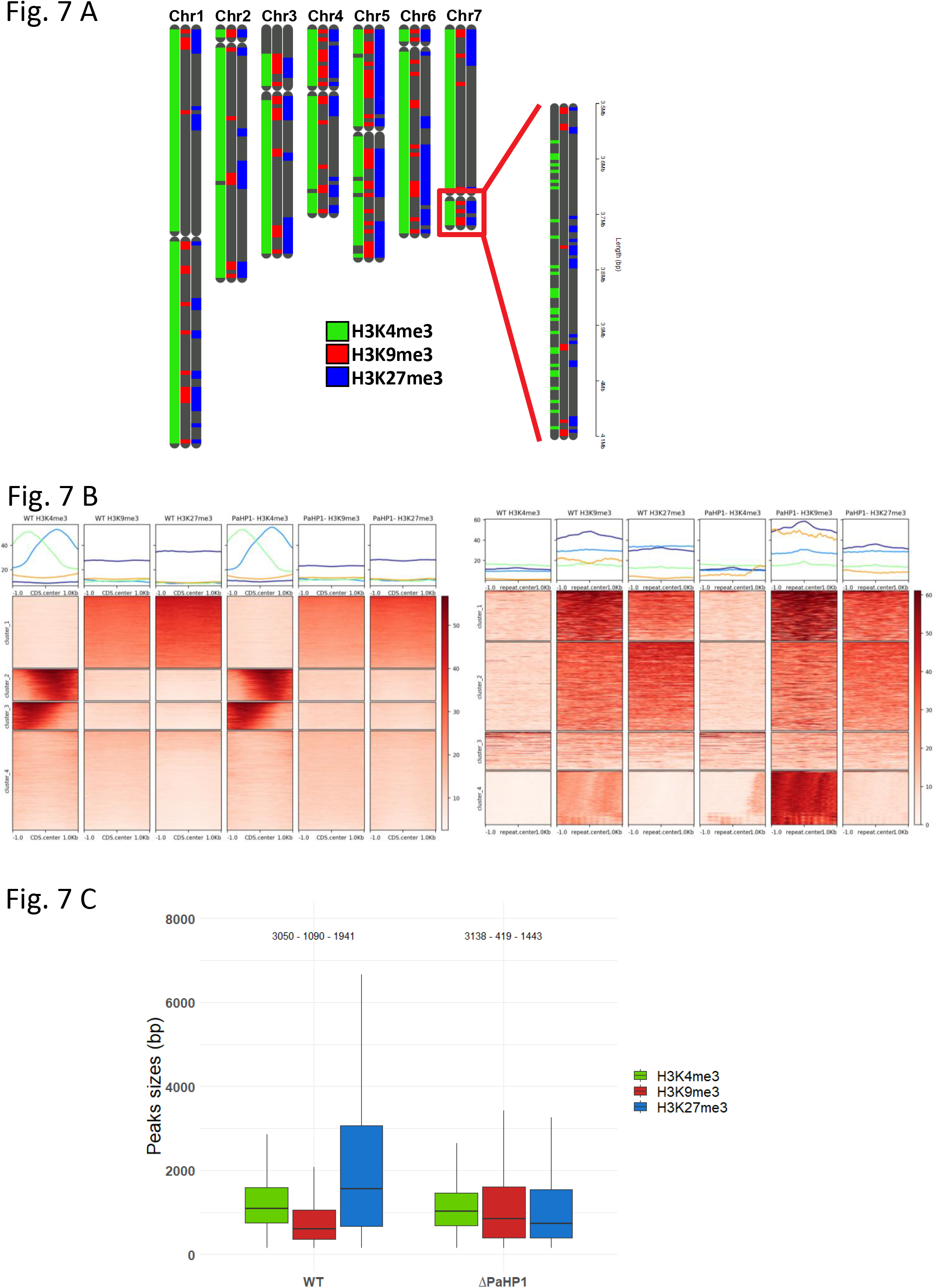
Histone marks in the Δ*PaHP1* strain. **A. Visualization of histone marks localization across the genome in the Δ*PaHP1 strain*.** Each of the seven *P. anserina’s* chromosomes is depicted three times so each of the three marks can be mapped even if overlapping. Presence of H3K4, H3K9 and H3K27 trimethylation is shown in green, red and blue, respectively. Centromeres are depicted as a neck on the chromosomes. The grey telomeric area of chromosome 3 represents the rDNA. A close-up of the small arm of chromosome 7 is displayed to show the precise localization of the marks that cannot be seen on the somewhat rougher whole genome visualization. **B. Heatmaps of H3K4me3, H3K9me3 and H3K27me3 on CDS and repeats in the Δ*PaHP1* mutant strain.** Heatmaps and plots of normalized H3K4me3, H3K9me3 and H3K27me3 ChIP signal in Δ*PaHP1* mutant and wild type strains over CDS regions (left panel) and repeat region (right panel). **C. Peak sizes distribution in the Δ*PaHP1* mutant strain.** Box plots showing sizes in base pair of the MACS2-predicted peak of H3K4, H3K9 and H3K27 trimethylation are shown in green, red and blue, respectively. The box is delimited, bottom to top, by the first quartile, the median and the third quartile. Numbers on top of each box represents the number of peaks detected. Outliers are not shown.

### Loss of PaHP1 has no more impact than that of PaKmt1

Throughout all the *P. anserina*’s life cycle, the Δ*PaHP1* mutants behave similarly to the Δ*PaKmt1* ones (Table 1, Fig. S12A, S12C, S12D, S12E). Importantly, the double Δ*PaKmt1* Δ*PaHP1* mutants did not show any aggravated phenotypes (Table 1, Fig. S13A). When the Δ*PaHP1* allele was associated to the Δ*PaKmt6* one, all the phenotypes displayed by the single Δ*PaKmt6* mutants were epistatic upon those of the Δ*PaHP1* mutants. The only exception is related to the germination efficiency of the triple Δ*PaKmt1ΔPaHP1ΔPaKmt6*, which was similar to the double Δ*PaHP1ΔPaKmt6* rather than the one of the double Δ*PaKmt1*Δ*PaKmt6* mutants (Table 2). This may be due to a slightly reduced pigmentation of the ascospores produced by the homozygous Δ*PaKmt1ΔPaHP1ΔPaKmt6* crosses. Indeed, even if only the darker wild-type looking ones were used in this test, slightly reduced pigmentation facilitates germination.

### Subcellular localization of PaKmt1-mCherry and PaHP1-GFP

Because enzymatic activity was fully restored in the Δ*PaKmt1* strain by introduction of the *PaKmt1-mCherry* allele encoding the PaKmt1-mCherry chimeric protein (See Fig. 1B, Material and Method section and below), we explored its cellular localization in two-day-old growing mycelium, along with that of PaHP1-GFP. In both PaKmt1-mCherry and PaHP1-GFP expressing strains, the fluorescence signal was nuclear and punctuated (Fig. 8). When merged, the two fluorescent signals co-localized. However, when the *PaHP1-GFP* allele was present in a Δ*PaKmt1* background, the GFP signal was present but delocalized (Fig. 8) as already observed in *N. crassa* (Freitag et al. 2004). Instead of forming foci, the GFP fluorescence was still nuclear but fuzzy, suggesting that, without H3K9me3 modification, the PaHP1 protein is no longer recruited by the appropriate heterochromatic genomic territories. On the contrary, in the same experimental conditions (two-day-old mycelium), when present in a Δ*PaKmt6*, the PaHP1-GFP protein was localized as in a wild-type background. Therefore, PaKmt6, by itself or through H3K27me3 modification, is not involved in targeting the PaHP1 to its proper genomic compartments.

**Figure 8.**
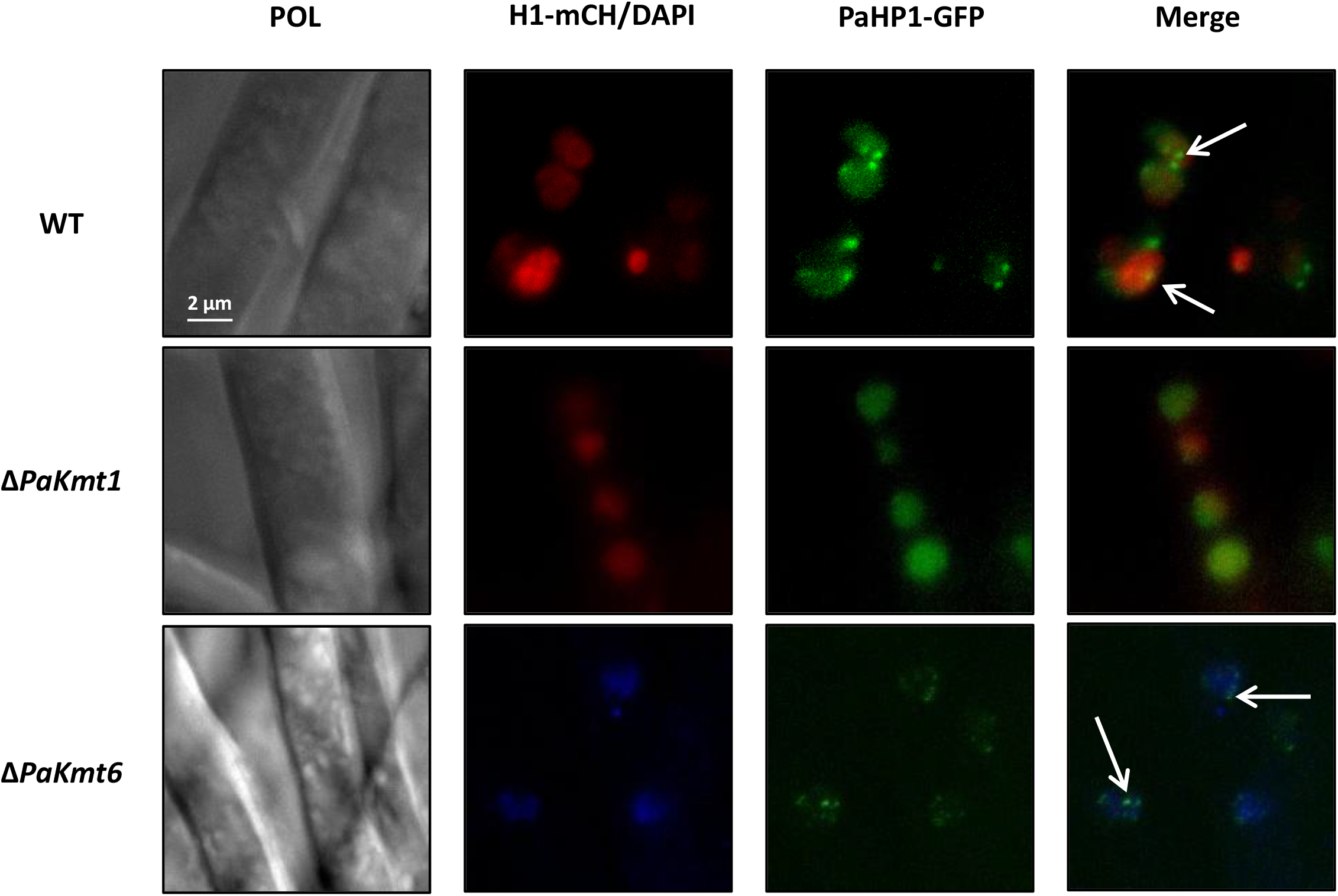
Subcellular localization of PaHP1. Fluorescence microscopy pictures show PaHP1-GFP chimeric protein during vegetative growth in mycelium. Nuclei are marked using the *PaH1-mCherry* allele that encodes histone H1 protein tagged with mCherry fluorophore (red) in the indicated strain (left) or with DAPI (1μg/μl, blue) for Δ*PaKmt6, PaHP1-GFP* strain. Indicated scale is the same for all the pictures (2 μm). PaHP1-GFP displays nuclear localization and forms foci that likely correspond to heterochromatin domains (arrows). This specific localization is disturbed only in Δ*PaKmt1* mutants.

## Discussion

In eukaryotes, the deposition of histone modification is essential to chromatin features and regulation of gene expression that impacts cell fate. Because *P. anserina* unlike *N. crassa* displays no DNA methylation but carries conserved SET-domain SU(VAR)3–9 enzyme and PRC2 complex, we aimed at understanding the impact of heterochromatin assembly on *P. anserina* life cycle in relation with its genome organization.

### H3K27me3 and H3K9me3 modifications are not mutually exclusive

To get the first hints of *P. anserina*’s genome organization, we took advantage of ChIP-seq experiments to uncover genome-wide chromatin landscapes in a wild-type background. Our results highlighted two uncommon features of both the constitutive and the facultative heterochromatin. Firstly, in most animals and plants, the H3K9me3 modification covers fairly large domains, while the H3K27me3 modification forms shorter ones. But in *P. anserina*’s genome we observed the opposite, since H3K27me3 blocks were in average three times longer than the H3K9me3 blocks. Secondly, if as expected the H3K9me3 modification was restricted to repeats-rich regions, mostly located near telomeres and centromeres (Freitag 2017), the H3K27me3 modification encompass both annotated CDS and H3K9me3 marked repeats. Co-occurrence of H3K9me3 and H3K27me3 at same genomic loci is of particular interest because these two heterochromatic marks were long considered mutually exclusive in flowering plants, mammals, and fungi (Simon and Kingston 2013; Wiles and Selker 2017). Yet recently this dogma has been challenged, since, as demonstrated by mass spectrometry experiments, H3K27me3 and H3K9me3 can coexist within the same histone octamer in embryonic stem cells (Voigt et al. 2012). Furthermore, in the ciliate *Paramecium tetraurelia*, H3K9me3 and H3K27me3 co-occur at TEs loci (Frapporti et al. 2019), as well as in the nematode *Caenorhabditis elegans* (Ho et al. 2014) and in the bryophyte *Marchantia polymorpha* (Montgomery et al. 2020). In mammals, these two marks can be found together, in association with DNA methylation, at a subset of developmentally regulated genes in mouse extra-embryonic early embryo lineages (Alder et al. 2010).

Notably, some of the human cancer cells also display both of these repressive marks (Widschwendter et al. 2007; Ohm et al. 2007; Schlesinger et al. 2007). This co-occurrence can also be found in plants, as such H3K9me3 and H3K27me3 overlap, exclusively associated with DNA methylation, had also been described at *A. thaliana’s* ETs loci (Deleris et al. 2012). The results of this study further reinforce the biological relevance of the dual H3K27/H3K9 methylation in specific heterochromatin loci and show that these superpositions of heterochromatin marks can also happen in a genomic context devoid of DNA methylation.

### Hypothetical bivalent-like genes in fungi?

As expected, the H3K4me3 modification, typically associated with active chromatin, was found in gene-rich segments of *P. anserina’s* genome. As described in most eukaryotes, this modification was shown to be mutually exclusive with both of the repressive marks H3K9me3 and H3K27me3. However in mammals, totipotent embryonic stem cells show bivalent genes marked by both active H3K4me3 and repressive H3K27me3 chromatin modification (Thalheim et al. 2017). Such bivalent genes are proposed to play a central role in developmental and lineage specification processes. Remarkably, fungal nuclei conserved their totipotency throughout their whole life cycle. Indeed, almost any hyphal fragment can give raise to complete and totipotent new individuals (Roper et al. 2011; Money 2002). The molecular basis of this peculiar feature is not yet understood. In this study, we did find genes harboring both H3K4me3 and H3H27me3 marks and moreover a sizeable part of them belonging to the *het* gene family. In *P. anserina*, the HET domain has been shown to be essential in inducing the program cell death as a consequence of HI reaction (Paoletti, Saupe, and Clavé 2007). Being members of the conserved STAND proteins (Signal-transducing ATPases with Numerous Domains), they also display NACHT and WD40 repeat domains <u>(Choi et al., 2013; Leipe et al., 2004; Paoletti et al., 2007; Saupe et al., 1995)</u>. The proteins of this family, which can oligomerize, are innate immune receptors, including NOD-like receptors in animals and NB-LRR resistance proteins in plants (Rairdan and Moffett 2006). Likewise, in fungi, STAND proteins have been proposed to act as receptors to mediate the HI allorecognition process (Paoletti, Saupe, and Clavé 2007; Paoletti and Saupe 2009). More generally, the HI may derive from a fungal innate immune system, where the STAND proteins could mediate non-self-recognition during interactions involving either individuals of the same species or of different species. By definition, the HI reaction must be quick and massive upon contacts. Therefore, expression of genes involved in this process could be finely tuned by marking them with both repressing and activating modifications.

### Chromatin status may confer distinct epigenetic identity to *P. anserina* chromosomes

On a broad scale, the patterns of histone modifications that we described in this study suggest that *P. anserina* chromosomes harbor distinct combinations of epigenomic features that may be related to functional compartmentalization of its genome.

The first combination, composed of high content of H3K4me3 and low content of both H3K27me3 and H3K9me3, is shared by chromosomes one, two, and seven. The second combination, made of reduced content of H3K4me3 and enhanced content of both H3K27me3 and H3K9me3, is that of chromosomes four and six. The last combination, which corresponds to the highest contents in the repressive H3K27me3 and H3K9me3 marks and the lowest content in the active H3K4me3 mark concerns the chromosome five only. Remarkably, this chromosome contains a TE embedded spore-killer locus, responsible for drastic distortion of meiotic segregation (Grognet, Lalucque, et al. 2014a). Chromosome three showed composite features since it grouped with chromosomes one, two and seven regarding its H3K4me3 and H3K27me3 contents but with chromosomes four and six regarding H3K9me3 content. Drop of H3K4me3 on chromosome three was due to the presence of the rDNA array on this chromosome. This non-random distribution of repressive mark cannot only be accounted for by the TE/repeat content of these chromosomes. Indeed, if the fifth chromosome is enriched in repetitive sequences, the fourth and sixth are not. These observations are reminiscent of what had been described in *Z. tritici* (Schotanus et al. 2015). This fungus is endowed with 13 core chromosomes and up to eight accessory chromosomes, the latter showing significantly higher H3K9me3 and H3K27me3 enrichments, in line with the twofold higher proportion of repetitive DNA (Möller et al. 2019). In this context, it has been proposed that the epigenetic imprint of these accessory chromosomes would signal that they could be lost, in contrast to the core chromosomes that have to be retained ((Möller et al. 2019) and Table 3). No such dispensable chromosomes are present in the *P. anserina* genome.

**Table 3:**
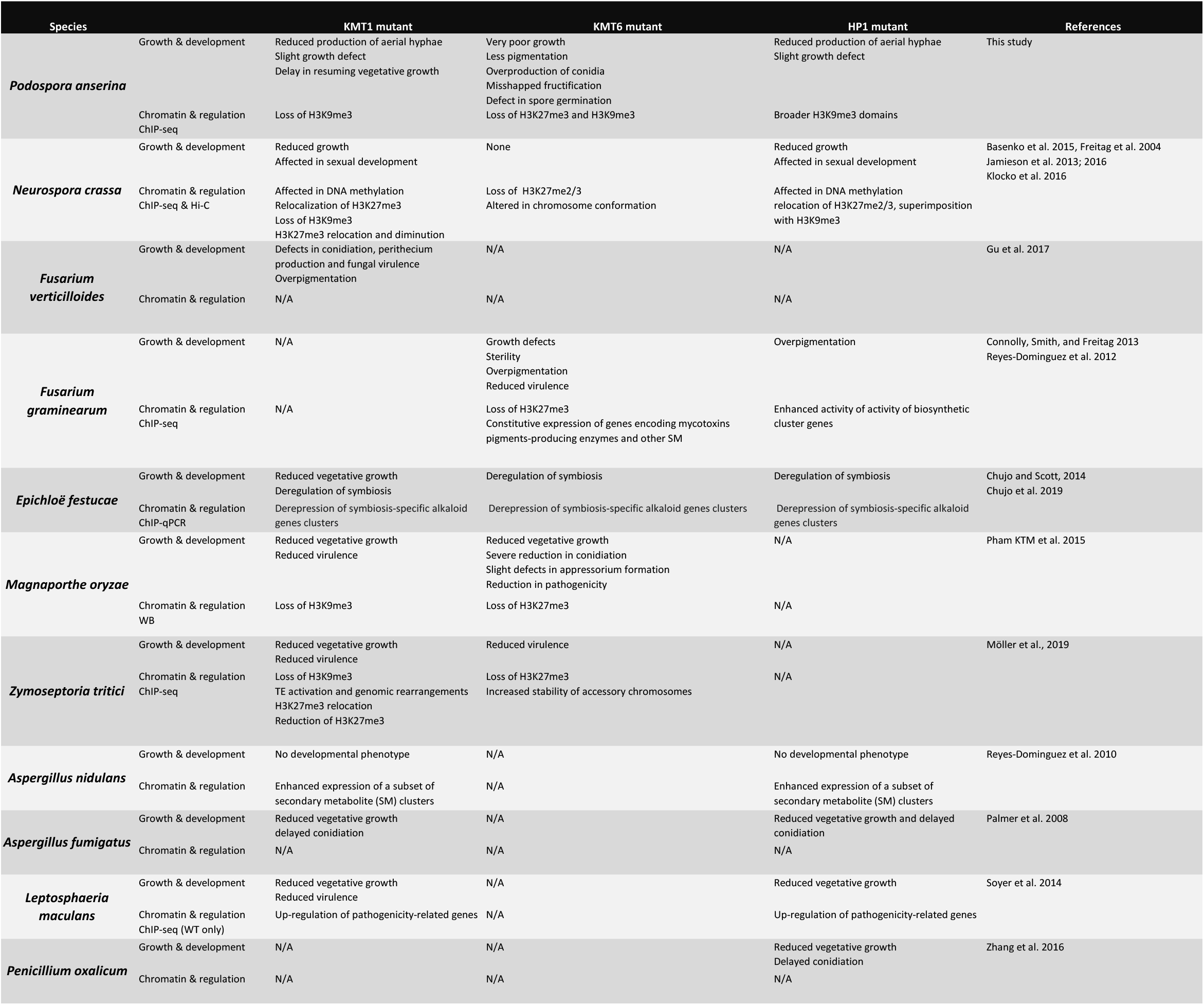
Phenotypes associated to either loss of KMT1, KMT6 or HP1 in fungal species. Growth & development: macroscopic major defects caused by the specified mutant allele. Chromatin & regulation: H3K9me3 or H3K27me3 modification status into the mutant background and major impact on regulation of gene expression. Technics used to characterize chromatin status are listed. ChiP-seq: chromatin immunoprecipitation followed by sequencing, Hi-C: genome-wide chromosome conformation capture followed by sequencing, WB: Western-blot.

### PaKmt1 and PaKmt6 are not independent actors of chromatin remodeling

The ChIP-seq carried out here can allow us to test for potential crosstalk between constitutive and facultative heterochromatin. Elimination of PaKmt1 suppressed H3K9me3, which demonstrates that there is no other H3K9 methyltransferase activity in *P. anserina*. In the absence of the PaHP1, half of the H3K9me3 peaks identified in the wild-type background were missing, indicating that some positive feedback between the H3K9me3 reader and its methylation enzyme may be needed to install or/and maintain the constitutive chromatin. Unexpectedly, our results also show that complete or partial loss of constitutive heterochromatin in Δ*PaKmt1* mutants and in Δ*PaHP1* mutants impacted facultative heterochromatin properties, since in these mutant backgrounds we identified fewer H3K27me3 peaks of reduced average size. Our data suggest that initiation and propagation of the PRC2 complex are inter-dependent processes in *P. anserina*. Alternatively, if, as shown in *A. thaliana* for the FLC locus (Angel et al. 2011), PRC2 complex initiation and propagation are independent in *P. anserina*, both of them are impaired when the H3K9me3-heterochromatin pathway is damaged. However, H3K27me3 reduction in both Δ*PaKmt1* and Δ*PaHP1* mutants was not associated with any substantial genomic redistribution by contrast to what has been described in the corresponding *N. crassa* and *Z. tritici* mutants (Jamieson et al. 2013; 2016; Möller et al. 2019) (Table3). This latter result means that although impaired in both Δ*PaKmt1* and Δ*PaHP1* mutants, the H3K27me3-heterochromatin is none of the less properly localized and does not spread over the vacant H3K9me3 genomic compartments.

Conversely, the *PaKmt6* mutants showed massive loss of H3K27me3 but also some significant reduction of H3K9me3, which could be explained if PaKmt6 targets PaKmt1 to its genomic compartment. Again theses data are in sharp contrast with those obtained in *N. crassa* (Jamieson et al. 2013; 2016) and *Z. tritici* (Möller et al. 2019) for which H3K9me3 content is not affected by loss of H3K27me3 (Table 3). Moreover, the few H3K27me3 marks that remained in the Δ*PaKmt6* mutants were found delocalized when compared to their wild-type distribution. But importantly, these new locations did not overlap with the otherwise H3K9me3-marked genomic segments.

In general, two distinct enzymes catalyze methylation of either H3K9 or H3K27 residues. However, the ciliate *P. tetraurelia* is endowed with only one class of histone methlytransferase, the Enhancer-of-zeste-like protein Ezl1, which methylates both H3K9 and H3K27 residues (Frapporti et al. 2019). This observation suggests that during the course of evolution H3K9 and H3K27 can be substrates of the same methyltransferase. In the light of the above assertion, we can hypothesize that (i) the *P. anserina’s* SU(VAR)3-like homolog (PaKmt1) may have a residual H3K27 histone-methyltransferase activity, that would account for the remaining delocalized H3K27me3 marks in the Δ*PaKmt6* mutants, (ii) SET-domain Enhancer-of-zeste-like PaKmt6 cannot be a catalytic substitute to PaKmt1, since H3K9me3 peaks were no longer detected in the Δ*PaKmt1* mutants.

### Constitutive chromatin is dispensable whereas facultative chromatin is required for most of the *P. anserina* developmental programs

Assessing the phenotype of Δ*PaKmt1, ΔPaHP1*, and Δ*PaKmt6* single mutants, we established that the loss of both PaKmt1 and PaHP1 has almost no effects on *P. anserina* life cycle, whereas the absence of PaKmt6 drastically impairs every developmental step. Hence, if briefly recapitulated, the Δ*PaKmt6* inactivation resulted in (i) slow, irregular, and thermo-sensitive mycelium growth, (ii) absence of aerial hyphae, (iii) striking over-production of male gametes, (iv) reduced fertility of the female gametes and/or of fertilization efficiency, (v) formation of crippled and misorientated fruiting bodies, and (vi) reduced yields of ascospores (vii) reduced germination efficiency of ascospores (Table 3). This means that almost every aspects of *P. anserina* life cycle are impacted by the loss of most of H3K27me3 marks and consequently of the facultative heterochromatin (see below). By contrast, the single Δ*PaKmt1* and Δ*PaHP1* mutants showed only marginal defects on aerial hyphae production and growth resuming after a period of arrest (Table 3). More importantly, the single Δ*PaKmt1* and Δ*PaHP1* and double Δ*PaKmt1* Δ*PaHP1* mutants are fully fertile. These phenotypes are the complete opposite of what had been described for those of *N. crassa*, as well as most filamentous fungi studied to date, i.e. *Aspergillus fumigatus, Epichloë festucae, Leptoshaeria maculans, Penicillium oxalicum*, *Z. tritici* (Table 3). Conversely, for *F. graminearum* and *F. fujikuroi*, Enhancer-of-zeste-like enzymes are key for their developmental integrity, as in *P. anserina* (Table 3). *F. graminearum kmt6* mutants show aberrant germination patterns, over-pigmented mycelium, restricted growth, reduced pathogenicity, over-production of several SM clusters, and sterility, while loss of Kmt1 or HP1 orthologues does not lead to any noticeable phenotypes (Connolly, Smith, and Freitag 2013). Similarly, *F. fujikuroi kmt6* mutants form stunted mycelia that produce a reduced yield of conidia (Studt, Rösler, et al. 2016).

### Conserved function of PRC2 complex in fungal development

This intriguing contrasted picture suggests that two categories of fungi may be distinguished with respect to facultative heterochromatin functions (Table 3). In *P. anserina, F. graminearum*, and *F. fujikuroi*, methylation of H3K27 by the PRC2 complex represses developmentally regulated genes, especially by establishing and maintaining long-term gene silencing memory accordingly to cell fate specification. This is in line with what has been reported for flowering plants (Goodrich et al. 1997), insects (Birve et al. 2001), and higher eukaryotes (O’Carroll et al. 2001b) where disruption of PRC2 activity leads to abnormal differentiation of embryos and abnormal cell proliferation in cancer (Conway, Healy, and Bracken 2015). In other fungi, as in *N. crassa* (Basenko et al. 2015), *E. festucae* (Chujo and Scott 2014b), *Z. tritici* (Möller et al. 2019) and in *Cryptococcus neoformans* (Dumesic et al. 2015), the role of facultative heterochromatin-mediated gene silencing is no longer required for proper development. Intriguingly, these two categories are not congruent with the fungal evolution history. It does not correlate either with DNA methylation contents. Furthermore, abnormal development of *Kmt1/HP1* mutants in a panel of fungi, including *N. crassa* (Table 3), may point toward a transfer of reversible gene silencing function from facultative to constitutive heterochromatin. Previous studies already suggested that the ancestral role of Enhancer-of-zeste enzymes, along with the SET-domain SU(VAR)3–9 enzymes may be to silence TEs, through the concerted action of both H3K27me3 and H3K9me3 (Frapporti et al. 2019; Montgomery et al. 2020; Veluchamy et al. 2015). The frequent overlap of these two histone modifications we uncovered in *P. anserina’s* chromatin could be a relic of this genome defense procedure. Under this hypothesis, specialization of the Enhancer-of-zeste proteins toward developmental and lineage-specific roles may have evolved latter on during evolution or may have been lost as in *Mucor, Rhizopus and Aspergillus* (Freitag 2017). What makes PCR2 machinery dispensable for some fungi while it is mandatory to achieve their life cycle for others? What are the evolutionary drives that selected for PCR2-dependant versus PCR2-independent regulation of gene expression? These questions remain to be further addressed. Moreover, in fungi showing a conserved function of PRC2, the PRC1 complex is not present. This implies that if H3K27me still serves as a repressive mark, alternative unknown mechanisms should replace the downstream PRC1 readers, although its H2A mono-ubiquitylation activity is not always required for silencing (Pengelly et al. 2015).

Altogether, our findings using *P. anserina* as model organism shed new light on interactions of facultative and constitutive heterochromatin in eukaryotes.

## Materials and Methods

### Strains and culture conditions

The strains used in this study derived from the “S” wild-type strain of *P. anserina* that was used for sequencing (Espagne et al. 2008; Grognet, Bidard, et al. 2014). Standard culture conditions, media and genetic methods for *P. anserina* have been described (Rizet and Engelmann 1949; Philippe Silar 2013) and the most recent protocols can be accessed at http://podospora.i2bc.paris-saclay.fr. Mycelium growth is performed on M2 minimal medium, in which carbon is supplied as dextrin and nitrogen as urea. Ascospores germinate on a specific germination medium (G medium) containing ammonium acetate. The methods used for nucleic acid extraction and manipulation have been described (Ausubel et al. 1987; Lecellier and Silar 1994). Transformations of *P. anserina* protoplasts were carried out as described previously (Brygoo and Debuchy 1985).

### Identification and deletions of the *PaKmt1, PaKmt6 and PaHP1* genes

*PaKmt1, PaKmt6 and PaHP1* genes were identified by searching the complete genome of *P. anserina* with tblastn (Altschul et al. 1990), using the *N. crassa* proteins DIM-5 (NCU04402), SET-7 (NCU07496) and HP1 (NCU04017). We identified three genes: *Pa_6_990, Pa_1_6940* and *Pa_4_7200* renamed *PaKmt1, PaKmt6 and PaHP1* respectively. To confirm gene annotation, *PaKmt1* transcripts were amplified by RT-PCR experiments performed on total RNA extracted from growing mycelium (1-day- and 4-day-old mycelium) and developing fruiting bodies (2 and 4 days post-fertilization) using primers 5s-DIM5 and 3s-DIM5 (Table S4A). Functional annotation was performed using InterProScan analysis (http://www.ebi.ac.uk/interpro/search/sequence-search), Panther Classification System (http://www.pantherdb.org/panther/), PFAM (http://pfam.xfam.org/) and Prosite (http://prosite.expasy.org/).

Deletions were performed on a Δ*mus51::bleoR* strain lacking the mus-51 subunit of the complex involved in end-joining of DNA double strand breaks as described in (Lambou et al. 2008). In this strain, DNA integration mainly proceeds through homologous recombination. Replacement of the *PaKmt1* wild-type allele with hygromycin-B resistance marker generated viable mutants carrying the Δ*PaKmt1* null allele. Replacement of the *PaKmt6* wild-type allele with nourseothricin resistance marker generated viable but severely impaired mutants. Replacement of the *PaHP1* wild-type allele with hygromycin-B resistance marker also generated viable mutants carrying the Δ*PaHP1* null allele. All deletions were verified by Southern blot (Fig. S5 and Fig S12A).

### Constructing of the fluorescent-HA-tagged chimeric proteins PaHP1-GFP-HA, PaKmt1-mCherry-HA, PaKmt6-GFP-HA

The pAKS106 and pAKS120 plasmids were described in (Grognet et al. 2019). All the cloned inserts were sequenced before transformation.

To gain insight into PaHP1 localization, we constructed a PaHP1-GFP-HA fusion chimeric protein. To this end, *PaHP1* gene was PCR amplified from the S strain genomic DNA using primers FC3-BamH1 and FC4-GFP (Table S4A). The resulting amplicon harbors 520 bp of promoter sequence followed by *PaHP1* CDS cleared from its stop codon. In parallel, *eGFP* CDS was amplified from peGFP plasmid using FC5-HP1 and FC6-HindIII primers. The two DNA segments were fused by PCR using primers FC3-BamH1 and FC6-BamH1. This chimeric DNA fragment was digested with *Bam*HI and *Hind*III and cloned into the pAKS106 plasmid linearized with the same enzymes. When transformed into Δ*PaHP1* mutant strain, the *PaHP1-GFP-HA* allele is able to restore a wild-type phenotype (growth rate and aerial mycelium), showing that the tagged version of PaHP1 protein is functional.

*PaKmt1-mCherry-HA* allele construction followed the same experimental design used for *PaHP1-GFP-HA* construction. First, *PaKmt1* was PCR amplified using Dim5mChFBamH1 and Dim5mChR and resulted in a DNA amplicon composed of 454 bp of promoter followed by the *PaKmt1* CDS. The *mCherry* CDS was amplified from pmCherry plasmid using mChDim5F and mChDim5RHindIII primers. Fragments were fused by PCR, digested and cloned in pAKS106. *Kmt1-mCherry-HA* allele was able to restore wild-type growth phenotype when transformed in Δ*PaKmt1* mutant strains, suggesting that this chimeric protein is functional.

*PaKmt6* (*Pa_1_6940*) was PCR amplified from S genomic DNA using FC73-GA1KMT6 and FC74-GA1KMT6 primers, resulting in 4232 bp DNA amplicons composed of 912 bp of promoter followed by *PaKmt6* CDS cleared from its stop codon. In parallel, *GFP* allele was amplified from peGFP plasmid using FC75-GA1GFP and FC76-GA1GFP. Both fragments were cloned together in one step in the pAKS-106 plasmid previously digested with *Not*I and *Bam*HI restriction enzymes. This cloning step was performed using Gibson Assembly kit (New England Biolabs). This yielded the chimeric *PaKmt6-GFP-HA* allele. An additional *PaKmt6-HA* allele was generated by PCR amplification from the wild-type strain DNA of *PaKmt6* CDS along with its own promoter, using FC65-NotI and FC72-BamHI primers. These amplicons were digested and cloned in pAKS106 plasmid using *Not*I and *Bam*HI restriction enzymes. When transformed into Δ*PaKmt6* mutant strain, both *PaKmt6-GFP-HA* and *PaKmt6-HA* alleles were able to restore wild-type growth and sexual development, suggesting that this chimeric version of the PaKmt6 protein is functional. However, no GFP fluorescence signal was detected in the Δ*PaKmt6, PaKmt6-GFP-HA* complemented strains.

To overexpress *PaHP1*, the endogenous promoter were replaced by the *AS4* promoter that drives high and constitutive expression throughout the life cycle (P Silar and Picard 1994). *PaHP1-GFP* allele was PCR amplified from pAKS106-*HP1-GFP* (this study) using primers FC21-BamH1 and FC6-HindIII. The resulting 1877 bp amplicons harbor the chimeric construction cleared from its stop codon and promoter. The PCR product was digested with *Bam*HI and *Hind*III and then cloned in pAKS120 previously digested with the same enzymes. The *pAS4-PaHP1-GFP-HA* resulting allele was introduced in Δ*PaHP1* strains by transformation.

*pAS4-PaKmt1-mCherry-HA* over-expressed allele construction followed the same experimental procedures as the one used for *pAS4-HP1-GFP-HA* construction. *PaKmt1-mCherry* CDS was PCR amplified using FC22-*Bam*HI and mChDim5R-*Hind*III using *pAKS106-PaKmt1-mCherry-HA* plasmid as DNA template. Amplified DNA fragments were then digested and cloned in pAKS120 plasmid using *Hind*III and *Bam*HI restriction enzymes. This allele was transformed in Δ*PaKmt1* strain.

### Complementation experiments with GFP-tagged proteins

To perform complementation experiments, *PaKmt1-mCherry* allele was transformed into the Δ*PaKmt1* strain. Because *PaKmt1-mCherry* allele was linked to a phleomycin marker, 55 phleomycin resistant transformants were recovered. Among those, five wild-type looking independent transgenic strains were selected and crossed to a wild-type strain. In the progeny, all the phleomycin and hygromycin resistant haploid strains behaved similarly to wild-type strain. Two of these independent transformants were selected for further phenotypic characterization. To ask whether a constitutively over-expressed *PaKmt1* allele could have an impact on the physiology of *P. anserina*, we introduced the *AS4-PaKmt1-mCherry* allele into the Δ*PaKmt1* strain. All the phleomycin resistant transformants looked like a wild-type strain (N=30 phleomycin resistant). Two of them were crossed to a wild-type strain. In their progeny, all the phleomycin resistant strains carrying either the *PaKmt1*^+^ allele or the Δ*PaKmt1* were indistinguishable from the wild-type strains.

Similarly, we transformed a *PaHP1-GFP-HA* allele, linked to a phleomycin marker, into a mutant Δ*PaHP1* strain. Among the 20 phleomycin resistant transformants that were recovered, five independent transformants displaying a wild-type phenotype were selected and crossed to the wild-type strain. In the progeny, all the phleomycin and hygromycin resistant haploid strains behaved similarly to the wild-type strain. Transformation of the *AS4-PaHP1-GFP-HA* allele into the Δ*PaHP1* strain generated phleomycin resistant transformants (N=19) showing intermediate to full complementation but no extra phenotypes. Two transformants presenting full complementation were crossed to a wild-type strain. In their progeny, all the phleomycin resistant strains carrying either the *PaHP1*^+^ allele or the Δ*PaHP1* behaved like the wild-type strains.

Finally, the *PaKmt6-HA* allele and the *PaKmt6-GFP-HA* allele were independently transformed into a Δ*PaKmt6* strain. For the two transformation experiments, among the phleomycin resistant transformants that were recovered (N=20 and N=32, respectively), two independent transgenic strains displaying a wild-type phenotype were selected and crossed to a wild-type strain. In the progeny of these four crosses, all the phleomycin and nourseothricin resistant haploid strains behaved similarly to wild-type strain.

### Phylogenetic analysis

Orthologous genes were identified using the MycoCosm portal (Grigoriev et al. 2014) and manually verified by reciprocal Best Hit BLAST analysis. Sequences were aligned using Muscle (http://www.ebi.ac.uk/Tools/msa/muscle/) and trimmed using Jalview (version 2.9.0b2) to remove non-informative regions (i.e. poorly aligned regions and/or gaps containing regions). Trees were constructed with PhyML 3.0 software with default parameters and 100 bootstrapped data set (Guindon et al. 2010). The trees were visualized with the iTOL server (http://itol.embl.de/).

### Phenotypic analyses

Spermatium counting was performed as follows: each strain was grown on M2 medium with two cellophane layers at 18°C for 14 days (P. Silar’s personal communication). To collect spermatia, cultures were washed with 1.5 mL of sterile water. Numeration proceeded through Malassez counting chamber.

### Cytological analysis

Light microscopy was performed on two-day-old mycelium. Explants taken from complemented strains expressing either *PaHP1-GFP-HA* or *PaKmt1-mCherry-HA* were analyzed by fluorescence microscopy. Pictures were taken with a Leica DMIRE 2 microscope coupled with a 10-MHz Cool SNAPHQ charge-coupled device camera (Roper Instruments), with a z-optical spacing of 0.5 mm. The GFP filter was the GFP-3035B from Semrock (Ex: 472nm/30, dichroïc: 495nm, Em: 520nm/35). The Metamorph software (Universal Imaging Corp.) was used to acquire z-series. Images were processed using the ImageJ software (NIH, Bethesda).

### Western blot analysis

Western blots were performed on nuclear extracts made of 5.10^7^ protoplasts resuspended in 3 mL of New Cell Lysis Buffer (Tris-HCL pH 7.5 50 mM, NaCl 150 mM, EDTA 5 mM, NP-40 0.5%, Triton 1% and protease inhibitor (Roche-04693132001). The nuclear extracts were collected after centrifugation (300 g, 5 min) and resuspended in 300 μL of Nuclei Lysis Buffer (Tris-HCl pH 8 50 mM, EDTA 10 mM, SDS 0.5% and protease inhibitor (Roche-04693132001). Nuclei were then sonicated (Diagenode bioruptor plus sonicatore, 20 min at high level). Protein contents of nuclear extracts were quantified using the Qubit Protein assay kit (Invitrogen). Then, 35 μg of nuclear extract were diluted in Laemmli sample buffer and loaded on a 10% SDS polyacrylamide gel. Membranes were probed using the following high affinity ChIP-grade monoclonal antibodies: anti-Histone 3 (Abcam ab12079), anti-H3K9me3 (Abcam ab8898), anti-H3K27me3 (Millipore 17-622) and anti-H3K4me3 (Active Motif 39159).

### Chromatin immuno-precipitation

Each ChIP-seq experiment was performed on two independent biological replicates for each genotype (*mat*+ strains only). Chromatin extractions were made from mycelium grown for 60 h at 27°C in liquid culture (KH2PO4 0.25 g/L, K2HPO4 0.3 g/L, MgSO4 0.25 g/L, urea 0.5 g/L, biotin 0.05 mg/L, thiamine 0.05 mg/L, oligo-element 0.1%, yeast extract 5 g/L and dextrin 5.5 g/L). Mycelium was filtered and quickly washed with phosphate-buffered saline (PBS), resuspended and then cross-linked in 100 mL of PBS with 1% paraformaldehyde for 15 min at 27°C. Reaction was stopped by addition of glycine (60 mM). Cross-linked mycelium was filtered, washed with cold PBS buffer and dried on Whatman paper. Dried mycelium was frozen in liquid nitrogen and ground to powder in a mortar. Aliquots of mycelium powder (200 mg) were resuspended in 1 mL of Lysis Buffer (Hepes pH 7.5 50 mM, NaCl 0.5 M, EDTA pH 8 1 mM, Triton X-100 1%, Sodium deoxycholate 0.1%, CaCl_2_ 2 mM and protease inhibitor (Roche)). Chromatin was sheared to 0.15-0.6-kb fragments using Micrococcal nuclease (NEB #M0247) for 30 min at 37°C (10000 gels units/mL). The enzymatic digestion was stopped by addition of EGTA (15 mM). After centrifugation (15000 g, 15 min, twice) of the digested samples, pelleted genomic DNA fragments were discarded. Concentration of the soluble chromatin fractions was assayed using the fluorescence system Qubit dsDNA HS assay kit (Invitrogen). Prior to immunoprecipitation, 5 μg of soluble chromatin was washed in 1.1 mL of Lysis buffer containing 30 μL of magnetic beads (Invitrogen 10001D, 4 h at 4°C). From the 1.1 mL washed soluble chromatin cleared from magnetic beads, 100 μL was kept for non-IP input control. Specific antibodies were added individually to the remaining 1 mL of the washed soluble chromatin (IP samples) for an overnight incubation at 4°C on a rotating wheel. Antibodies used in this study are anti-H3K9me3 (Abcam ab8898), anti-H3K27me3 (Millipore 17-622) and anti-H3K4me3 (Active Motif 39159). One additional mock sample was done with an anti-GFP antibody (Abcam ab290) in the wild-type strain only. Chromatin was IPed by adding 20 μL of magnetic beads, incubated for 4 h at 4°C on a rotating wheel. Beads were successively washed in 1 mL for 10 min in of the following buffers: Lysis buffer (twice), Lysis buffer plus 0.5 M NaCl, LiCl Wash Buffer (Tris-Hcl pH 8 10 mM, LiCl 250 mM, NP40 0.5%, Sodium deoxycholate 0.5%, EDTA pH 8 1mM; twice for H3K9me3 immunoprecipitation) and finally Tris-EDTA Buffer (Tris-Hcl pH 7.5 10 mM, EDTA pH 8 1 mM). Elution of IP samples was done with TES buffer (Tris-Hcl pH 8 50 mM, EDTA pH 8 10 mM with SDS 1%) at 65°C. TES buffer was also added to non-IPed input sample. Both IP and input samples were uncrosslinked at 65°C overnight and treated successively with RNAse A (Thermo Fisher Scientific) (0.2 mg/mL, 2 h at 50°C) and Proteinase K (Thermo Fisher Scientific) (0.8 mg/mL, 2 h at 50°C). DNA was then subjected to phenol-chloroform purification and ethanol precipitation. DNA pellets were resuspended in 30 μL of Tris-EDTA buffer.

### RT-qPCR experiments

Cultures for RNA extractions were performed for 60 h at 27°C in liquid culture (KH2PO4 0.25 g/L, K_2_HPO_4_ 0.3 g/L, MgSO_4_ 0.25 g/L, urea 0.5 g/L, biotin 0.05 mg/L, thiamine 0.05 mg/L, oligo-element 0.1%, yeast extract 5 g/L and dextrin 5.5 g/L). For each replicate, 100 mg of mycelium was harvested, flash-frozen in liquid nitrogen and ground with a Mikro-Dismembrator S (Sartorius, Göttingen, Germany) for one minute at 2600 rpm in a Nalgene Cryogenic vial (ref # 5000-0012, ThermoFischer Scientific, Waltham, USA) with a chromium steel grinding ball (ref # BBI-8546916, Sartorius, Göttingen, Germany). Total RNAs were extracted with the RNeasy Plant Mini Kit (ref # 74904, Qiagen, Hilden, Germany), according to the protocol described for Plants and Fungi with buffer RLT. Contaminating DNA was digested in RNA solutions with RNase-free DNase (ref # 79254, Qiagen, Hilden) and RNAs were purified once more with the RNeasy Plant Mini Kit (ref # 74904, Qiagen, Hilden, Germany), according to the protocol described for RNA cleanup. RNAs were quantified with a DeNovix DS-11 spectrophotometer (Willmington, USA), checked for correct 260/280 and 260/230 ratios, and RNA quality was checked by gel electrophoresis. Reverse transcription was performed with the SuperScript III Reverse Transcriptase (ref # 18080093, Invitrogen, Carlsbad, USA) and oligo-dT. A non-reverse transcribed (NRT) control was systematically performed on a pool of NRT controls to ensure that the Cq was above the Cq obtained from corresponding reverse transcribed RNAs. For genes without introns, NRT control was performed for each replicate sample to ensure that the Cq was above the Cq obtained from corresponding reverse transcribed RNAs. Experiments were run with five biological replicates for each strain, and each biological replicate was run in technical triplicates (see Table S3 for Cq), except NRT control which were in technical duplicates (see Table S3 for Cq). When possible, primers were designed on two consecutive exons (Table S4B). Normalization genes were selected among a set of 8 housekeeping genes using geNorm (see Table S3 for details of analysis) (Vandesompele et al. 2002). RT-qPCR normalization according to the ΔΔCt method (Pfaffl, Horgan, and Dempfle 2002), standard error and 95% confidence interval calculations, and statistical analyses were performed using REST 2009 software (Qiagen, Hilden, Germany). Genes were defined as downregulated in the mutant strain if the ratio of their transcript level in the mutant strain compared with that in the wild-type strain showed a-fold change > 0 and < 1, with a p-value <0.05. On the other hand, genes were defined as up-regulated in the mutant strain if the ratio of their transcript level in the mutant strain compared with that in the wild-type strain showed a > 1-fold change, with a p-value <0.05. Ratios with a 95 % confidence interval, including 1, were not considered significant (du Prel et al. 2009). RT-qPCR experiments were MIQE compliant (Bustin et al. 2009).

### qPCR Analysis to test ChIP efficency

Quantitative PCR experiments were performed with dsDNA fluorescent detection method using the FastStart Universal SYBR Green Master from Roche (04913850001).

IP enrichments were assayed with the following protocol: 5 μL of IPed DNA samples and non-IPed input were diluted 10 times in nuclease free water. qPCR experiments were done on 2 μL of the diluted samples using the pairs of primers (0.5 μM) FC77-Actin/FC78-Actin and FC125-580/FC126-580 (see Table S4B). For each IP sample, relative concentrations were determined as an “Input percentage” with the following equation:

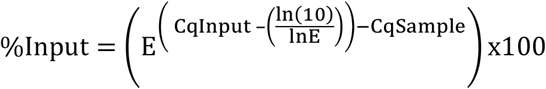

### Sequencing

ChIP-seq libraries were built using the NEB Next UltraII DNA library Prep kit for Illumina (New England Biolabs #E7645S/L) and Agencourt Ampure XP beads (Beckman Coulter, #A63880) according to the supplier protocols. PCR amplification was made of 12 cycles. ChIP-sequencing was performed by the I2BC High-throughput sequencing facility (NextSeq 500/550 High Output Kit v2 (75 cycles), SR 75 bp). Reads were trimmed with Cutadapt 1.15 and filtered for control quality by FastQC v0.11.5.

### ChIP-seq sequencing data analysis

Sequencing generated between 8 and 45 million reads depending on samples. The ChIP-seq analysis was performed on sets of data that contained 8 to 10 million reads at most. If needed, the number of reads was down-sized randomly. Reads were mapped on the S *mat+ P. anserina* genome (Espagne et al. 2008) using Bowtie 2 software (version 2.3.0, see annex for results). For visualization purpose, data were normalized using Deeptools 2.0 (Ramírez et al. 2016) bamcoverage with the BPM method. Spearman correlation factors were calculated on normalized data using Deeptools 2.0. For each condition, we made sure that the two biological replicates correlate (Fig. S14 and Table S6 and S7) and then merge them for peak calling. Peak calling was performed using MACS2 software (version 2.1.1) using the mock sample as control. Further peaks annotations and comparisons were done using Bedtools (version 2.26.0). MACS2 predicted peaks have been mapped on the genome using the ChromoMap R package (v. 0.2, https://cran.r-project.org/package=chromoMap). Genome-wide visualization of peaks was also generated with the Circos software (Krzywinski et al. 2009). Box plots have been drawn using ggplot2 (Wickham 2009). All the over plots have been generated using Deeptools2. Genomic regions have been extracted from the current genome annotation. Promoters have been arbitrarily defined as 1 kb in 5’ of the start codon. Scores of the heatmap represent the mapped reads after normalization, hence not only the MACS2 predicted peaks. The Euler diagram has been generated by the eulerr R package (v 6.0.0, https://cran.r-project.org/package=eulerr).

## Availability of data and materials’ statement

The datasets generated and/or analyzed during the current study are available in the NCBI Sequence Read Archive (SRA) (BioProject: PRJNA574032,, https://dataview.ncbi.nlm.nih.gov/object/PRJNA574032?reviewer=7beuetatcp00b0g33k3vqsno59).

## Authors’ contributions

Conceived and designed the experiments: FM. Performed the experiments: FC, VC, RD, LM, FM. Analyzed the data: FC, RD, DN, PG, FM. Contributed reagents/materials/analysis tools/expertise sharing: PG, CS, DN. Wrote the paper: FC, DN, PG, FM.

## Acknowledgments

We acknowledge the technical assistance of Sylvie François. We acknowledge the high-throughput sequencing facility of I2BC for its sequencing and bioinformatics expertise. We are thankful to Sébastien Bloyer for providing expertise to start the project as well as for fruitful scientific discussions. We thank Gaëlle Lelandais for help with statistical analysis.

---

**Supplementary Figure 1.**
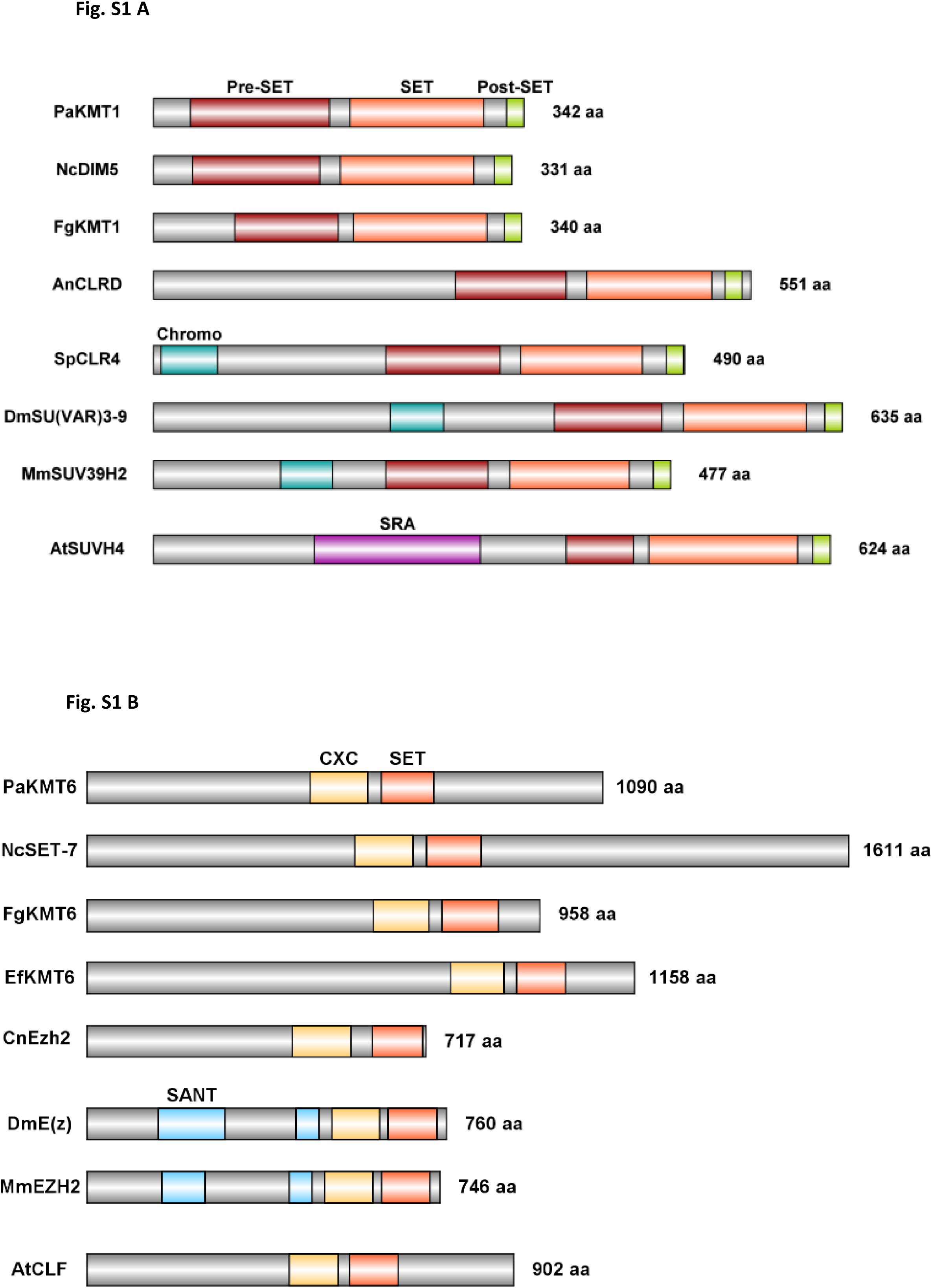
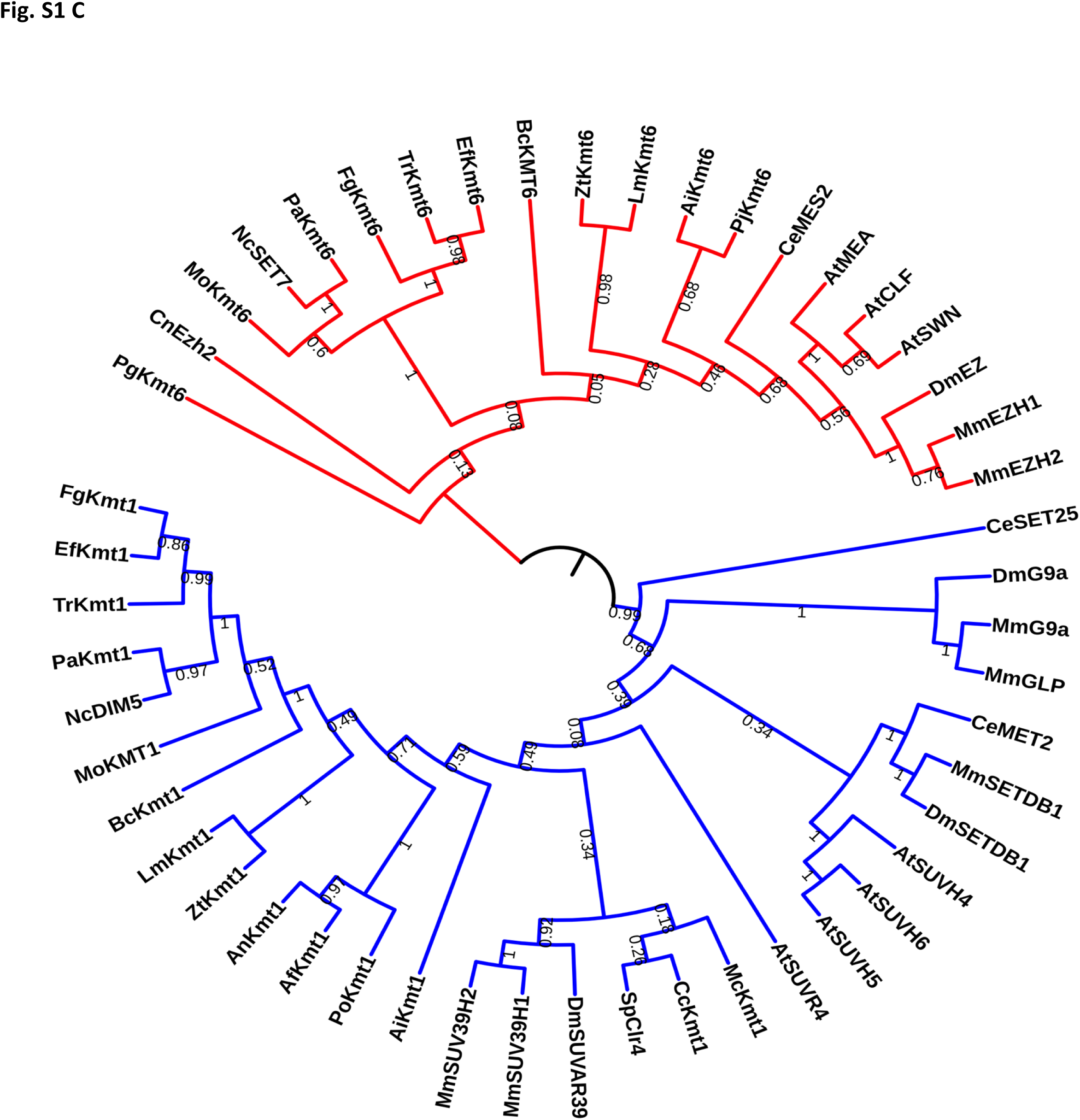
Conserved structure and phylogenetic analysis of histone methyltransferases involved in heterochromatin assembly Domain structure comparison of histone methyltransferase Kmt1 (A) and Kmt6 (B). Sizes in amino acid (aa) are given (right). Pre-SET (red, IPR007728), SET (orange, IPR001214) and Post-SET (green, IPR003616) conserved domains are required for H3K9 methyltransferase activity of KMT1 homologue proteins. SRA (SET and RING finger-associated, purple, IPR003105) and SANT (blue, IPR001005) are protein-protein interaction domains. CXC (yellow, IPR026489) is a cysteine rich conserved domain located in the H3K27 methyltransferase catalytic domain of Kmt6 homologues. Filamentous fungi: *Podospora anserina* (Pa), *Neurospora Crassa* (Nc), *Fusarium graminearum* (Fg), *Epichloë festucae* (Ef), *Aspergillus nidulans* (An); yeasts: *Schizosaccharomyces pombe* (Sp) and *Cryptococcus neoformans* (Cn); the worm *Caenorhabditis elegans* (Ce), the fruit-fly *Drosophila melanogaster* (Dm), the mouse *Mus musculus* (Mm) and the model plant *Arabidopsis thaliana* (At). Accession numbers for proteins used in alignments are listed in Table S5. **C. Phylogenetic analysis of Kmt1 and Kmt6 histone methyltransferase homologs.** This tree regroups both families of H3K9 (blue) and H3K27 (red) methyltransferase proteins from fungi, plant and metazoan. Both *P. anserinás* histone methyltransferases are marked with rectangles. They cluster two different clades of histone methyltransferase. Their evolutionary history is canonical since the topology of the two branches of the tree is consistent with the evolution of species. Bootstraps are given. Filamentous fungi: *Podospora anserina* (Pa), *Neurospora crassa* (Nc), *Fusarium graminearum* (Fg), *Trichoderma reesei* (Tr), *Epichloë festucae* (Ef) *Botrytis cinerea* (Bc), *Magnaporthe oryzae* (Mo), *Zymoseptoria tritici* (Zt), *Leptosphaeria maculans* (Lm), *Aspergillus nidulans* (An), *Aspergillus fumigatus* (Af), *Penicillium oxalicum* (Po), *Ascobolus immersus* (Ai), *Puccinia graminis* (Pg), *Pneumocystis jirovecii* (Pj), *Conidiobolus coronatus* (Cc), *Mucor circinelloides* (Mc); yeasts: *Schizosaccharomyces pombe* (Sp) and *Cryptococcus neoformans* (Cn); the worm *Caenorhabditis elegans* (Ce), the fruit-fly *Drosophila melanogaster* (Dm), the mouse *Mus musculus* (Mm) and the model plant *Arabidopsis thaliana* (At). Accession numbers for proteins used in alignments are listed in Table S5.

**Supplementary Figure 2.**
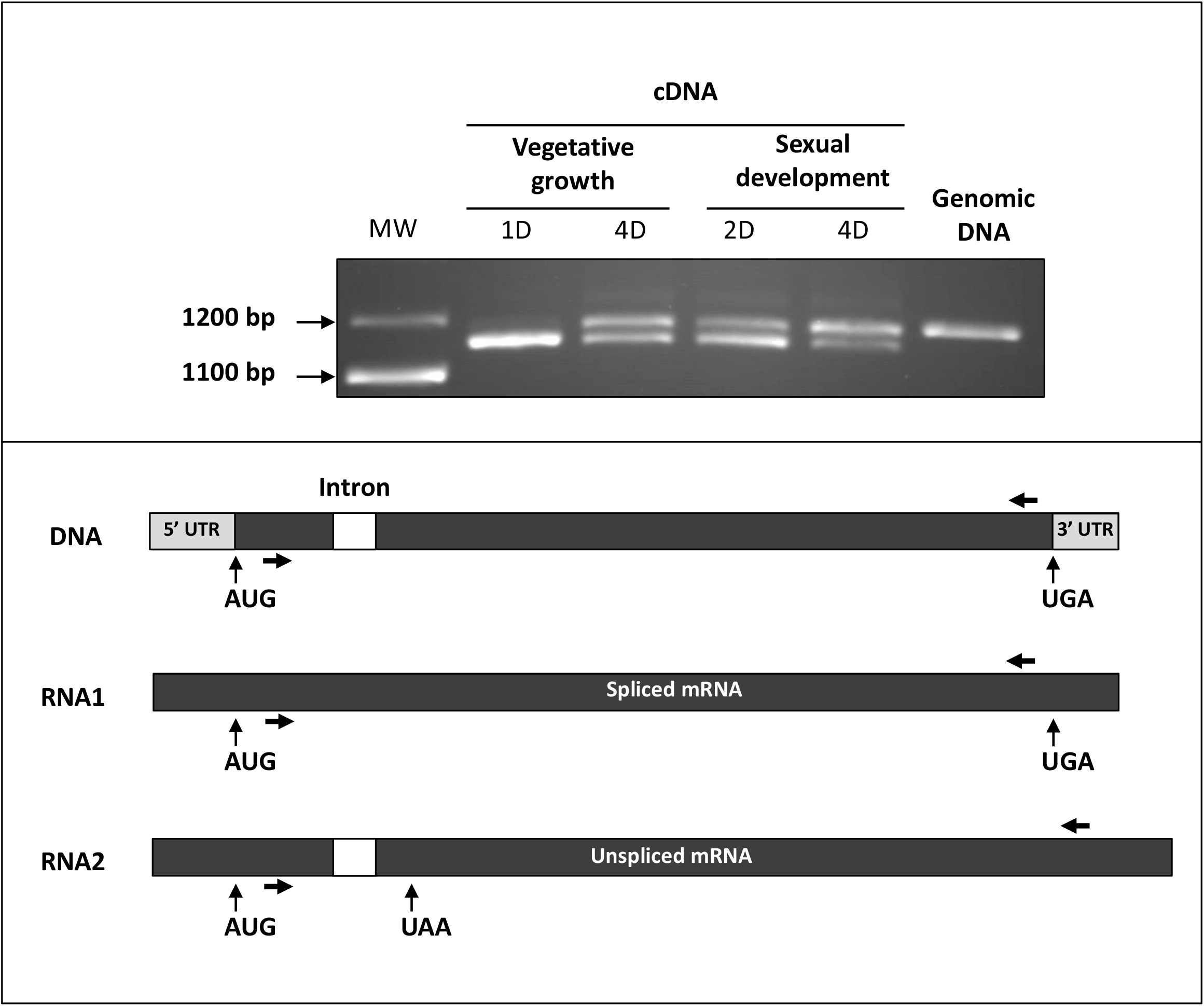
Expression kinetics *PaKmt1* in wild-type strain. Upper panel: RT-PCR on RNA extraction from wild-type vegetative growing mycelium (one day and four days), perithecia (two days and four days after the fertilization), and input genomic DNA. *PaKmt1* CDS was predicted to be made of two exons separated by a 62 bp intron (positions 49-110). However, two amplicons of distinct sizes are obtained, corresponding to both spliced (1108 bp) and unspliced (1170 bp) *PaKmt1* mRNAs. MW: molecular weight. Lower panel: schematic representation of the *PaKmt1* locus (DNA). mRNA1 corresponds to the spliced form of the transcripts while mRNA2 (1108 bp) corresponds to the unspliced form (1170 bp). Translation of the unspliced form would lead to a premature termination and thus to a truncated protein. Primers used for the reverse-transcription polymerase chain reaction (RT-PCR) are drawn as arrows above and below the *PaKmt1* CDS.

**Supplementary Figure 3.**
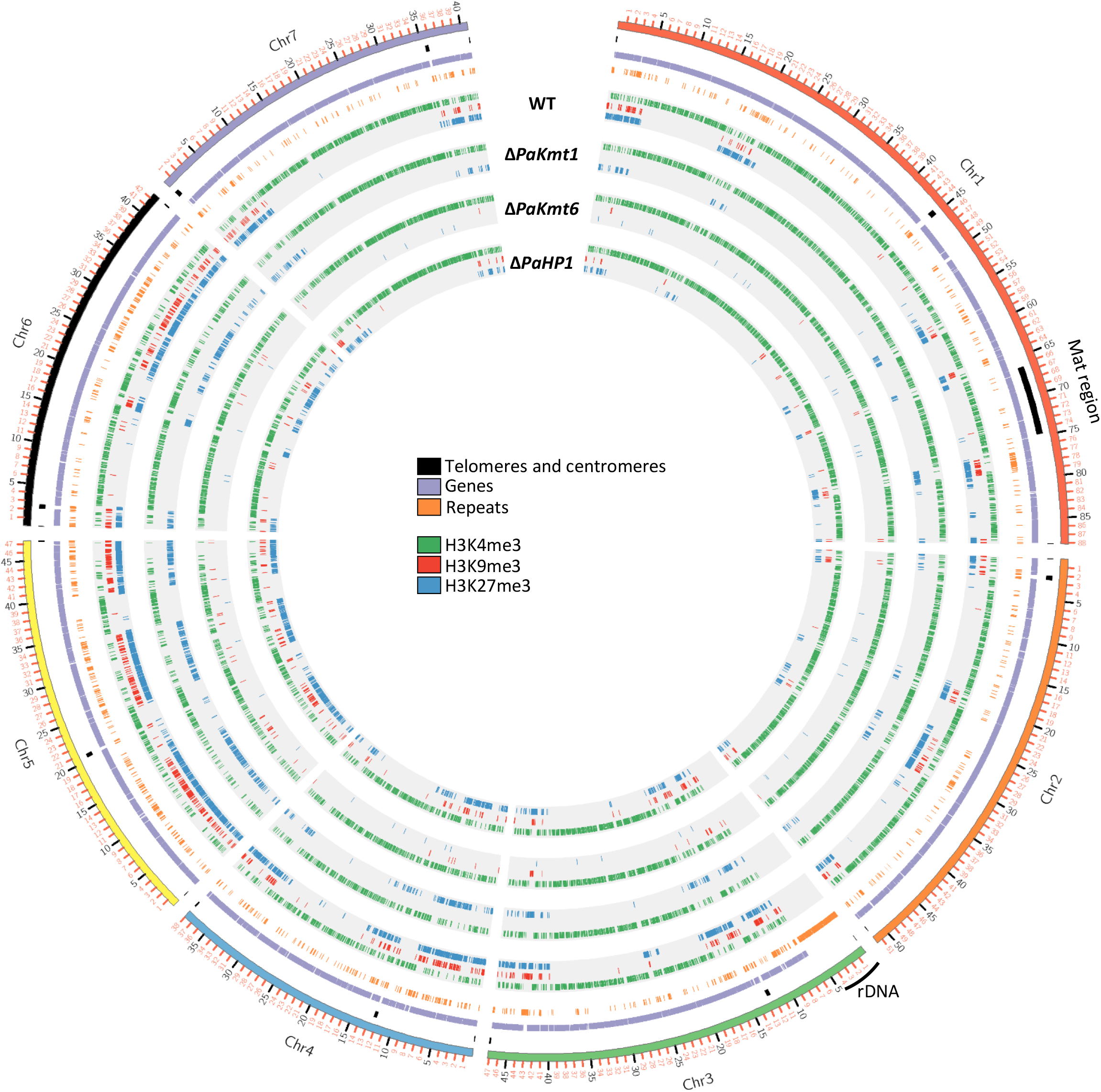
Epigenetic landscapes in wild-type and heterochromatin mutant strains of *P. anserina*. Panorama of genome-wide peak localization for each genotype, wild-type, Δ*PaKmt1*, Δ*PaKmt6* and Δ*PaHP1* strains. Telomeres sequences were arbitrarily defined as the segment going from the end of each arm of the chromosomes to the first annotated gene (at the exception of the rDNA cluster localized on chromosome 3) and centromeres are indicated. Mat region = Non recombining region containing the mating-type locus as defined in (Grognet, Bidard, et al. 2014).

**Supplementary Figure 4.**
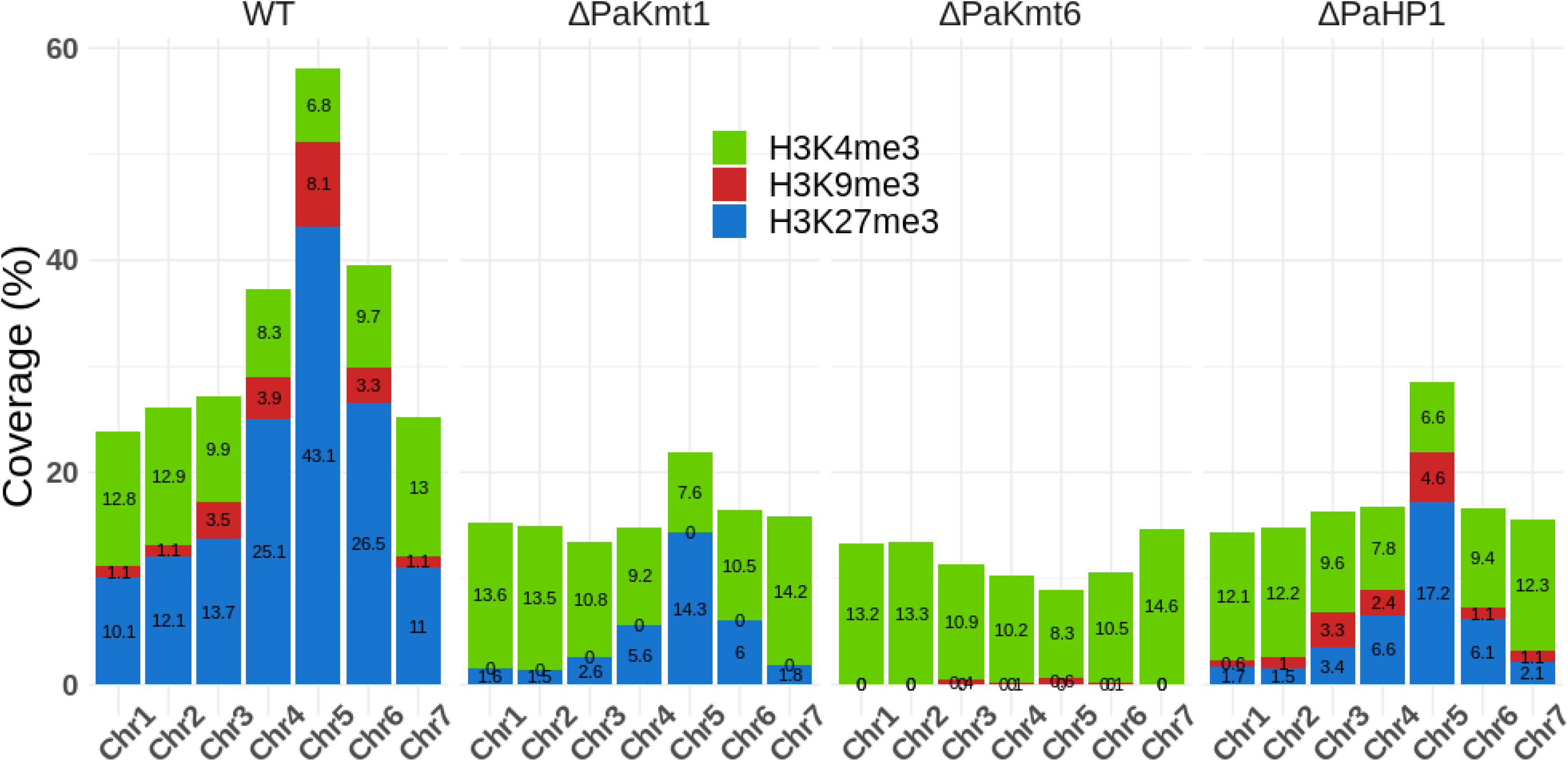
H3K27me3, H3K4me3 and H3K9me3 proportions on the *P. anserina*’s chromosomes in the WT, Δ*PaKmt1*, Δ*PaKmt6* and Δ*PaHP1* mutant strains. Plot showing the percentage of each chromosome covered with H3K4me3 (green), H3K9me3 (red) and H3K27me3 (blue). The coverage is the sum of all MACS2 predicted peak size.

**Supplementary Figure 5.**
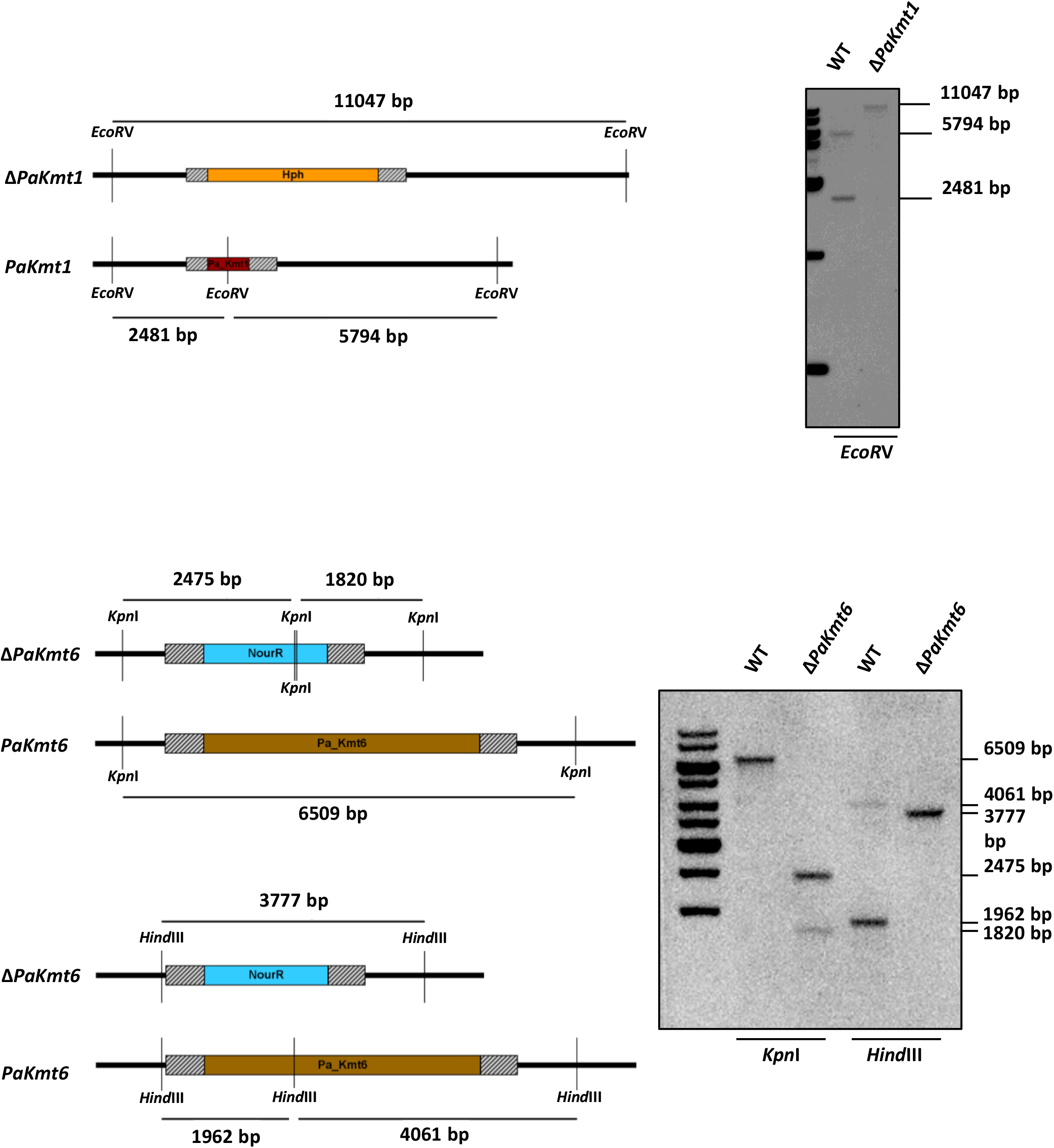
Molecular characterization of knockout mutants by Southern blot hybridization. Schematic representations of the endogenous and disrupted loci are given (Left). Replacement by homologous recombination of the wild type *PaKmt1* allele by the disrupted Δ*PaKmt1* allele results in the substitution of a 2.4 kb and 5.7 kb *Eco*RV fragments by a unique 11 kb *PstI* fragment as revealed by hybridization of the 5’ and 3’ digoxygenin-labeled probes (dashed rectangles *PaKmt1* locus). Replacement by homologous recombination of the wild type *PaKmt6* allele by the disrupted Δ*PaKmt6* allele results in the substitution of a unique 6.5 kb *KpnI* fragment by two 1.8 and 2.4 kb *KpnI* fragments as revealed by hybridization of the 5’ and 3’ digoxygenin-labeled probes (dashed rectangles *PaKmt6* locus). A second verification has been made with the *HindIII* enzymes and the same probes shows the substitution of two 1.9 kb and 4.6 kb *HindIII* fragments by a 3.7 kb *HindIII* fragment as revealed by hybridization of the 5’ and 3’ digoxygenin-labeled probes.

**Supplementary Figure 6.**
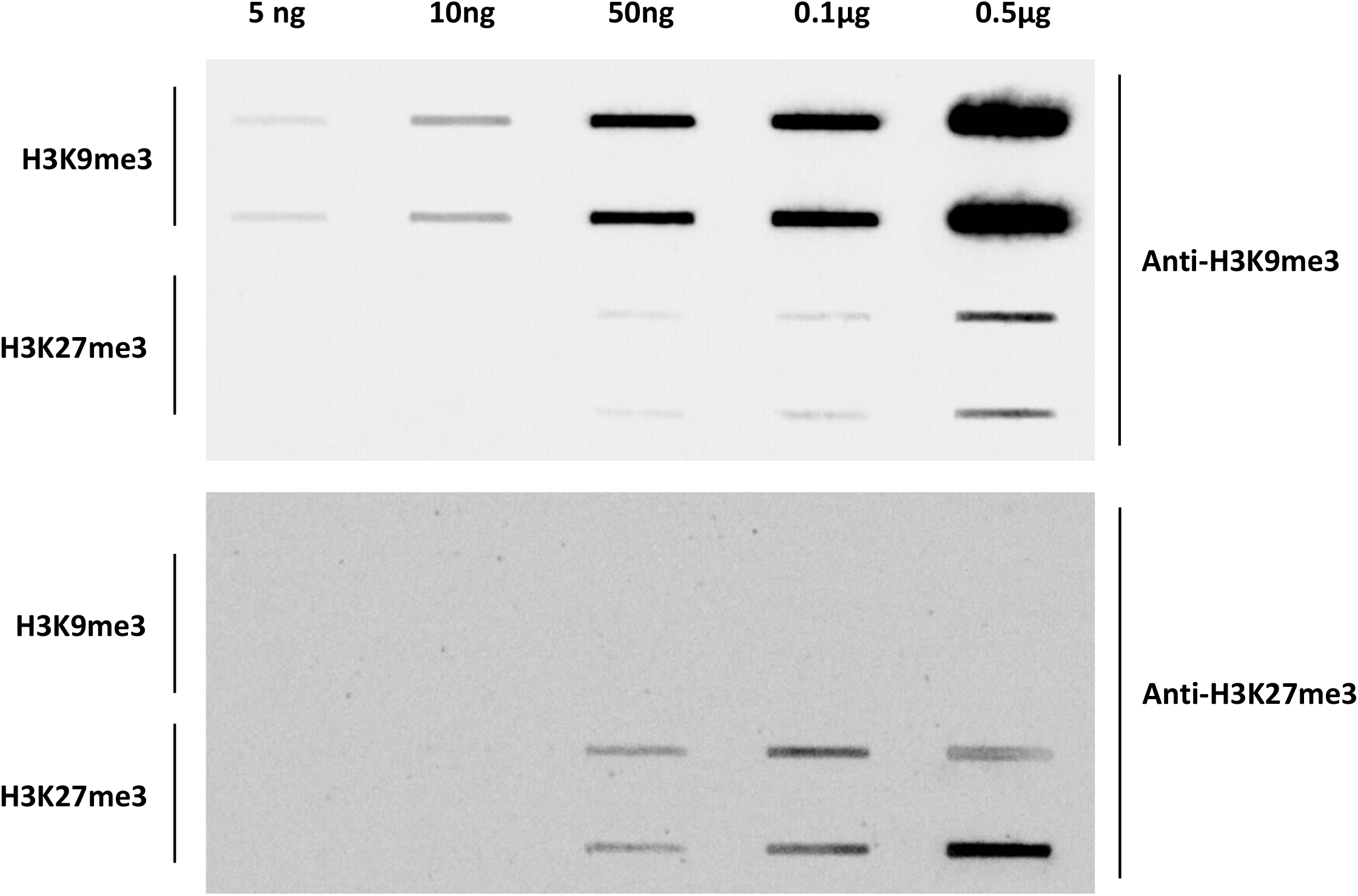
Antibody specificity analysis. Dot blot experiment results using H3K9me3 and H3K27me3 peptides (left) at different concentration (top). Membranes were inoculated with the corresponding antibody (right). Signal intensity comparison shows a cross reactivity between anti-H3K9me3 antibody and H3K27me3 evaluated at 3% compared to the immunogen reaction.

**Supplementary Figure 7.**
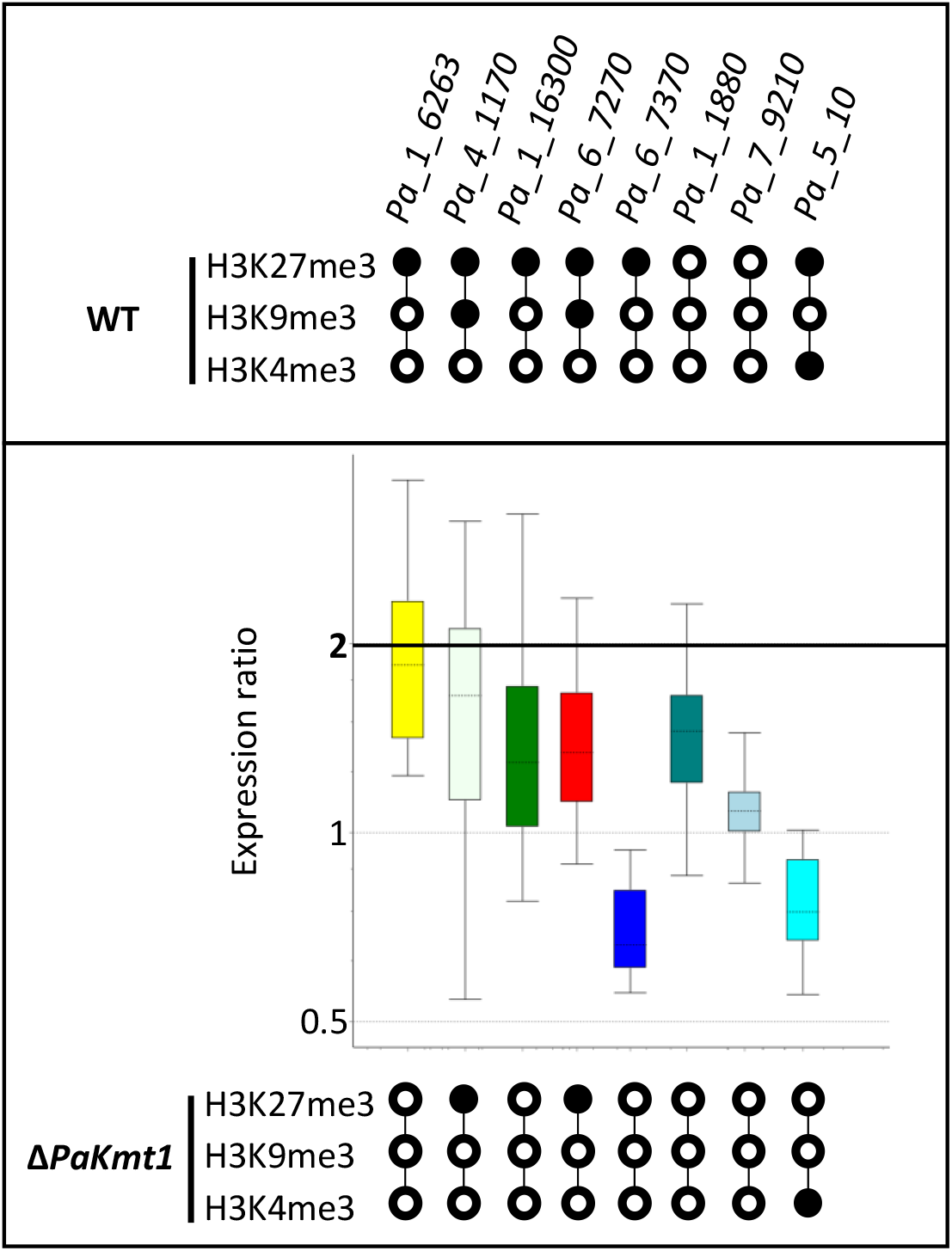
Relative expression of selected genes in the Δ*PaKmt1* mutants. Caption as in Figure 5A. No significant fold-change was assayed, except for *Pa_1_6263* and *Pa_6_7370*, which both lost the H3K27me3 mark and were upregulated and downregulated, respectively. *Pa_1_6263*: expression ratio = 1.855, p-value = 0.002; *Pa_4_1170*: expression ratio = 1.535, p-value = 0.094; *Pa_1_16300*: expression ratio = 1.393, p-value = 0.092; *Pa_6_7270*: expression ratio = 1.397, p-value = 0.035; *Pa_6_7370*: expression ratio = 0.696, p-value = 0.006; Pa_1_1880: expression ratio = 1.422, p-value = 0.034; Pa_7_9210: expression ratio = 1.082, p-value = 0.252; *Pa_5_10*: expression ratio = 0.759, p-value = 0.017.

**Supplementary Figure 8.**
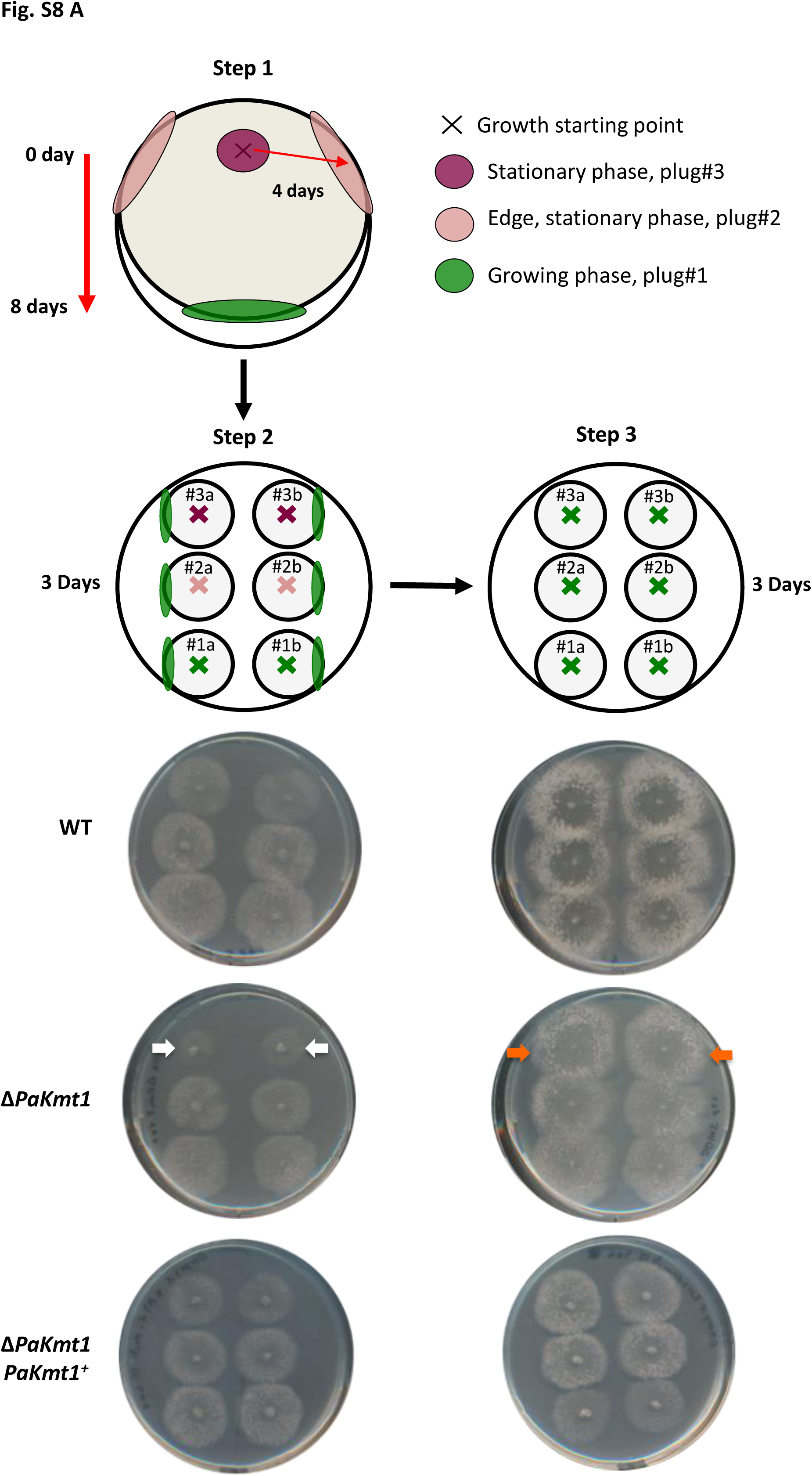
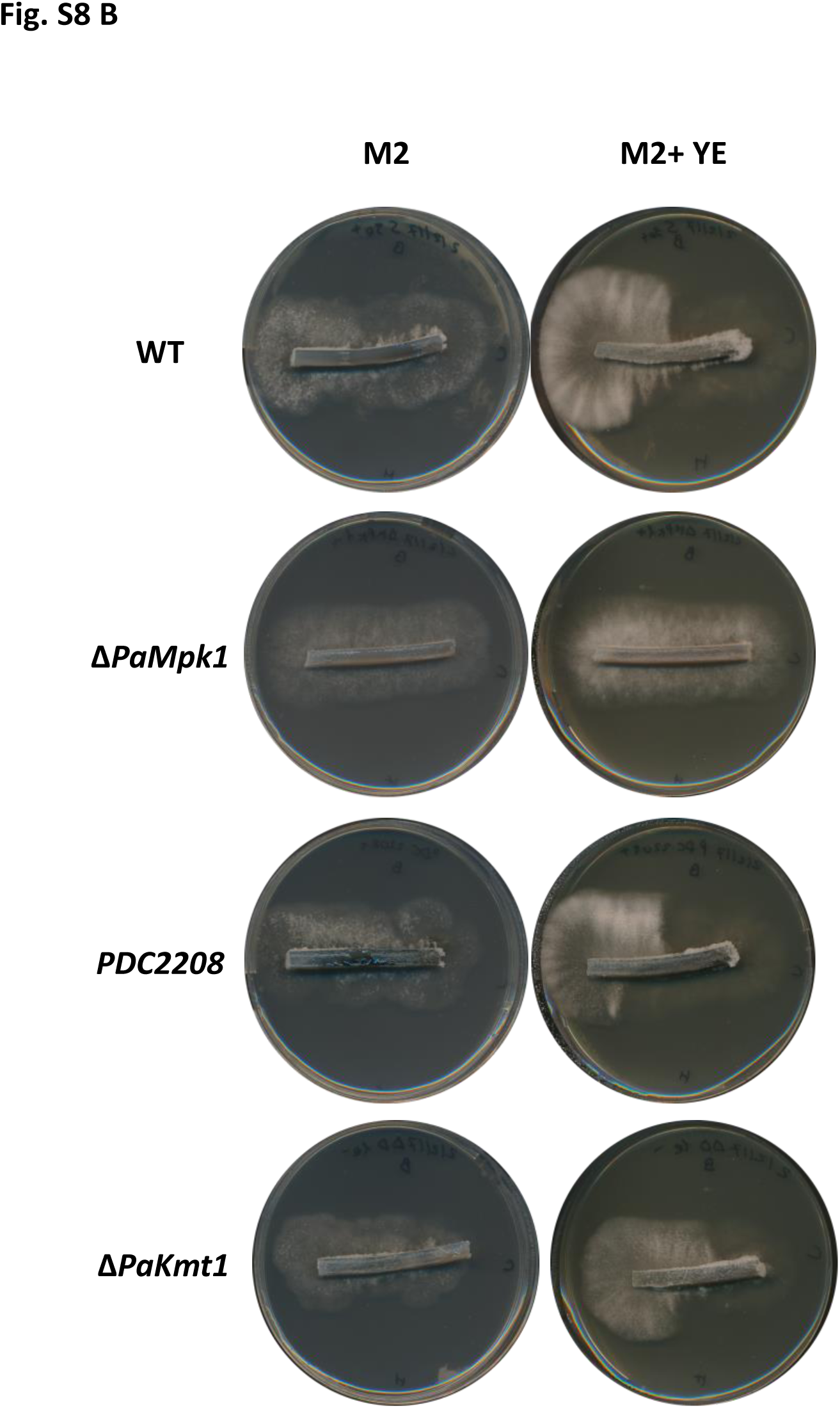
Growth features of Δ*PaKmt1* strain. A. Experimental procedure to test growth resuming capabilities of Δ*PaKmt1* strain. To set up this restart test, mycelium implants issued from germination thalli were inoculated onto a fresh M2 medium and incubated at 27°C for eight days (step 1). Two independent plugs from each location (purple cross), eight-day stationary phase (plug#3); pink cross, four-day stationary phase (plug#2); and green cross, growing phase (plug#1) were then transferred on a fresh M2 solid medium and incubated for three days at 27°C (step 2). The growing phase of each thallus originating from step 2 (green margins) were transplanted again on fresh M2 solid medium and incubated at 27°C for three days (step 3). Growth restart from stationary phase was impaired for Δ*PaKmt1* mutants, which resulted in smaller and thinner colonies than the wild-type ones (white arrows), whereas continuous growth (plug#1) was not altered. Complemented Δ*PaKmt1-PaKmt1^+^* strains behaved as wild-type strains. As control experiments (step 3), we then transferred mycelia from growing margins (marked in green, step 2) of thalli deriving from Plug#1, Plug#2 and Plug#3. In this case, Δ*PaKmt1* mutants did not show any delay to resume growth (orange arrows). **B. Crippled growth test for Δ*PaKmt1* strain.** Crippled growth (CG) process can be shown using a ‘‘band test’. Strains were incubated on M2 medium for 7 days at 27°C. Two 1-mm-width slices of agar were then inoculated onto fresh M2 media with or without yeast extract (YE) in the following method for 3 days. Picture shows actively growing apical hyphae (right) and resting stationary phase hyphae (left). The surface side of the slice on plate has been orientated at the top. CG is a degenerative process caused by C element production in the stationary phase. When inoculated on yeast extract medium, it displays slow growth, alteration of pigmentation, inability to differentiate aerial hyphae, and female sterility. Δ*PaMpk1* mutant strain is impaired for CG development while *PDC2208* strain is supposed to display CG on M2 medium without yeast extract. The single Δ*PaKmt1* mutants are not impaired for CG.

**Supplementary Figure 9.**
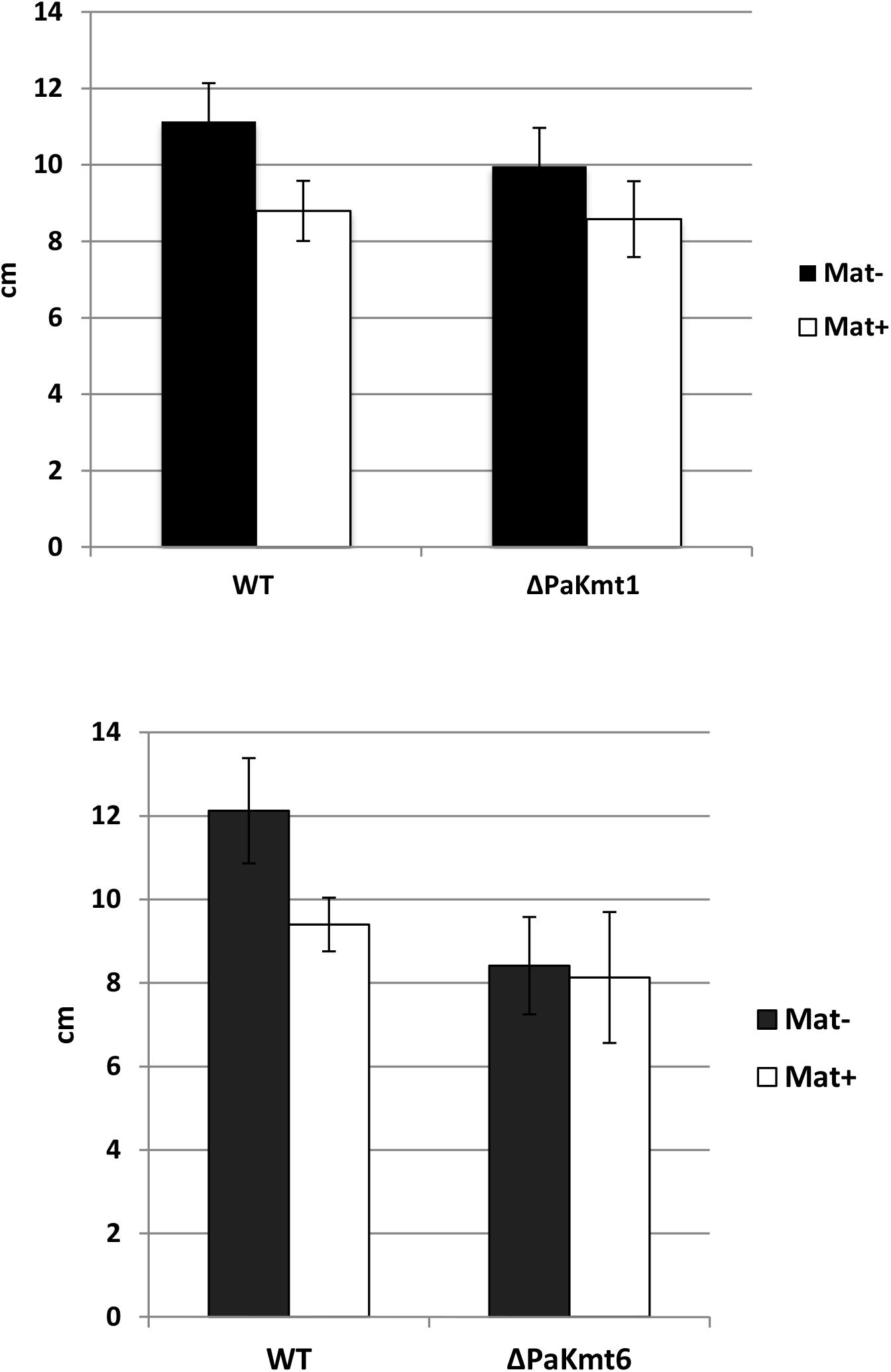
Longevity tests for Δ*PaKmt1* and Δ*PaKmt6* strains. Graphs show the maximal growth length on M2 medium at 27°C in race tube for wild-type (WT), Δ*PaKmt1* and Δ*PaKmt6* mutant strains for both mating type. Data correspond to the mean of 3 technical samples from 5 biological samples.

**Supplementary Figure 10.**
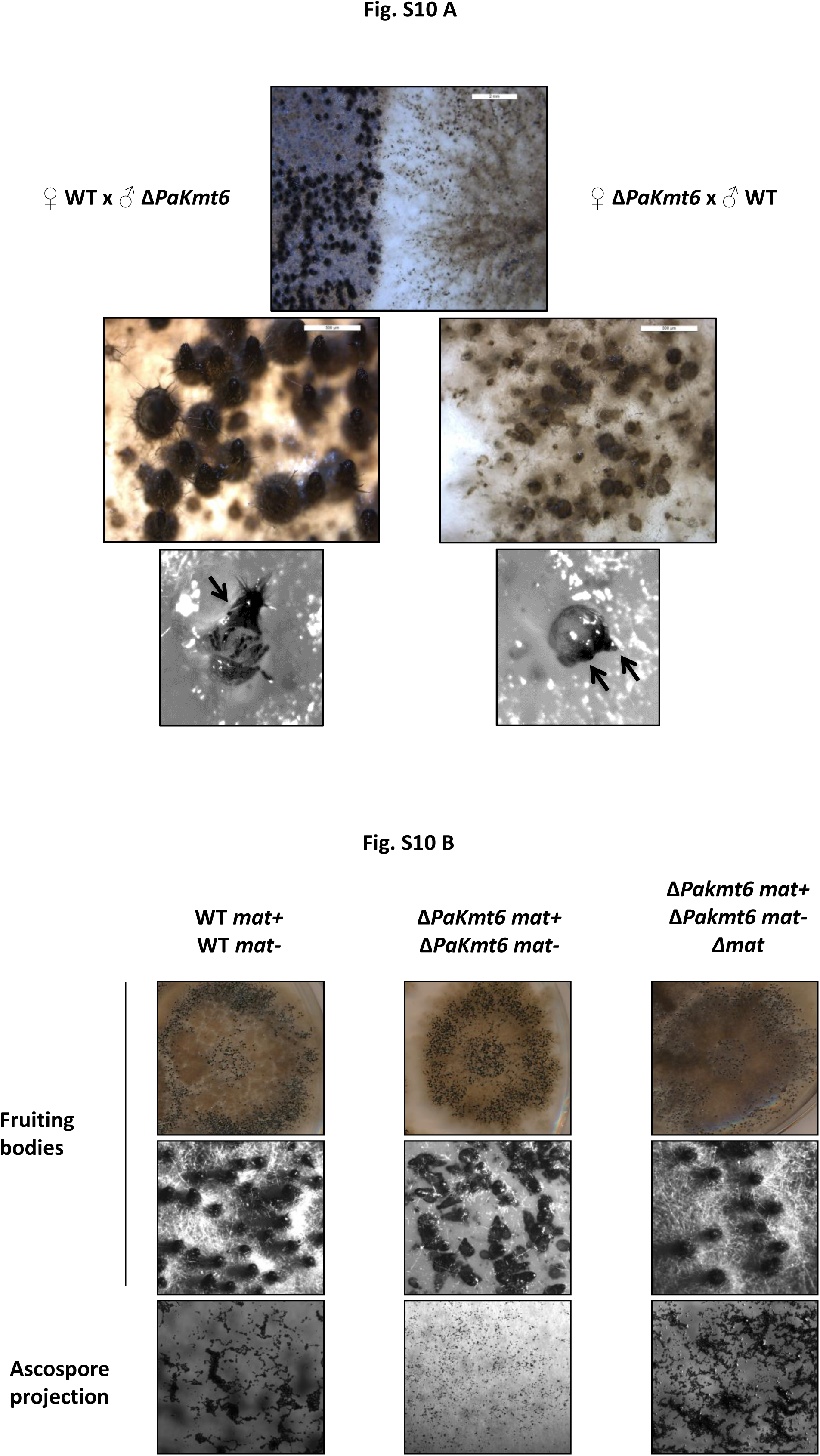
Role of *PaKmt6* during sexual development. **A. Heterozygote oriented crosses between wild-type strains (WT) and Δ*PaKmt6* mutants**. Fertilization of wild-type ascogonia with Δ*PaKmt6* male gametes results in normal fruiting body development displaying a single neck (black arrow), while fertilization of Δ*PaKmt6* ascogonia with wild-type spermatia results in fewer crippled Δ*PaKmt6*-like fruiting bodies (two necks are indicated by two black arrows). These features indicate that *PaKmt6* is a maternal gene. **B. Mosaic analyses using the Δ*mat* mutant strain.** Wild-type or Δ*PaKmt6 mat+* and *mat-* strains were mixed with or without Δ*mat* mycelium and inoculated onto fresh M2 medium. After a week of incubation on M2 medium, Δ*PaKmt6* dikaryon displayed crippled and mis-orientated perithecia, resulting in scattered and reduced ascospore production. The tricaryon Δ*PaKmt6 mat+* / Δ*PaKmt6 mat*− / Δ*mat* showed nearly wild-type restoration of mycelium growth, as well as perithecia and ascospore production. These features confirm that *PaKmt6* is a gene expressed in the maternal tissues of the perithecium and not in its zygotic tissues (centrum).

**Supplementary Figure 11.**
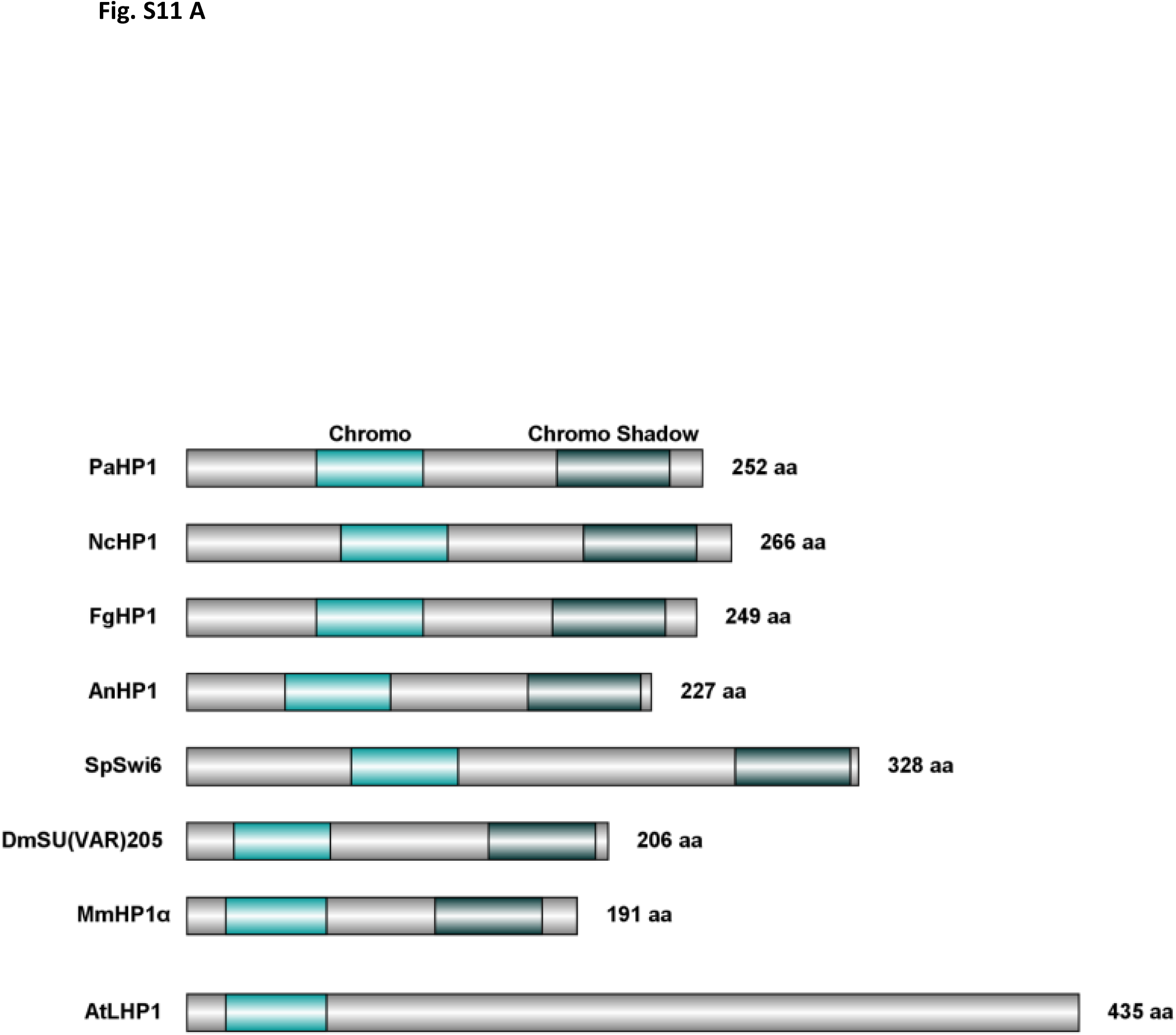
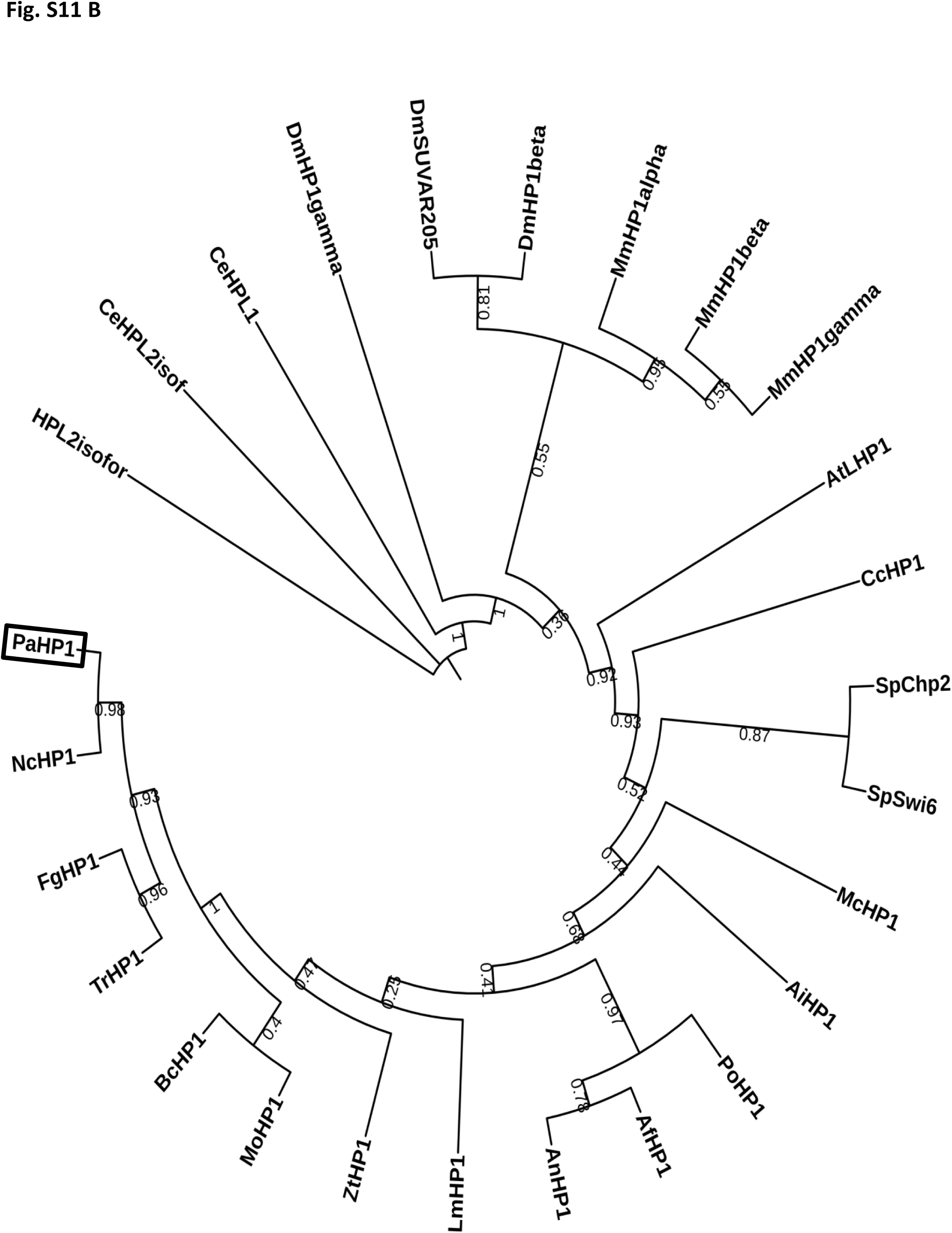
Structure and evolutionary relationships of HP1 orthologues. **A. Domain structure comparison of heterochromatin protein-1 homologues.** Sizes in amino acid (aa) are given (right). Chromodomain (turquoise, IPR023780) recognizes H3K9me2/3 histone modification. Chromo-shadow (dark green, IPR008251) is a protein-protein interaction domain. Filamentous fungi: *Podospora anserina* (Pa), *Neurospora crassa* (Nc), *Fusarium graminearum* (Fg), *Aspergillus nidulans* (An); yeasts: *Schizosaccharomyces pombe* (Sp); the fruit-fly *Drosophila melanogaster* (Dm), the mouse *Mus musculus* (Mm) and the model plant *Arabidopsis thaliana* (At). Accession numbers for proteins used in alignments are listed in Table S5. **B. Tree showing evolution of some HP1 orthologues from fungi, plant and metazoans.** The *P. anserina’s* heterochromatin protein-1 homologue is marked with a rectangle. Bootstraps are given. Filamentous fungi: *Podospora anserina* (Pa), *Neurospora crassa* (Nc), *Fusarium graminearum* (Fg), *Trichoderma reesei* (Tr), *Epichloë festucae* (Ef) *Botrytis cinerea* (Bc), Magnaporthe oryzae (Mo), *Zymoseptoria tritici* (Zt), *Leptosphaeria maculans* (Lm), *Aspergillus nidulans* (An), *Aspergilus fumigatus* (Af), *Penicillium oxalicum* (Po), *Ascobolus immersus* (Ai), *Puccinia graminis* (Pg), *Pneumocystis jirovecii* (Pj), *Conidiobolus coronatus* (Cc), *Mucor circinelloides* (Mc); yeasts: *Schizosaccharomyces pombe* (Sp) and *Cryptococcus neoformans* (Cn); the worm *Caenorhabditis elegans* (Ce), the fruit-fly *Drosophila melanogaster* (Dm), the mouse *Mus musculus* (Mm) and the model plant *Arabidopsis thaliana* (At). Accession numbers for proteins used in alignments are listed in Table S5.

**Supplementary Figure 12.**
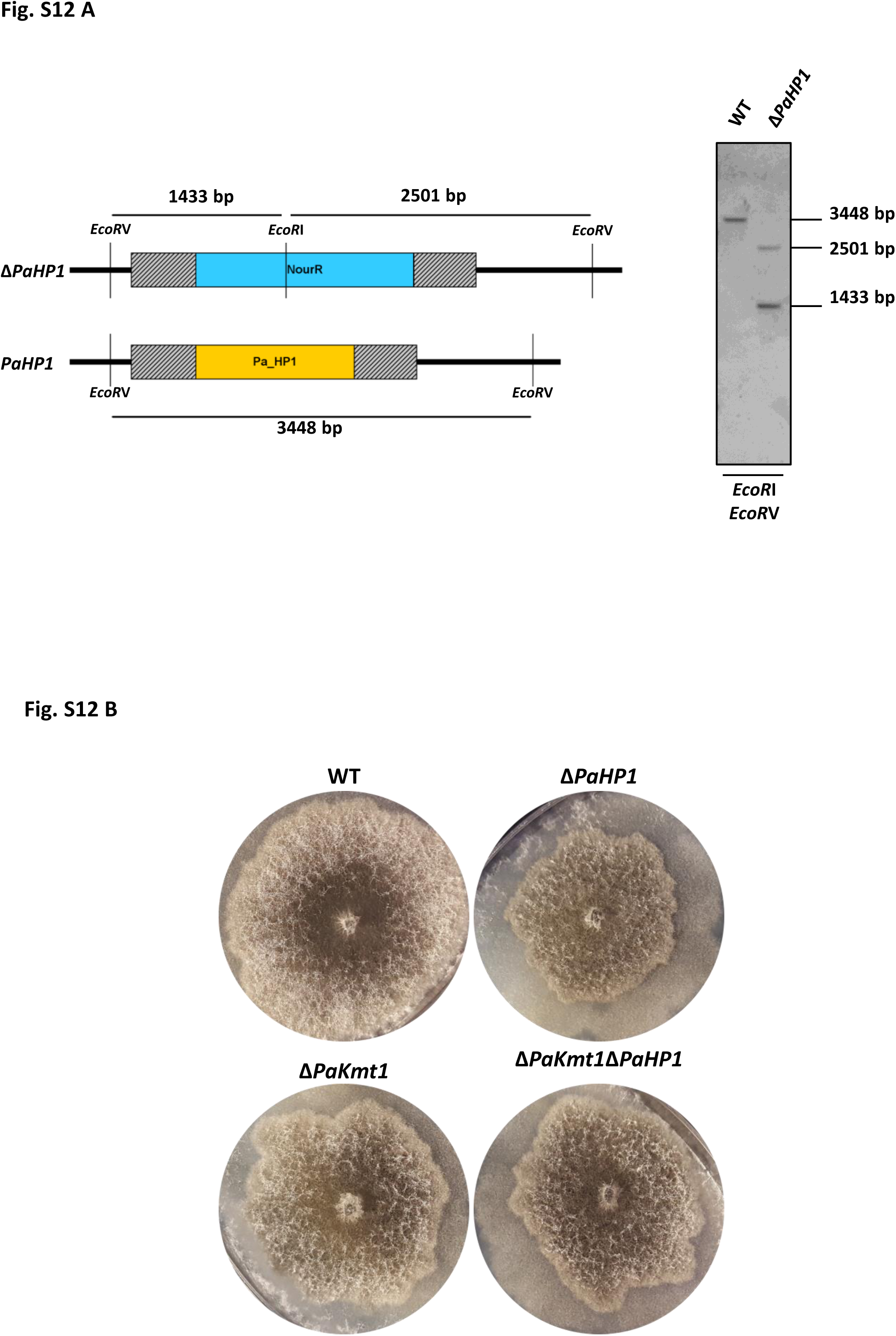
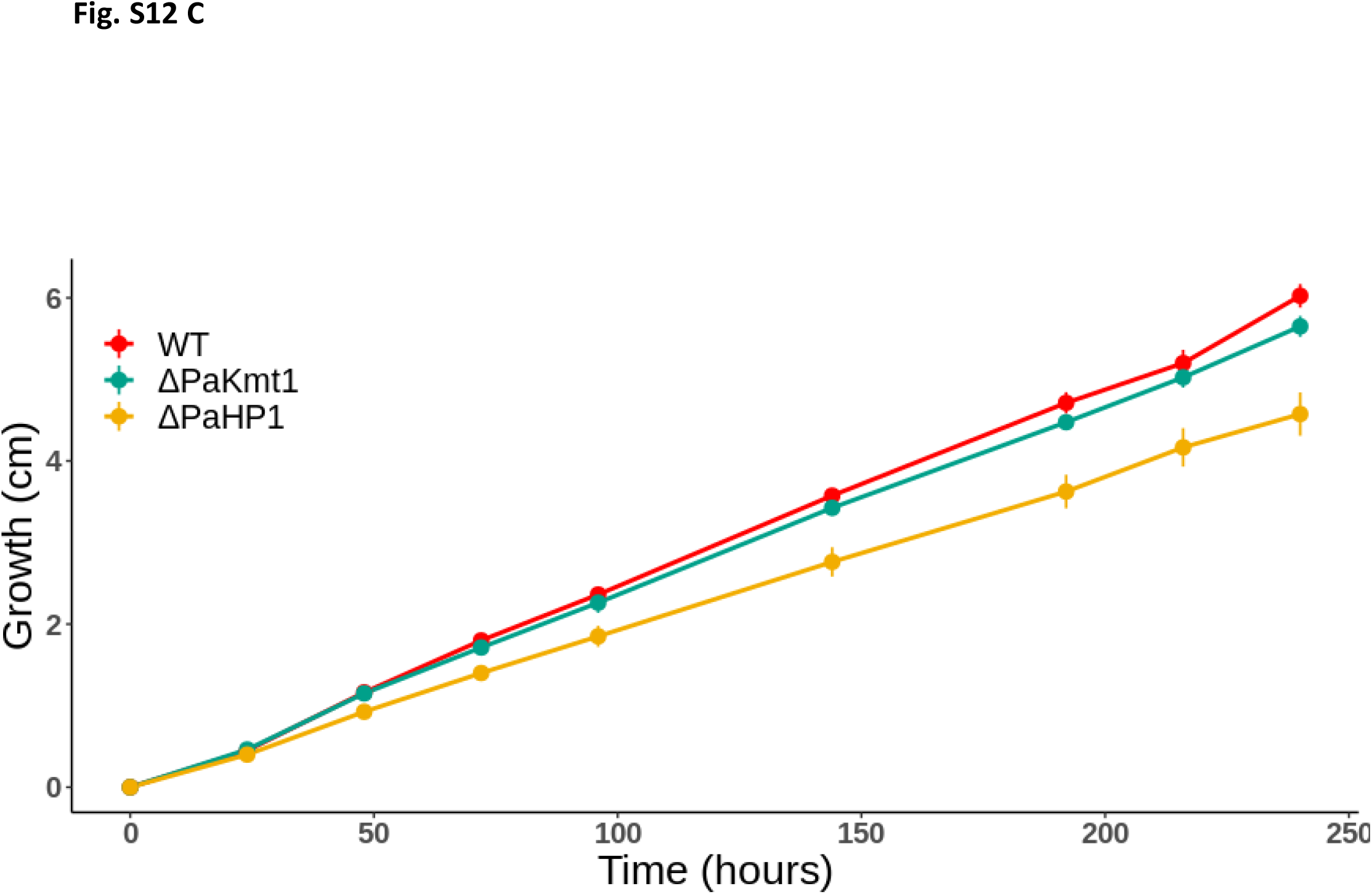
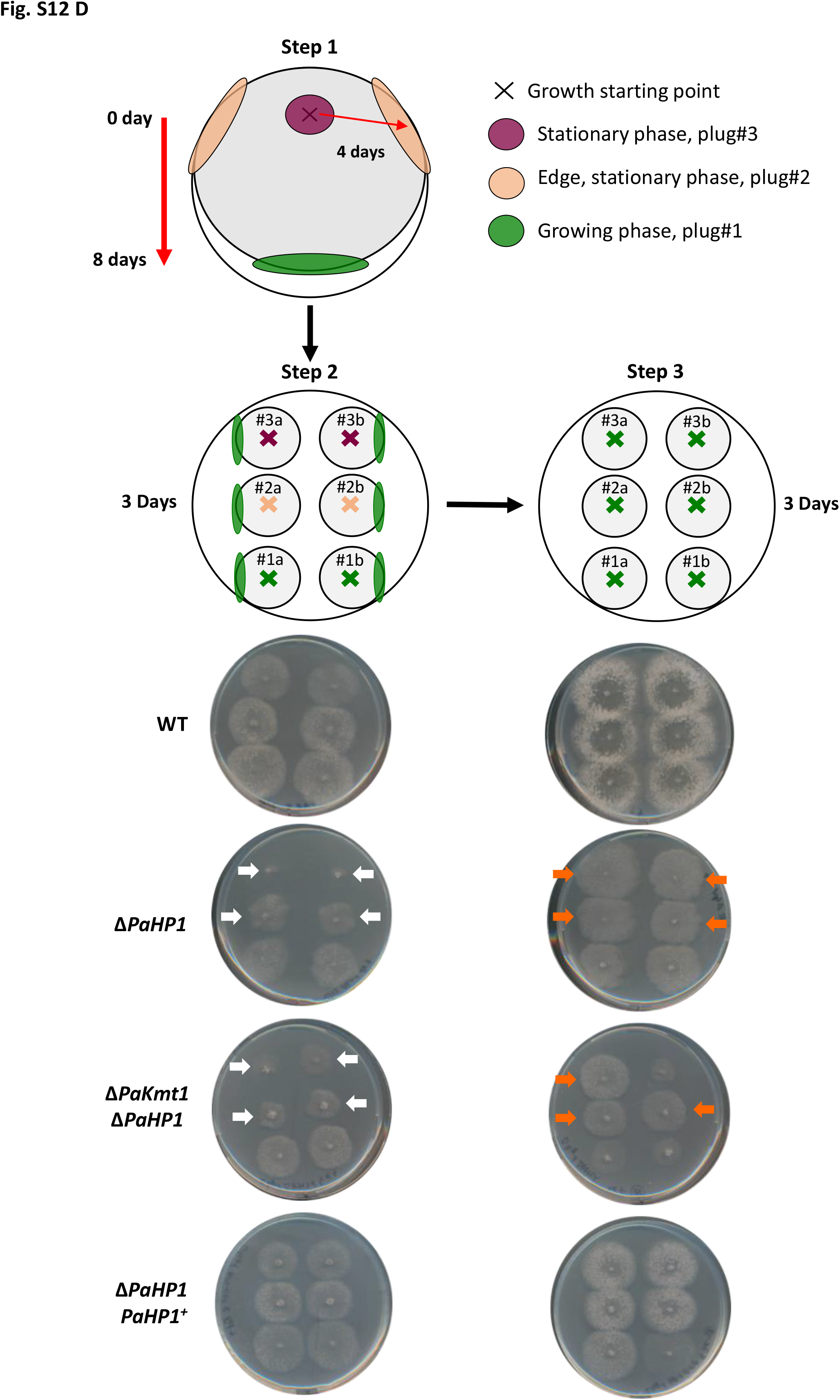
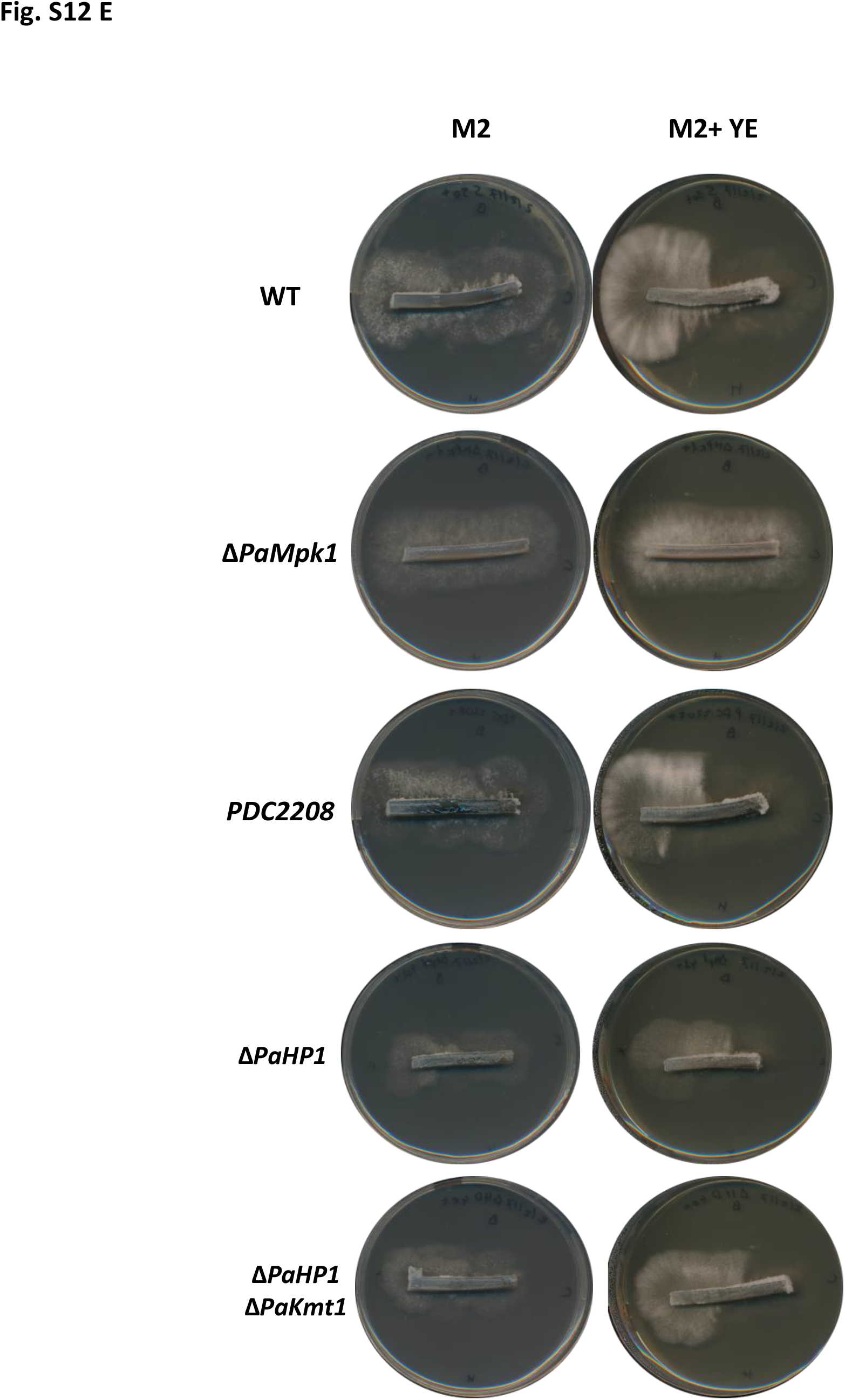
Growth features of the Δ*PaHP1* strain. **A. Molecular characterization of knockout Δ*PaHP1* mutants by Southern blot hybridization.** Replacement by homologous recombination of the wild type *PaHP1* allele by the disrupted Δ*PaHP1* allele results in the substitution of a unique 3.4 kb *Eco*RI*/Eco*RV fragment by two 1.4 and 2.5 kb *EcoRI/EcoRV* fragments as revealed by hybridization of the 5’UTR and 3’UTR digoxygenin-labeled probes (dashed rectangles *PaHP1* locus). **B. Vegetative growth of Δ*PaHP1 single* mutants and Δ*PaKmt1ΔPaHP1* double mutants compared to wild-type strains.** Δ*PaKmt1* and Δ*PaHP1* single mutants are impaired in aerial mycelium production. This defect was more pronounced for the single Δ*PaHP1* mutants*. ΔPaKmt1ΔPaHP1* double mutants show Δ*PaKmt1* morphologic phenotype. The strains have been grown on M2 medium 3 days. **C. Vegetative growth kinetics of Δ*PaHP1* mutants and Δ*PaKmt1* mutants compared to wild-type strains.** See Material and Methods section for details. **D. Experimental procedure to test growth resuming capabilities of Δ*PaHP1* and double Δ*PaHP1*Δ*PaKmt1* mutants.** For description of experimental setting see Supplementary Figure 8A. Growth restart from stationary phase (plug#2 and plug#3) was impaired for Δ*PaHP1* and double Δ*PaHP1ΔPaKmt1* mutants, which resulted in smaller and thinner colonies than the wild-type ones (white arrows), whereas continuous growth (plug#1) was not altered. Complemented Δ*PaHP1-PaHP1^+^* strains behaved as wild-type strains. As control experiments (step 3), we then transferred mycelia from growing margins (marked in green, step 2) of thalli deriving from Plug#1, Plug#2 and Plug#3. In this case, neither Δ*PaKmt1* mutants nor Δ*PaHP1*Δ*PaKmt1* mutants showed any delay to resume growth (orange arrows). **E. Crippled growth test of Δ*PaHP1* single mutants and Δ*PaHP1*Δ*PaKmt1* double mutants.** For description of experimental setting see Supplementary Figure 8B. The single Δ*PaHP1* and the double Δ*PaKmt1ΔPaHP1* mutants are not impaired for CG.

**Supplementary Figure 13.**
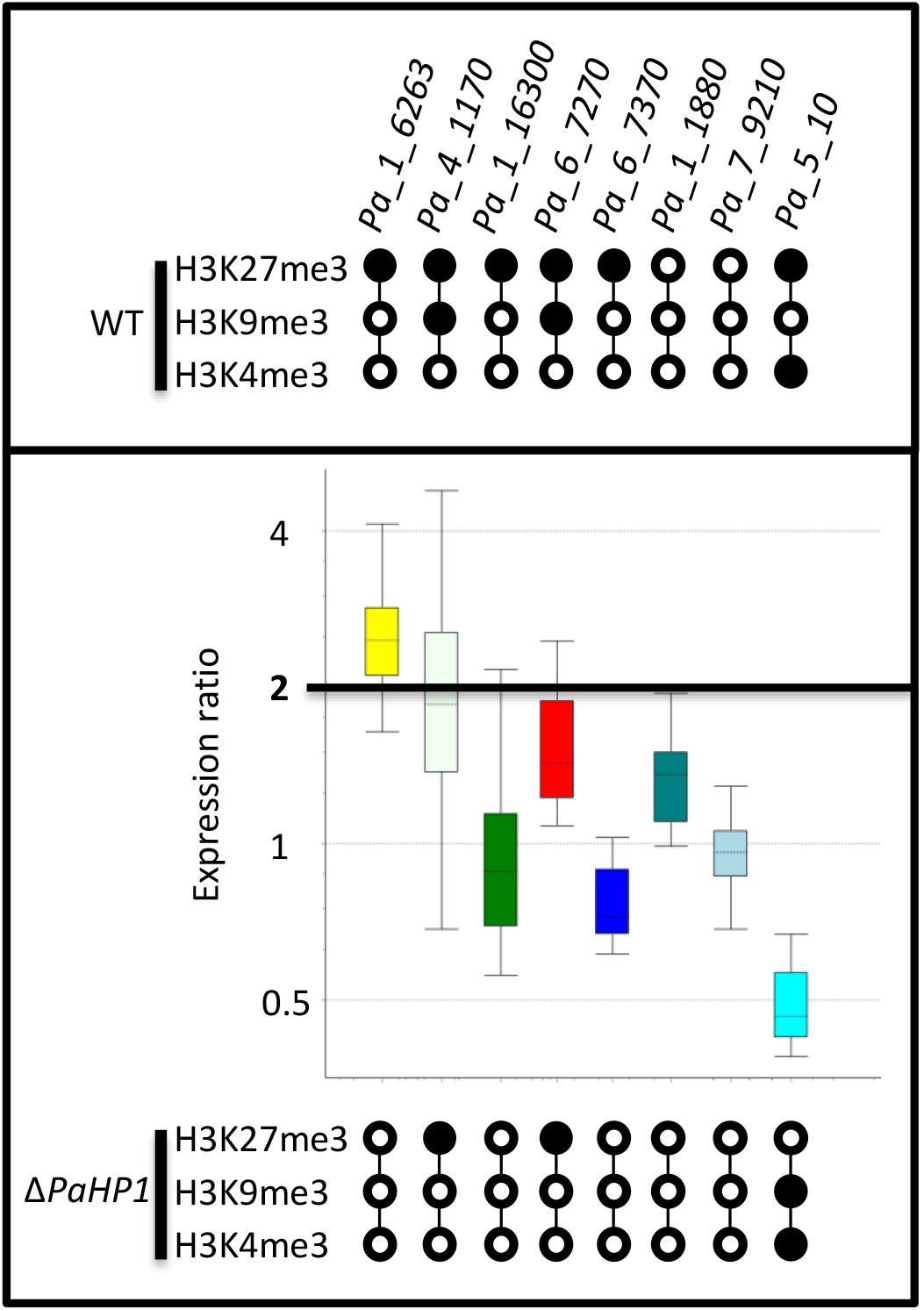
Relative expression of selected genes in the Δ*PaHP1* strain. Caption as in Figure 5A. No significant fold-change was assayed, except for *Pa_1_6263* and *Pa_5_10*, which both lost the H3K27me3 mark and were upregulated and downregulated, respectively. *Pa_1_6263:* expression ratio = 2.463, p-value = 0.004; *Pa_4_1170:* expression ratio = 1.911, p-value = 0.032; *Pa_1_16300:* expression ratio = 0.931, p-value = 0.693; *Pa_6_7270*: expression ratio = 1.527, p-value = 0.004; *Pa_6_7370*: expression ratio = 0.766, p-value = 0.009; Pa_1_1880: expression ratio = 1.313, p-value = 0.013; Pa_7_9210: expression ratio = 0.951, p-value = 0.547; *Pa_5_10:* expression ratio = 0.490, p-value = 0.006.

**Supplementary Figure 14.**
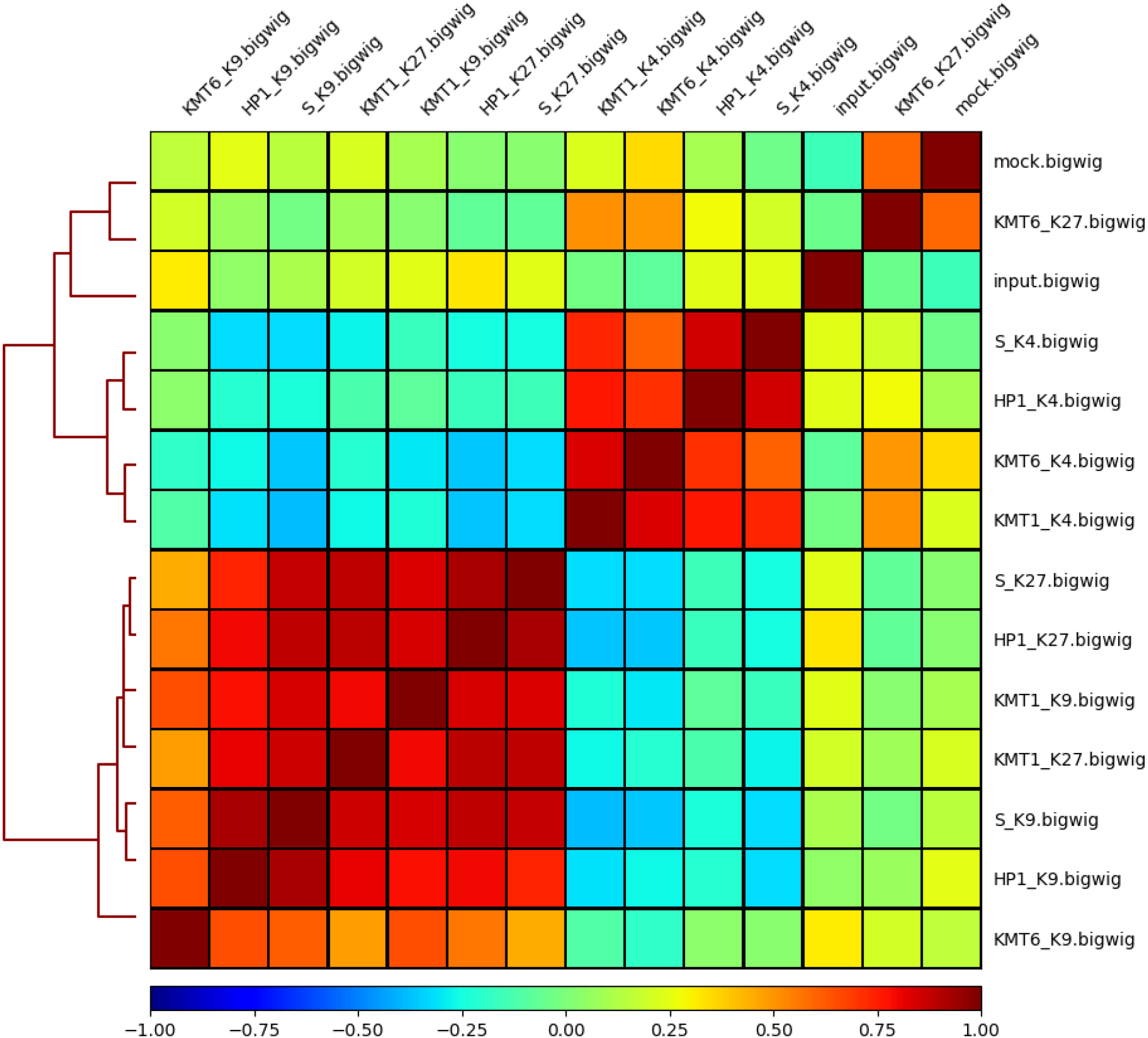
Heat map of Spearman’s correlation coefficient comparison: clustering analysis of histone marks in wild-type background (WT) and in mutant backgrounds Δ*PaKmt1, ΔPaKmt6* and Δ*PaHP1*. Mock = IP performed with GFP antibody in absence of GFP tag in *P. anserina*’s genome (see Material and methods). Raw data are given in Table S7.

**Table S1:** Main features of significantly enriched peaks. Peak numbers, sizes, and genome coverages of H3K4me3, H3K9 and H3K27me3 modifications, in wild-type background (WT) and in mutant backgrounds Δ*PaKmt1, ΔPaKmt6* and Δ*PaHP1*.

|  |  | Genome occupancy |  | Peaks detected |  | Peaks size |  | Marked features |  |  |
| --- | --- | --- | --- | --- | --- | --- | --- | --- | --- | --- |
|  |  | Mb | Genome % | Total | WT % | Average | WT % | Genes | Repeats | Repeats in CDS |
| H3K4me3 | WT | 3,84 | 10,98 | 3 050,00 | 100,00 | 1 259,52 | 100,00 | 3 503,00 | 63,00 | 49,00 |
|  | ΔPaHP1 | 3,66 | 10,47 | 3 138,00 | 102,89 | 1 167,59 | 92,70 | 3 587,00 | 64,00 | 52,00 |
|  | ΔPaKmt1 | 4,15 | 11,85 | 3 307,00 | 108,43 | 1 253,66 | 99,53 | 3 793,00 | 67,00 | 58,00 |
|  | ΔPaKMT6 | 4,19 | 11,98 | 3 433,00 | 112,56 | 1 220,96 | 96,94 | 3 983,00 | 85,00 | 69,00 |
| H3K9me3 | WT | 1,03 | 2,94 | 1 090,00 | 100,00 | 945,03 | 100,00 | 537,00 | 728,00 | 213,00 |
|  | ΔPaHP1 | 0,66 | 1,88 | 419,00 | 38,44 | 1 571,95 | 166,34 | 93,00 | 861,00 | 71,00 |
|  | ΔPaKmt1 | - | - | - | - | - | - | - | - | - |
|  | ΔPaKMT6 | 0,06 | 0,18 | 85,00 | 7,80 | 730,72 | 77,32 | 2,00 | 129,00 | 4,00 |
| H3K27me3 | WT | 6,75 | 19,28 | 1 941,00 | 100,00 | 3 476,75 | 100,00 | 2 992,00 | 1 042,00 | 307,00 |
|  | ΔPaHP1 | 1,80 | 5,15 | 1 443,00 | 74,34 | 1 250,34 | 35,96 | 1 027,00 | 551,00 | 239,00 |
|  | ΔPaKmt1 | 1,56 | 4,45 | 1 326,00 | 68,32 | 1 174,28 | 33,78 | 1 003,00 | 538,00 | 259,00 |
|  | ΔPaKMT6 | 0,01 | 0,04 | 47,00 | 2,42 | 266,49 | 7,66 | 38,00 | - | - |

**Table S2.**
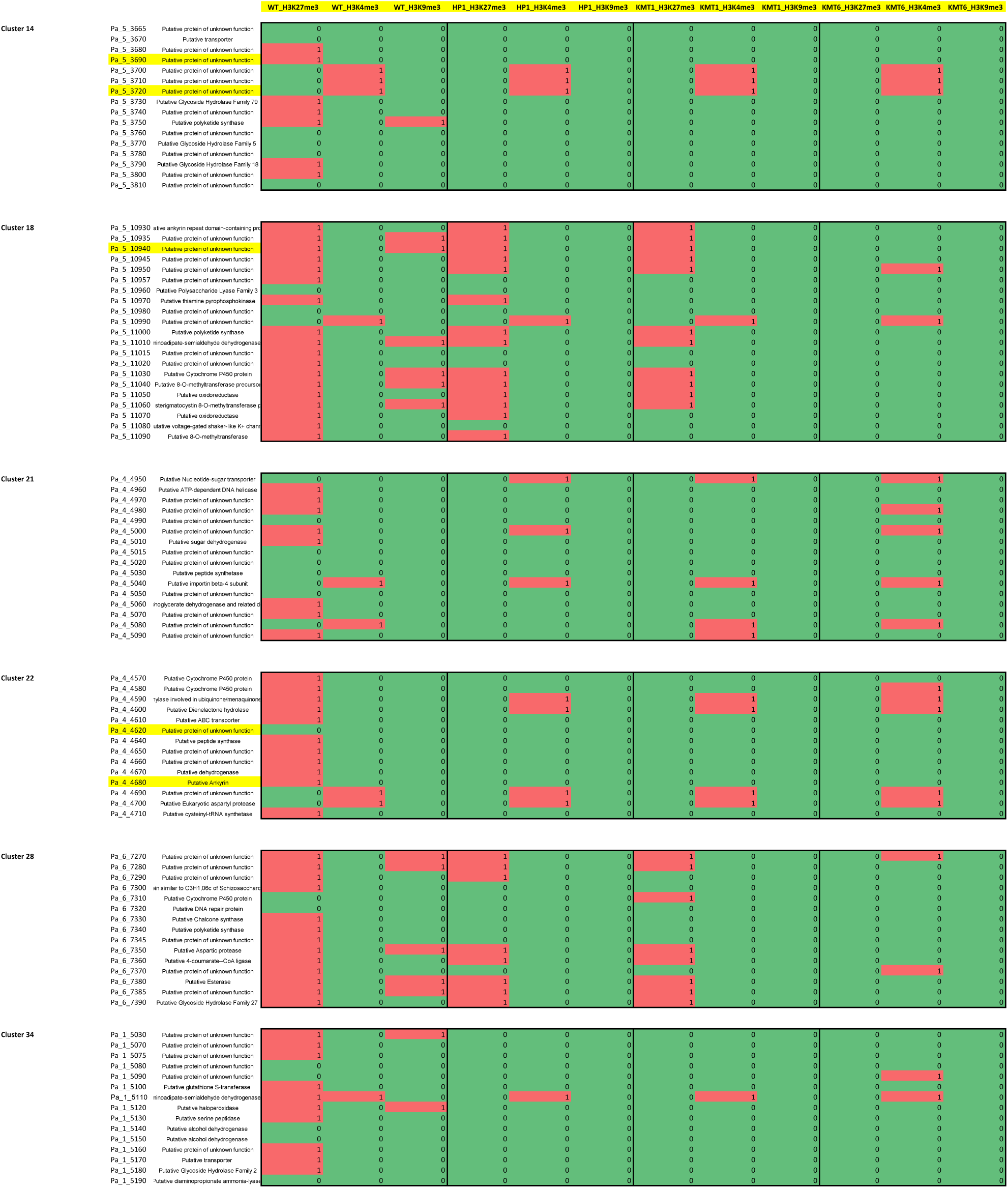
**Distribution of H3K4me3, H3K9me3 and H3K27me3 on the complete set of *P. anserina*’s** gene in wild-type background (WT) and in Δ*PaKmt1, ΔPaKmt6* and Δ*PaHP1* mutants. List of genes where H3K4me3 and H3K27me3 can be found overlapping in the wild-type background. Distribution of H3K4me3, H3K9me3 and H3K27me3 on a sub-set of *P. anserina’s* secondary metabolite gene clusters in wild-type background (WT) and in *PaKmt1*, Δ*PaKmt6* and Δ*PaHP1* mutants.

**Table S3.** Cq for RT-qPCR experiments and geNorm analysis of candidate normalization genes.

| replicate number | P. anserina strain | mating type | culture condition |
| --- | --- | --- | --- |
| 1-1 | *wild-type* | *mat+* | 60 h, 27°C, liquid medium as for ChIPseq, see<br>M & M for details |
| 1-2 | *wild-type* | *mat+* |  |
| 1-3 | *wild-type* | *mat+* |  |
| 1-4 | *wild-type* | *mat+* |  |
| 1-5 | *wild-type* | *mat+* |  |
| 2-1 | $\Delta Kmt1$ | *mat+* | |
| 2-2 | $\Delta Kmt1$ | *mat+* | |
| 2-3 | $\Delta Kmt1$ | *mat+* | |
| 2-4 | $\Delta Kmt1$ | *mat+* | |
| 2-5 | $\Delta Kmt1$ | *mat+* | |
| 3-1 | $\Delta Kmt6$ | *mat+* | |
| 3-2 | $\Delta Kmt6$ | *mat+* | |
| 3-3 | $\Delta Kmt6$ | *mat+* | |
| 3-4 | $\Delta Kmt6$ | *mat+* | |
| 3-5 | $\Delta Kmt6$ | *mat+* | |
| 4-1 | $\Delta Hp1$ | *mat+* | |
| 4-2 | $\Delta Hp1$ | *mat+* | |
| 4-3 | $\Delta Hp1$ | *mat+* | |
| 4-4 | $\Delta Hp1$ | *mat+* | |
| 4-5 | $\Delta Hp1$ | *mat+* | |

Table S3
|  | AS1 | CIT1 | GPD | PAH1 | PDF2 | TBP | TIP | UBC | Pa_6_7270 | Pa_6_7370 | Pa_1_6263 | Pa_1_16300 | Pa_4_1170 | Pa_1_1880 | Pa_5_10 | Pa_7_9210 | Pa_1_5210 |
| --- | --- | --- | --- | --- | --- | --- | --- | --- | --- | --- | --- | --- | --- | --- | --- | --- | --- |
| 1-1 | 15,39 | 19,97 | 13,74 | 17,64 | 18,31 | 17,86 | 21,79 | 15,14 | 22,44 | 22,03 | 29,7 | 28,35 | 29,24 | 21,65 | 22,31 | 21,97 | 22,27 |
| 1-2 | 14,99 | 19,35 | 13,49 | 17,52 | 18,11 | 17,78 | 21,5 | 15,21 | 22,5 | 21,3 | 30,02 | 29,01 | 29,6 | 21,62 | 22,21 | 21,46 | 23,1 |
| 1-3 | 14,79 | 19,34 | 13,8 | 18,1 | 17,74 | 18,69 | 21,54 | 14,8 | 22,99 | 21,34 | 29,38 | 27,45 | 29,47 | 21,6 | 22,54 | 21,44 | 23,54 |
| 1-4 | 15,23 | 19,46 | 13,9 | 17,94 | 18,04 | 18,45 | 21,83 | 15,1 | 22,98 | 21,55 | 29,93 | 27,73 | 28,05 | 21,38 | 22,7 | 22,09 | 23,29 |
| 1-5 | 15,25 | 19,6 | 13,71 | 17,85 | 18,05 | 17,98 | 21,59 | 14,94 | 22,37 | 21,98 | 29,47 | 28,1 | 28,86 | 21,29 | 22,22 | 21,71 | 21,68 |
| 2-1 | 14,88 | 18,78 | 13,22 | 17,61 | 17,9 | 17,76 | 21,43 | 14,96 | 22,15 | 21,75 | 28,71 | 27,72 | 28,25 | 21,19 | 22,43 | 21,26 | 21,38 |
| 2-2 | 15,2 | 19,7 | 13,8 | 17,83 | 18,06 | 18,21 | 21,89 | 15,08 | 22,13 | 22,3 | 27,93 | 27,31 | 29,1 | 20,79 | 22,77 | 21,61 | 20,85 |
| 2-3 | 15,12 | 19,26 | 13,59 | 17,92 | 17,95 | 18,2 | 21,59 | 15,09 | 22,16 | 22,18 | 28,79 | 27,66 | 27,6 | 21,07 | 22,95 | 21,57 | 21,67 |
| 2-4 | 15,25 | 19,93 | 13,72 | 17,89 | 18,09 | 18,08 | 21,43 | 15,06 | 21,99 | 22,21 | 28,39 | 27,23 | 28,2 | 20,59 | 22,81 | 21,9 | 20,47 |
| 2-5 | 14,97 | 19,24 | 13,77 | 18,26 | 17,84 | 18,34 | 21,36 | 15,11 | 22,08 | 22,02 | 28,77 | 27,51 | 27,79 | 20,88 | 22,71 | 21,34 | 21,64 |
| 3-1 | 20,02 | 24,01 | 18,72 | 22,36 | 23,43 | 22,06 | 27,13 | 19,97 | 24,93 | 24,32 | 29,35 | 30,96 | 31,48 | 25,64 | 26,26 | 26,82 | 26,84 |
| 3-2 | 20,04 | 24,22 | 18,82 | 22,73 | 23,45 | 22,45 | 27,5 | 19,91 | 25,07 | 24,43 | 29,57 | 30,88 | 31,89 | 26,36 | 26,45 | 27,13 | 26,9 |
| 3-3 | 20,26 | 24,51 | 19,25 | 23,3 | 23,77 | 22,47 | 27,97 | 19,96 | 25,32 | 24,26 | 30,25 | 30,23 | 32,27 | 26,7 | 26,61 | 27,45 | 27,07 |
| 3-4 | 19,62 | 23,75 | 18,57 | 22,48 | 23,12 | 21,96 | 27,6 | 19,32 | 24,63 | 24,01 | 29,24 | 29,96 | 31,16 | 26,05 | 26,25 | 26,81 | 26,43 |
| 3-5 | 19,69 | 23,82 | 17,92 | 22,15 | 22,43 | 21,04 | 27,15 | 18,67 | 25,02 | 22,1 | 30,43 | 29,24 | 31,28 | 25,95 | 25,29 | 27,55 | 26,62 |
| 4-1 | 14,95 | 19,35 | 13,51 | 17,66 | 18,39 | 17,63 | 21,8 | 15,92 | 22,14 | 21,98 | 27,99 | 28,33 | 27,54 | 21,25 | 23,32 | 21,68 | 21,7 |
| 4-2 | 15,16 | 19,9 | 13,76 | 18,03 | 18,38 | 18,21 | 21,55 | 15,86 | 22,11 | 22,13 | 28,27 | 28,42 | 28,21 | 21,28 | 23,64 | 21,75 | 21,72 |
| 4-3 | 15,12 | 19,73 | 13,85 | 18,13 | 18,54 | 18,17 | 21,79 | 15,91 | 22,3 | 22,1 | 28,79 | 28,47 | 28,49 | 21,36 | 23,68 | 21,89 | 22,51 |
| 4-4 | 15,21 | 20,14 | 13,8 | 18,06 | 18,42 | 18,21 | 22,08 | 15,68 | 22,26 | 22,28 | 28,41 | 28,8 | 28,83 | 21,29 | 23,73 | 22,07 | 21,65 |
| 4-5 | 15,6 | 20,16 | 13,72 | 18,09 | 18,6 | 18,04 | 22,26 | 15,68 | 22,21 | 22,43 | 28,04 | 28,21 | 27,34 | 21,13 | 23,81 | 22,48 | 21,08 |
| Cqmin | 14,79 | 18,78 | 13,22 | 17,52 | 17,74 | 17,63 | 21,36 | 14,8 |  |  |  |  |  |  |  |  |  |
| Efficiency | 1,959 | 1,963 | 1,948 | 1,866 | 1,917 | 2,057 | 1,878 | 1,942 | 2,01 | 1,989 | 1,763 | 1,817 | 1,753 | 1,953 | 1,94 | 1,935 | 1,828 |

|  | AS1 | CIT1 | GPD | PAH1 | PDF2 | TBP | TIP | UBC |
| --- | --- | --- | --- | --- | --- | --- | --- | --- |
| 1-1 | 0,668004424 | 0,448151489 | 0,706990771 | 0,927877414 | 0,690089462 | 0,847141803 | 0,762624753 | 0,797985984 |
| 1-2 | 0,874164393 | 0,680824897 | 0,835239419 | 1 | 0,786013162 | 0,897459494 | 0,915551161 | 0,761759392 |
| 1-3 | 1 | 0,685432404 | 0,679263702 | 0,696420023 | 1 | 0,465555703 | 0,892760145 | 1 |
| 1-4 | 0,743883367 | 0,632141345 | 0,635447261 | 0,769515038 | 0,822646712 | 0,553538562 | 0,743640568 | 0,819455232 |
| 1-5 | 0,733946087 | 0,57518209 | 0,721275887 | 0,813952683 | 0,817310625 | 0,776905146 | 0,865067537 | 0,911265878 |
| 2-1 | 0,941275816 | 1 | 1 | 0,945405125 | 0,901115504 | 0,910499145 | 0,956844377 | 0,899249332 |
| 2-2 | 0,759042129 | 0,537666929 | 0,679263702 | 0,824171119 | 0,812009152 | 0,658148826 | 0,716046685 | 0,8304055 |
| 2-3 | 0,800992844 | 0,723432901 | 0,781361642 | 0,7791756 | 0,872266827 | 0,662912875 | 0,865067537 | 0,824912196 |
| 2-4 | 0,733946087 | 0,460406722 | 0,716482395 | 0,793894318 | 0,796310167 | 0,72284398 | 0,956844377 | 0,841502095 |
| 2-5 | 0,886000162 | 0,733257751 | 0,692988577 | 0,630268187 | 0,936996112 | 0,599243943 | 1 | 0,814034366 |
| 3-1 | 0,029693398 | 0,029378312 | 0,025542345 | 0,048841351 | 0,02465359 | 0,040961145 | 0,026349747 | 0,032340855 |
| 3-2 | 0,029296734 | 0,025498438 | 0,023894716 | 0,038774871 | 0,024334797 | 0,030917952 | 0,020869358 | 0,033654757 |
| 3-3 | 0,025268045 | 0,020968467 | 0,01793819 | 0,02717257 | 0,019760078 | 0,030475162 | 0,015519301 | 0,032556222 |
| 3-4 | 0,038857394 | 0,035009516 | 0,028229235 | 0,045318786 | 0,030164285 | 0,04402461 | 0,019594741 | 0,049786718 |
| 3-5 | 0,037070741 | 0,033395019 | 0,043544361 | 0,055677421 | 0,047260933 | 0,08548129 | 0,026019716 | 0,07664353 |
| 4-1 | 0,897996182 | 0,680824897 | 0,824174558 | 0,916373181 | 0,65508194 | 1 | 0,757833748 | 0,475511328 |
| 4-2 | 0,779735413 | 0,469817581 | 0,697624888 | 0,727503461 | 0,659358862 | 0,658148826 | 0,887151596 | 0,494829771 |
| 4-3 | 0,800992844 | 0,526896987 | 0,656990304 | 0,683508469 | 0,594158493 | 0,677412931 | 0,762624753 | 0,478677881 |
| 4-4 | 0,753955193 | 0,39960268 | 0,679263702 | 0,714015623 | 0,642416909 | 0,658148826 | 0,635241959 | 0,557623004 |
| 4-5 | 0,580032204 | 0,394248443 | 0,716482395 | 0,700777849 | 0,571406245 | 0,744001721 | 0,567118704 | 0,557623004 |

**Table S3**
|  | AS1 | CIT1 | GPD | PAH1 | PDF2 | TBP | TIP | UBC | Pa_6_7270 | Pa_6_7370 | Pa_1_6263 | Pa_1_16300 | Pa_4_1170 | Pa_1_1880 | Pa_5_10 | Pa_7_9210 | Pa_1_5210 |
| --- | --- | --- | --- | --- | --- | --- | --- | --- | --- | --- | --- | --- | --- | --- | --- | --- | --- |
| 1-1 | Gene with an | Gene with an | Gene with an | Gene with an | Gene with an | Gene with an | Gene with an | Gene with an | N/A | Gene with an | Gene with an | Gene with an | N/A | N/A | N/A | N/A | N/A |
| 1-2 |  |  |  |  |  |  |  |  | 35,44 |  |  |  | N/A | N/A | N/A | 33,85 | N/A |
| 1-3 |  |  |  |  |  |  |  |  | N/A |  |  |  | 38,78 | N/A | 36,14 | 33,92 | N/A |
| 1-4 |  |  |  |  |  |  |  |  | 36,27 |  |  |  | N/A | N/A | N/A | N/A | N/A |
| 1-5 |  |  |  |  |  |  |  |  | 36,84 |  |  |  | N/A | N/A | 35,55 | N/A | 34,19 |
| 2-1 |  |  |  |  |  |  |  |  | 34,94 |  |  |  | 39,87 | 35,43 | 37,18 | 33,53 | 33,53 |
| 2-2 |  |  |  |  |  |  |  |  | 39,25 |  |  |  | N/A | 35,32 | N/A | N/A | N/A |
| 2-3 |  |  |  |  |  |  |  |  | N/A |  |  |  | N/A | N/A | 35,28 | N/A | N/A |
| 2-4 |  |  |  |  |  |  |  |  | 33,14 |  |  |  | 38,68 | 32,77 | 33,69 | 31,87 | 32,2 |
| 2-5 |  |  |  |  |  |  |  |  | 37,23 |  |  |  | 38,54 | 34,49 | 33,96 | 32,35 | 34,57 |
| 3-1 |  |  |  |  |  |  |  |  | 35,01 |  |  |  | N/A | N/A | N/A | N/A | 33,95 |
| 3-2 |  |  |  |  |  |  |  |  | 34,72 |  |  |  | N/A | N/A | N/A | N/A | N/A |
| 3-3 |  |  |  |  |  |  |  |  | 34,57 |  |  |  | N/A | N/A | 35,13 | 33,52 | N/A |
| 3-4 |  |  |  |  |  |  |  |  | 33,96 |  |  |  | N/A | N/A | N/A | 34,16 | N/A |
| 3-5 |  |  |  |  |  |  |  |  | 33,57 |  |  |  | N/A | 34,47 | 34,69 | 32,54 | N/A |
| 4-1 |  |  |  |  |  |  |  |  | 32,74 |  |  |  | 40,01 | 33,75 | 34,26 | 32,77 | 33,23 |
| 4-2 |  |  |  |  |  |  |  |  | 30,21 |  |  |  | 37,88 | 31,96 | 31,6 | 30,19 | 31,17 |
| 4-3 |  |  |  |  |  |  |  |  | 31,31 |  |  |  | 38,04 | 33,74 | 33,14 | 30,89 | 33,32 |
| 4-4 |  |  |  |  |  |  |  |  | 32,11 |  |  |  | 40,45 | 34,47 | 33,5 | 31,14 | 32,8 |
| 4-5 |  |  |  |  |  |  |  |  | 32,37 |  |  |  | 39,47 | 34,36 | 33,33 | 32,82 | 32,88 |

**Table S4.**
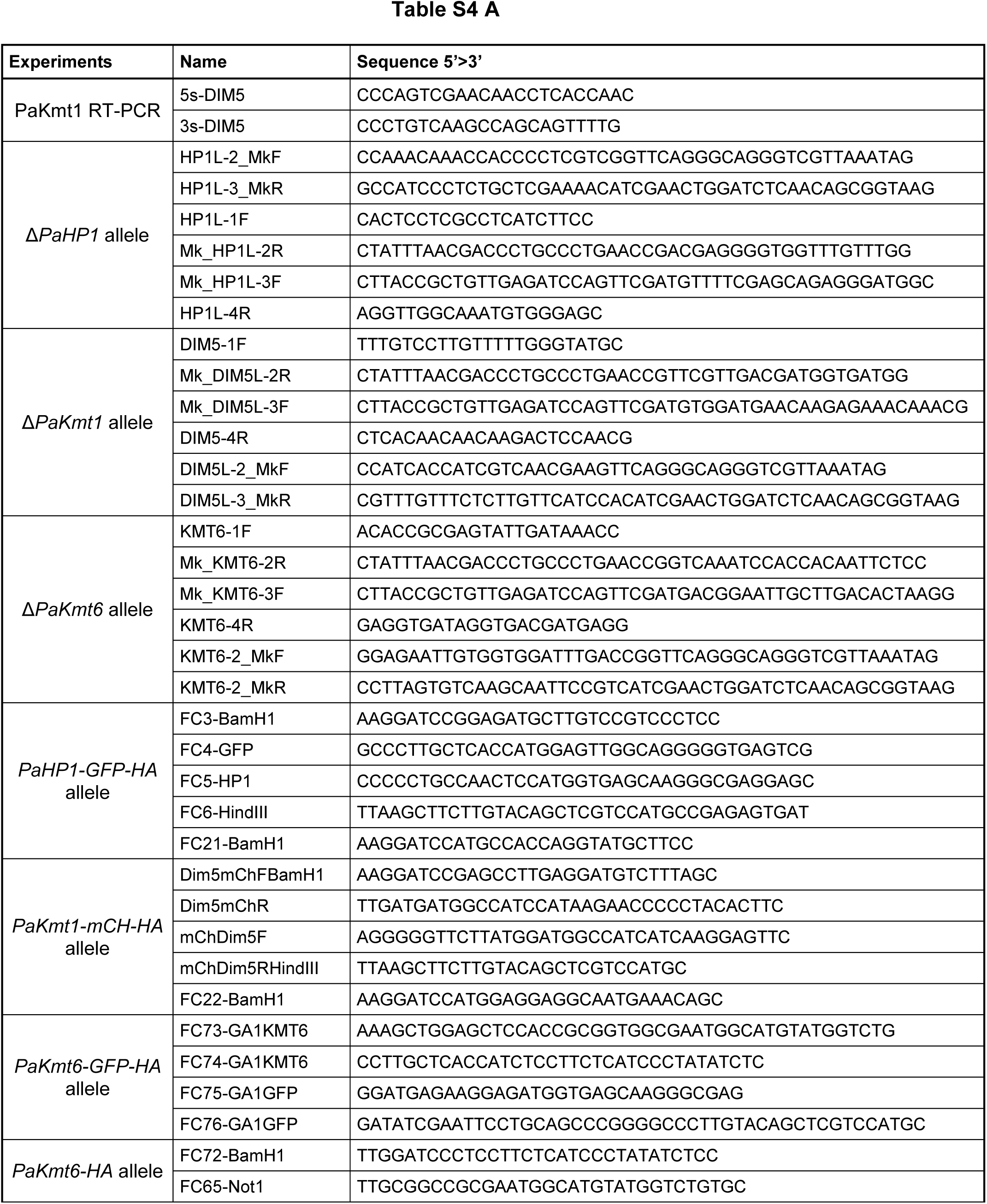
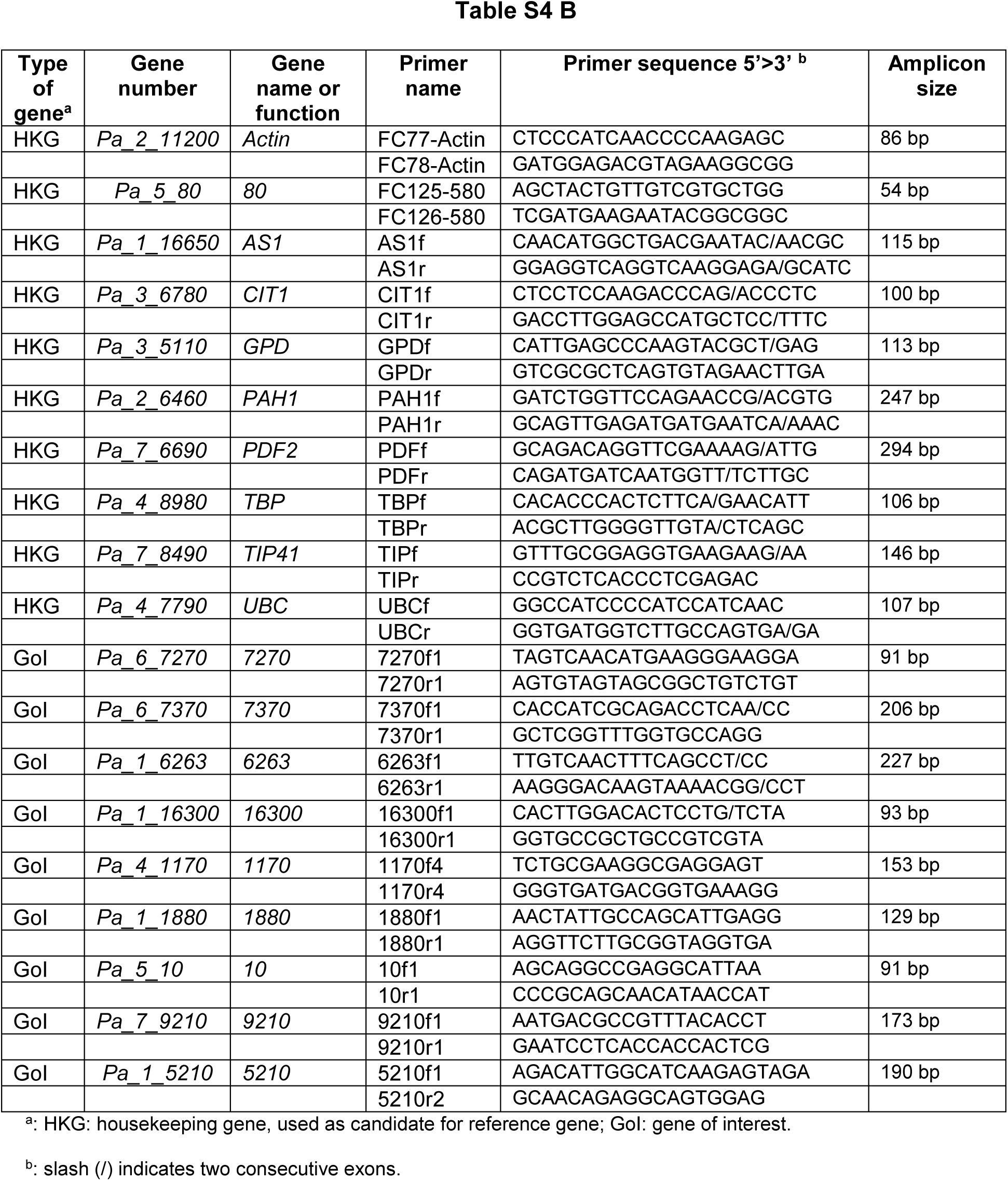
Primers used for PCR experiments. A. Primers used for RT-PCR and allele construction experiments. B. Primers used for RT-qPCR experiments.

**Table S5.**
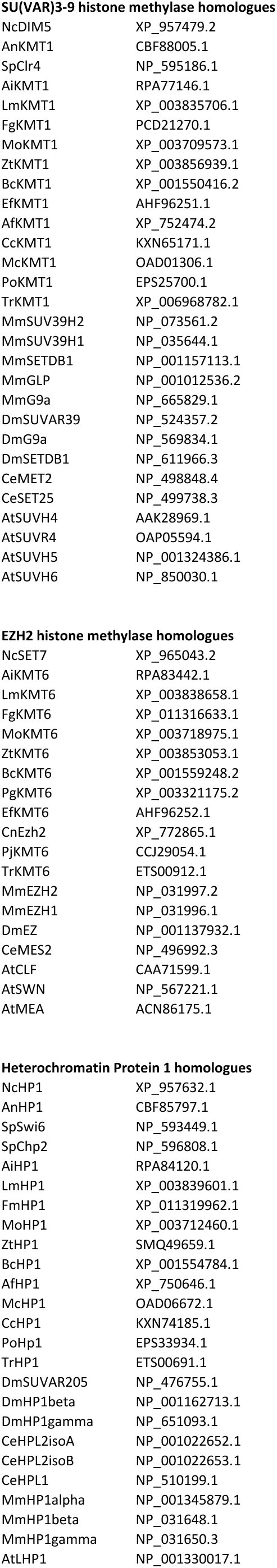
Accession numbers for proteins used in alignments to build the phylogenic trees.

SU(VAR)3-9 histone methylase homologues
|  |  |
| --- | --- |
| NcDIM5 | XP_957479.2 |
| AnKMT1 | CBF88005.1 |
| SpClr4 | NP_595186.1 |
| AiKMT1 | RPA77146.1 |
| LmKMT1 | XP_003835706.1 |
| FgKMT1 | PCD21270.1 |
| MoKMT1 | XP_003709573.1 |
| ZtKMT1 | XP_003856939.1 |
| BcKMT1 | XP_001550416.2 |
| EfKMT1 | AHF96251.1 |
| AfKMT1 | XP_752474.2 |
| CcKMT1 | KXN65171.1 |
| McKMT1 | OAD01306.1 |
| PoKMT1 | EPS25700.1 |
| TrKMT1 | XP_006968782.1 |
| MmSUV39H2 | NP_073561.2 |
| MmSUV39H1 | NP_035644.1 |
| MmSETDB1 | NP_001157113.1 |
| MmGLP | NP_001012536.2 |
| MmG9a | NP_665829.1 |
| DmSUVAR39 | NP_524357.2 |
| DmG9a | NP_569834.1 |
| DmSETDB1 | NP_611966.3 |
| CeMET2 | NP_498848.4 |
| CeSET25 | NP_499738.3 |
| AtSUVH4 | AAK28969.1 |
| AtSUVR4 | OAP05594.1 |
| AtSUVH5 | NP_001324386.1 |
| AtSUVH6 | NP_850030.1 |
| **EZH2 histone methylase homologues** |  |
| NcSET7 | XP_965043.2 |
| AiKMT6 | RPA83442.1 |
| LmKMT6 | XP_003838658.1 |
| FgKMT6 | XP_011316633.1 |
| MoKMT6 | XP_003718975.1 |
| ZtKMT6 | XP_003853053.1 |
| BcKMT6 | XP_001559248.2 |
| PgKMT6 | XP_003321175.2 |
| EfKMT6 | AHF96252.1 |
| CnEzh2 | XP_772865.1 |
| PjKMT6 | CCJ29054.1 |
| TrKMT6 | ETS00912.1 |
| MmEZH2 | NP_031997.2 |
| MmEZH1 | NP_031996.1 |
| DmEZ | NP_001137932.1 |
| CeMES2 | NP_496992.3 |
| AtCLF | CAA71599.1 |
| AtSWN | NP_567221.1 |
| AtMEA | ACN86175.1 |
| **Heterochromatin Protein 1 homologues** |  |
| NcHP1 | XP_957632.1 |
| AnHP1 | CBF85797.1 |
| SpSwi6 | NP_593449.1 |
| SpChp2 | NP_596808.1 |
| AiHP1 | RPA84120.1 |
| LmHP1 | XP_003839601.1 |
| FmHP1 | XP_011319962.1 |
| MoHP1 | XP_003712460.1 |
| ZtHP1 | SMQ49659.1 |
| BcHP1 | XP_001554784.1 |
| AfHP1 | XP_750646.1 |
| McHP1 | OAD06672.1 |
| CcHP1 | KXN74185.1 |
| PoHP1 | EPS33934.1 |
| TrHP1 | ETS00691.1 |
| DmSUVAR205 | NP_476755.1 |
| DmHP1beta | NP_001162713.1 |
| DmHP1gamma | NP_651093.1 |
| CeHPL2isoA | NP_001022652.1 |
| CeHPL2isoB | NP_001022653.1 |
| CeHPL1 | NP_510199.1 |
| MmHP1alpha | NP_001345879.1 |
| MmHP1beta | NP_031648.1 |
| MmHP1gamma | NP_031650.3 |
| AtLHP1 | NP_001330017.1 |

**Table S6.** Heat map of Spearman correlation coefficient comparison between ChIP-seq samples. Spearman’s correlation coefficient confirms the reproducibility of the experiments since biological replicates cluster together with high correlation coefficient. Moreover, H3K9me3 and H3K27me3 IP share a same cluster that negatively correlates with the H3K4me3 cluster. Both display no correlation with non-IP input and negative sample (Mock and H3K27me3 in the Δ*PaKmt6*), which means that IP results are significant and specific.

|  | KMT6-11d_K9 | KMT6-17d_K9 | KMT1-1d_K27 | HP1-4d_K9 | HP1-3d_K9 | S_2e_K9 | S_3d_K9 | KMT1-1d_K9 | KMT1-2d_K9 | HP1-4d_K27 | HP1-3d_K27 | KMT1-2d_K27 | S_2e_K27 | S_3d_K27 | HP1-4d_K4 | KMT6-11d_K4 | KMT1-1d_K4 | KMT6-17d_K4 | S_2e_K4 | S_3d_K4 | HP1-3d_K4 | KMT1-2d_K4 | KMT6-17d_K27 | mock | KMT6-11d_K27 | input |
| --- | --- | --- | --- | --- | --- | --- | --- | --- | --- | --- | --- | --- | --- | --- | --- | --- | --- | --- | --- | --- | --- | --- | --- | --- | --- | --- |
| KMT6-11d_K9 | 1 | 0,834 | 0,378 | 0,617 | 0,731 | 0,656 | 0,651 | 0,66 | 0,672 | 0,575 | 0,592 | 0,553 | 0,491 | 0,469 | -0,165 | -0,21 | -0,253 | -0,249 | -0,047 | -0,041 | 0,051 | -0,006 | -0,069 | 0,138 | 0,395 | 0,294 |
| KMT6-17d_K9 | 0,834 | 1 | 0,315 | 0,48 | 0,555 | 0,51 | 0,523 | 0,569 | 0,576 | 0,481 | 0,486 | 0,455 | 0,397 | 0,392 | -0,029 | -0,078 | -0,088 | -0,077 | 0,122 | 0,126 | 0,183 | 0,142 | 0,111 | 0,173 | 0,495 | 0,316 |
| KMT1-1d_K27 | 0,378 | 0,315 | 1 | 0,767 | 0,667 | 0,745 | 0,736 | 0,663 | 0,688 | 0,717 | 0,697 | 0,708 | 0,73 | 0,726 | 0,005 | -0,035 | -0,024 | -0,021 | -0,281 | -0,288 | -0,148 | -0,229 | 0,161 | 0,415 | 0,324 | 0,013 |
| HP1-4d_K9 | 0,617 | 0,48 | 0,767 | 1 | 0,843 | 0,867 | 0,83 | 0,703 | 0,721 | 0,701 | 0,681 | 0,667 | 0,677 | 0,651 | -0,116 | -0,149 | -0,107 | -0,131 | -0,303 | -0,315 | -0,205 | -0,283 | 0,092 | 0,355 | 0,237 | -0,076 |
| HP1-3d_K9 | 0,731 | 0,555 | 0,667 | 0,843 | 1 | 0,92 | 0,919 | 0,758 | 0,793 | 0,84 | 0,847 | 0,824 | 0,782 | 0,761 | -0,294 | -0,333 | -0,38 | -0,364 | -0,316 | -0,3 | -0,157 | -0,248 | -0,168 | 0,128 | 0,196 | 0,174 |
| S_2e_K9 | 0,656 | 0,51 | 0,745 | 0,867 | 0,92 | 1 | 0,968 | 0,823 | 0,848 | 0,864 | 0,848 | 0,85 | 0,871 | 0,845 | -0,272 | -0,326 | -0,323 | -0,338 | -0,323 | -0,324 | -0,182 | -0,271 | -0,124 | 0,167 | 0,18 | 0,081 |
| S_3d_K9 | 0,651 | 0,523 | 0,736 | 0,83 | 0,919 | 0,968 | 1 | 0,822 | 0,854 | 0,897 | 0,883 | 0,886 | 0,896 | 0,878 | -0,275 | -0,326 | -0,353 | -0,346 | -0,303 | -0,294 | -0,147 | -0,239 | -0,151 | 0,135 | 0,185 | 0,152 |
| KMT1-1d_K9 | 0,66 | 0,569 | 0,663 | 0,703 | 0,758 | 0,823 | 0,822 | 1 | 0,964 | 0,821 | 0,809 | 0,813 | 0,822 | 0,795 | -0,195 | -0,265 | -0,252 | -0,288 | -0,15 | -0,167 | 0,005 | -0,073 | -0,133 | 0,1 | 0,279 | 0,235 |
| KMT1-2d_K9 | 0,672 | 0,576 | 0,688 | 0,721 | 0,793 | 0,848 | 0,854 | 0,964 | 1 | 0,858 | 0,851 | 0,852 | 0,852 | 0,829 | -0,195 | -0,26 | -0,268 | -0,289 | -0,166 | -0,175 | -0,003 | -0,087 | -0,134 | 0,106 | 0,28 | 0,259 |
| HP1-4d_K27 | 0,575 | 0,481 | 0,717 | 0,701 | 0,84 | 0,864 | 0,897 | 0,821 | 0,858 | 1 | 0,941 | 0,945 | 0,916 | 0,911 | -0,266 | -0,315 | -0,379 | -0,348 | -0,255 | -0,236 | -0,063 | -0,156 | -0,198 | 0,041 | 0,202 | 0,306 |
| HP1-3d_K27 | 0,592 | 0,486 | 0,697 | 0,681 | 0,847 | 0,848 | 0,883 | 0,809 | 0,851 | 0,941 | 1 | 0,945 | 0,902 | 0,904 | -0,276 | -0,314 | -0,392 | -0,356 | -0,258 | -0,235 | -0,043 | -0,143 | -0,215 | 0,02 | 0,215 | 0,342 |
| KMT1-2d_K27 | 0,553 | 0,455 | 0,708 | 0,667 | 0,824 | 0,85 | 0,886 | 0,813 | 0,852 | 0,945 | 0,945 | 1 | 0,921 | 0,92 | -0,271 | -0,312 | -0,392 | -0,355 | -0,251 | -0,229 | -0,047 | -0,142 | -0,227 | 0,012 | 0,19 | 0,331 |
| S_2e_K27 | 0,491 | 0,397 | 0,73 | 0,677 | 0,782 | 0,871 | 0,896 | 0,822 | 0,852 | 0,916 | 0,902 | 0,921 | 1 | 0,957 | -0,26 | -0,311 | -0,344 | -0,336 | -0,274 | -0,266 | -0,084 | -0,176 | -0,192 | 0,043 | 0,162 | 0,229 |
| S_3d_K27 | 0,469 | 0,392 | 0,726 | 0,651 | 0,761 | 0,845 | 0,878 | 0,795 | 0,829 | 0,911 | 0,904 | 0,92 | 0,957 | 1 | -0,223 | -0,269 | -0,319 | -0,3 | -0,238 | -0,223 | -0,031 | -0,128 | -0,181 | 0,027 | 0,176 | 0,251 |
| HP1-4d_K4 | -0,165 | -0,029 | 0,005 | -0,116 | -0,294 | -0,272 | -0,275 | -0,195 | -0,195 | -0,266 | -0,276 | -0,271 | -0,26 | -0,223 | 1 | 0,854 | 0,744 | 0,794 | 0,655 | 0,641 | 0,574 | 0,59 | 0,447 | 0,33 | 0,236 | -0,013 |
| KMT6-11d_K4 | -0,21 | -0,078 | -0,035 | -0,149 | -0,333 | -0,326 | -0,326 | -0,265 | -0,26 | -0,315 | -0,314 | -0,312 | -0,311 | -0,269 | 0,854 | 1 | 0,754 | 0,859 | 0,636 | 0,63 | 0,562 | 0,578 | 0,435 | 0,31 | 0,206 | -0,017 |
| KMT1-1d_K4 | -0,253 | -0,088 | -0,024 | -0,107 | -0,38 | -0,323 | -0,353 | -0,252 | -0,268 | -0,379 | -0,392 | -0,392 | -0,344 | -0,319 | 0,744 | 0,754 | 1 | 0,857 | 0,506 | 0,467 | 0,38 | 0,417 | 0,714 | 0,4 | 0,199 | -0,235 |
| KMT6-17d_K4 | -0,249 | -0,077 | -0,021 | -0,131 | -0,364 | -0,338 | -0,346 | -0,288 | -0,289 | -0,348 | -0,356 | -0,355 | -0,336 | -0,3 | 0,794 | 0,859 | 0,857 | 1 | 0,568 | 0,55 | 0,465 | 0,492 | 0,617 | 0,395 | 0,253 | -0,1 |
| S_2e_K4 | -0,047 | 0,122 | -0,281 | -0,303 | -0,316 | -0,323 | -0,303 | -0,15 | -0,166 | -0,255 | -0,258 | -0,251 | -0,274 | -0,238 | 0,655 | 0,636 | 0,506 | 0,568 | 1 | 0,942 | 0,846 | 0,881 | 0,182 | -0,02 | 0,209 | 0,214 |
| S_3d_K4 | -0,041 | 0,126 | -0,288 | -0,315 | -0,3 | -0,324 | -0,294 | -0,167 | -0,175 | -0,236 | -0,235 | -0,229 | -0,266 | -0,223 | 0,641 | 0,63 | 0,467 | 0,55 | 0,942 | 1 | 0,858 | 0,887 | 0,154 | -0,043 | 0,207 | 0,262 |
| HP1-3d_K4 | 0,051 | 0,183 | -0,148 | -0,205 | -0,157 | -0,182 | -0,147 | 0,005 | -0,003 | -0,063 | -0,043 | -0,047 | -0,084 | -0,031 | 0,574 | 0,562 | 0,38 | 0,465 | 0,846 | 0,858 | 1 | 0,898 | 0,079 | -0,079 | 0,256 | 0,35 |
| KMT1-2d_K4 | -0,006 | 0,142 | -0,229 | -0,283 | -0,248 | -0,271 | -0,239 | -0,073 | -0,087 | -0,156 | -0,143 | -0,142 | -0,176 | -0,128 | 0,59 | 0,578 | 0,417 | 0,492 | 0,881 | 0,887 | 0,898 | 1 | 0,102 | -0,086 | 0,234 | 0,321 |
| KMT6-17d_K27 | -0,069 | 0,111 | 0,161 | 0,092 | -0,168 | -0,124 | -0,151 | -0,133 | -0,134 | -0,198 | -0,215 | -0,227 | -0,192 | -0,181 | 0,447 | 0,435 | 0,714 | 0,617 | 0,182 | 0,154 | 0,079 | 0,102 | 1 | 0,554 | 0,37 | -0,208 |
| mock | 0,138 | 0,173 | 0,415 | 0,355 | 0,128 | 0,167 | 0,135 | 0,1 | 0,106 | 0,041 | 0,02 | 0,012 | 0,043 | 0,027 | 0,33 | 0,31 | 0,4 | 0,395 | -0,02 | -0,043 | -0,079 | -0,086 | 0,554 | 1 | 0,449 | -0,149 |
| KMT6-11d_K27 | 0,395 | 0,495 | 0,324 | 0,237 | 0,196 | 0,18 | 0,185 | 0,279 | 0,28 | 0,202 | 0,215 | 0,19 | 0,162 | 0,176 | 0,236 | 0,206 | 0,199 | 0,253 | 0,209 | 0,207 | 0,256 | 0,234 | 0,37 | 0,449 | 1 | 0,268 |
| input | 0,294 | 0,316 | 0,013 | -0,076 | 0,174 | 0,081 | 0,152 | 0,235 | 0,259 | 0,306 | 0,342 | 0,331 | 0,229 | 0,251 | -0,013 | -0,017 | -0,235 | -0,1 | 0,214 | 0,262 | 0,35 | 0,321 | -0,208 | -0,149 | 0,268 | 1 |

**Table S7.** Raw data from the heatmap of Spearman’s correlation coefficient comparison in Fig. S14.

| | $\Delta PaKmt6\_H3K9me3$ | $\Delta PaHP1\_H3K9me3$ | WT_H3K9me3 | $\Delta PaKmt1\_H3K27me3$ | $\Delta PaKmt1\_H3K9me3$ | $\Delta PaHP1\_H3K27me3$ | WT_H3K27me3 | $\Delta PaKmt1\_H3K4me3$ | $\Delta PaKmt6\_H3K4me3$ | $\Delta PaHP1\_H3K4me3$ | WT_H3K4me3 | input | $\Delta PaKmt6\_H3K27me3$ | mock |
| --- | --- | --- | --- | --- | --- | --- | --- | --- | --- | --- | --- | --- | --- | --- |
| $\Delta PaKmt6\_H3K9me3$ | 1.00 | 0.65 | 0.62 | 0.49 | 0.65 | 0.57 | 0.46 | -0.11 | -0.18 | 0.04 | 0.04 | 0.31 | 0.21 | 0.16 |
| $\Delta PaHP1\_H3K9me3$ | 0.65 | 1.00 | 0.92 | 0.82 | 0.78 | 0.80 | 0.75 | -0.31 | -0.26 | -0.21 | -0.32 | 0.05 | 0.08 | 0.26 |
| WT_H3K9me3 | 0.62 | 0.92 | 1.00 | 0.86 | 0.85 | 0.89 | 0.88 | -0.38 | -0.36 | -0.23 | -0.31 | 0.12 | -0.02 | 0.15 |
| $\Delta PaKmt1\_H3K27me3$ | 0.49 | 0.82 | 0.86 | 1.00 | 0.80 | 0.89 | 0.88 | -0.26 | -0.21 | -0.12 | -0.27 | 0.21 | 0.09 | 0.22 |
| $\Delta PaKmt1\_H3K9me3$ | 0.65 | 0.78 | 0.85 | 0.80 | 1.00 | 0.85 | 0.84 | -0.22 | -0.29 | -0.08 | -0.16 | 0.25 | 0.03 | 0.10 |
| $\Delta PaHP1\_H3K27me3$ | 0.57 | 0.80 | 0.89 | 0.89 | 0.85 | 1.00 | 0.93 | -0.36 | -0.35 | -0.16 | -0.24 | 0.33 | -0.06 | 0.03 |
| WT_H3K27me3 | 0.46 | 0.75 | 0.88 | 0.88 | 0.84 | 0.93 | 1.00 | -0.32 | -0.32 | -0.15 | -0.25 | 0.25 | -0.06 | 0.03 |
| $\Delta PaKmt1\_H3K4me3$ | -0.11 | -0.31 | -0.38 | -0.26 | -0.22 | -0.36 | -0.32 | 1.00 | 0.84 | 0.77 | 0.74 | -0.02 | 0.51 | 0.23 |
| $\Delta PaKmt6\_H3K4me3$ | -0.18 | -0.26 | -0.36 | -0.21 | -0.29 | -0.35 | -0.32 | 0.84 | 1.00 | 0.71 | 0.62 | -0.07 | 0.49 | 0.36 |
| $\Delta PaHP1\_H3K4me3$ | 0.04 | -0.21 | -0.23 | -0.12 | -0.08 | -0.16 | -0.15 | 0.77 | 0.71 | 1.00 | 0.86 | 0.24 | 0.29 | 0.10 |
| WT_H3K4me3 | 0.04 | -0.32 | -0.31 | -0.27 | -0.16 | -0.24 | -0.25 | 0.74 | 0.62 | 0.86 | 1.00 | 0.25 | 0.21 | -0.03 |
| input | 0.31 | 0.05 | 0.12 | 0.21 | 0.25 | 0.33 | 0.25 | -0.02 | -0.07 | 0.24 | 0.25 | 1.00 | -0.05 | -0.15 |
| $\Delta PaKmt6\_H3K27me3$ | 0.21 | 0.08 | -0.02 | 0.09 | 0.03 | -0.06 | -0.06 | 0.51 | 0.49 | 0.29 | 0.21 | -0.05 | 1.00 | 0.60 |
| mock | 0.16 | 0.26 | 0.15 | 0.22 | 0.10 | 0.03 | 0.03 | 0.23 | 0.36 | 0.10 | -0.03 | -0.15 | 0.60 | 1.00 |

